# Teleport-Stabilized Quantum-Walk Transport for Robust Neoantigen Ranking in Near-Tie Regimes

**DOI:** 10.64898/2026.04.22.720240

**Authors:** Ioannis Grigoriadis, Christos Emmanouilides

## Abstract

Personalized neoantigen vaccination is a patient-specific decision problem: given a tumor’s molecular signature—somatic mutations, clonality, RNA expression, and antigen-processing context—we must choose a small, manufacturable peptide set that stays therapeutically relevant under uncertainty. In late-stage pipelines, candidates often collapse into near-ties: binding/presentation estimates, immunogenicity surrogates, and structure-based refinement compress many peptides into narrow score bands, making the final top-K fragile to small shifts in calibration, scaling, sampling, or docking protocols. Similar instability arises in peptide–target discovery when multiple hypotheses remain comparably supported. We introduce a transport-stabilized ranking layer that prioritizes redundancy structure over marginal score differences. Peptides (and structural microstates) become nodes in a patient-conditioned evidence graph; edges encode evidence overlap (motifs/HLA restrictions, processing features, target neighborhoods, pocket/contact fingerprints). We apply symmetry-aware quotient reduction of a normalized graph operator, collapsing near-symmetric neighborhoods into basin units while preserving effective shortlist couplings. Discriminative basin fingerprints are then extracted using coherent quantum-walk transport, ∣*ψ*(*t*)⟩ = *e*^―*iHt*^∣*ψ*(0)⟩, with visitation *P*(*v*,*t*) = ∣⟨*v*∣*ψ*(*t*)⟩∣^2^. Because coherent dynamics are oscillatory and horizon-dependent, we introduce a teleport-consensus channel that mixes unitary transport with restart to yield a stationary marginal suitable for stable ranking, *ρ*_*t*+1_ = (1 ― *α*)*Uρ*_*t*_*U*^†^ + *α* Σ_*j*_*v*_*j*_∣*j*⟩⟨*j*∣, and *π*_*i*_ = Tr(Π_*i*_*ρ*). Information-theoretic polygraphs—entropy, dispersion, and consensus traces—quantify stabilization and provide an interpretable tie-breaking audit trail. We demonstrate consistent stabilization across colorectal-cancer contexts spanning peptide–target mechanistic triage, microstate symmetry auditing, multimodal evidence fusion, docking-ensemble geometrization, and patient-specific neoantigen shortlist construction.

## 1. Introduction

Selecting a manufacturable neoantigen set for a personalized cancer vaccine is a constrained, high-leverage decision: per patient, thousands of mutant peptide candidates are enumerated, but only a small top-K can be synthesized and administered. This bottleneck induces a characteristic late-stage regime: after aggressive filtering, the remaining candidates form an enriched band of simultaneously plausible peptides. The shortlist therefore often enters a near-tie regime in which binding/presentation, processing, immunogenicity surrogates, and structural refinement compress candidates into narrow score bands, so top-K membership becomes sensitive to routine perturbations (model updates, calibration, feature scaling, sampling, docking protocol choices, and correlated evidence reuse) despite unchanged biological interpretation. In pattern-recognition terms, the objective shifts from maximizing a single score to engineering a decision rule that is stable to nuisance variation, exposes invariances/symmetries, and aggregates correlated evidence in an auditable manner.

This review treats shortlist instability as a property of **evidence geometry** rather than a defect of any single predictor. Candidates are redundancy-rich (overlapping k-mers, shared motifs, shared HLA restrictions, shared mutation/RNA support, and—under docking/refinement—shared microstate narratives), so ranking is better posed as comparing **neighborhoods** and **basins** of support. We encode evidence as a weighted affinity graph 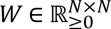 with degrees *D* = diag(*W***1**) and operator backbone

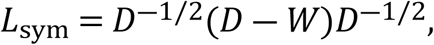

which unifies diffusion baselines, spectral coarse-graining, and coherent transport on graphs. The core design question is then operator/readout design: how to represent redundancy and near-symmetry explicitly, and how to extract a reproducible ranking marginal from dynamics that reveal competition among similar hypotheses. Restart/teleport mixing supplies the stabilizer.

A personalized vaccine setting makes the stakes explicit: the final shortlist is a manufacturing action under uncertainty. The same near-tie geometry also appears in CRC nutraceutical peptide–target inference, motivating a single stability-first pattern-recognition frame across both domains.

### 1.1 Near ties as a redundancy + symmetry phenomenon

Near ties are an expected output of conservative screening: one mutation yields multiple overlapping k-mers; one allele admits multiple binders; multiple alleles yield portfolios clustered in a high-confidence band. Evidence channels are coupled (shared locus-level support; predictor inductive biases), producing redundancy where small perturbations can reorder near-interchangeable candidates. A disciplined method should therefore be invariant to nuisance transformations and should summarize what truly distinguishes candidates.

#### Graph neighborhoods and measurable tie risk

We formalize redundancy as overlap neighborhoods: edges represent shared provenance, HLA restriction/presentation, processing support, immunogenicity signals, pathway/target neighborhood similarity, or microstate/contact consistency. Dense near-symmetric clusters imply brittle micro-ordering; separated basins imply intrinsically more stable ordering.

#### Interchangeability as approximate symmetry and quotient reduction

Near ties correspond to approximate symmetries: permutations within a redundancy basin change little about the evidence graph. Given a partition matrix **S** encoding basin blocks, we coarse-grain via the quotient operator

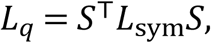

shifting the decision unit from individual micro-competitors to basin-level representatives.

#### Transport as a basin probe: diffusion versus coherent quantum walks

Diffusion provides a monotone baseline,

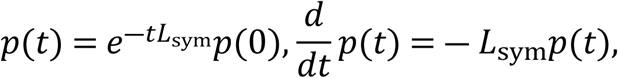

while coherent transport probes basin competition via CTQW

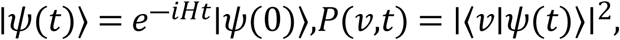

(or coined DTQW ∣*ψ*_*t*+1_⟩ = *U*∣*ψ*_*t*_⟩, *U* = **S**(*C*(*θ*) ⊗ *I*), with marginals *P*(*x*,*t*) = Σ_*c*_∣⟨*x*,*c*∣*ψ*_*t*_⟩∣^2^). Coherent-versus-diffusive contrast is used as an interpretability control: shared-basin candidates show similar spreading; competing basins yield sharper, oscillatory dominance patterns.

#### Polygraph diagnostics

We audit tie risk and stabilization using entropy and dispersion:

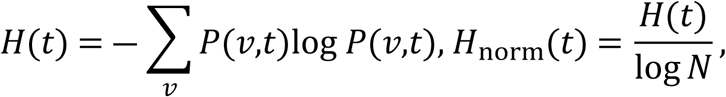

with companion spread/dispersion traces and coherent–classical divergence Δ*P*(*v*,*t*) = *P*_*Q*_(*v*,*t*) ― *P*_*C*_(*v*,*t*) when needed.

#### Teleport-consensus stabilization

Because coherent dynamics are non-convergent, we convert oscillatory evidence into a stationary decision marginal. Classical restart ranking uses

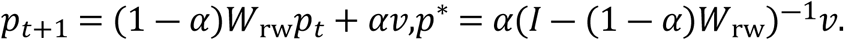

For coherent evidence we use a restart-mixed open-system readout

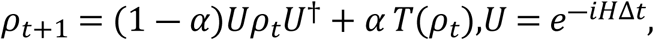

with practical consensus reset *T*(*ρ*) = Σ_*j*_*v*_*j*_∣*j*⟩⟨*j*∣ and marginals *π*_*i*_(*t*) = Tr(Π_*i*_*ρ*_*t*_), Π_*i*_ = ∣*i*⟩⟨*i*∣. Operationally, *α* is treated as an explicit controller parameter (often *α* ≈ 0.18) rather than a hidden tuning trick.

### 1.2 Contributions and why they matter

i. **Unified evidence → graph → operator → transport → readout spine.** Heterogeneous evidence is mapped to *W*, summarized by *L*_sym_, optionally coarse-grained by *L*_*q*_, probed by diffusion/quantum transport, and stabilized by teleport-consensus to yield a reproducible ranking marginal.
ii. **Symmetry-aware basin ranking for near ties.** Quotient reduction converts brittle micro-ordering into basin-level decision objects aligned with manufacturable top-K selection.
iii. **Auditable coherent-versus-diffusive diagnostics.** Intensity maps plus entropy/spread polygraphs quantify whether dominance reflects a reproducible basin (decision-stable) or unresolved competition (tie-risk).
iv. **Teleport-consensus as decision readout.** Restart mixing turns horizon-dependent coherent evidence into a stationary, comparable marginal suitable for governance of late-stage shortlist decisions.
v. **Compatibility with learned representations.** Upstream predictors or learned graph affinities can supply *W* and *v*; the proposed layer acts as a post-representation stabilizer for the compressed, redundancy-rich band where “best” must mean best under perturbations, not merely highest under a single scoring snapshot.

## 2. Related work

### 2.1 Pattern recognition and stability-oriented ranking

Ranking on data manifolds, diffusion maps, and local/global consistency establish the value of graph structure for stable propagation and retrieval [8,34,35]. Robust rank aggregation addresses instability when multiple weakly consistent lists must be fused [9]. Calibration and uncertainty are essential when decisions depend on narrow score gaps [10–12].

### 2.2 Spectral graph operators, clustering, and quotient structure

Normalized Laplacians and spectral embeddings provide principled geometry for graphs and clustering [4,5]. Community structure and modularity have long been central for mechanistic interpretation in biological networks [3,32]. Signal processing on graphs formalizes how features propagate under graph operators [33].

### 2.3 Teleportation/restart as convergence control

PageRank and its generalizations operationalize restart/teleportation to guarantee convergence and reduce sensitivity to traps and disconnected components [6,7]. The teleport-consensus channel used here is a restart-mixed readout in the same conceptual family, adapted to coherent transport outputs (SI Appendix I, Figs. 2h, 8a).

### 2.4 Quantum walks as discriminative transport probes

Quantum walks provide interference-driven transport signatures distinct from diffusion [13–16]. They can yield faster early spreading, oscillatory basin competition, and transient amplification useful for tie-breaking—but also introduce horizon dependence (explicitly visualized in the coherent intensity maps and entropy polygraphs; Figs. 2e,g; 9m).

### 2.5 Neoantigen prediction context

Neoantigen vaccination and prioritization depend on antigen processing/presentation and evidence fusion across multiple channels [19–28]. These advances improve recall but can increase near-tie frequency in final shortlists, strengthening the need for stability-oriented decision layers.

## 3. Method

### 3.1 Candidate/microstate overlap graph

Let 𝒫 = {*p*_1_,…,*p*_*N*_} denote **patient-conditioned peptide–HLA candidates** and/or **structure-derived microstates** (e.g., docking poses, refined pocket-contact states). We represent the late-stage shortlist as a **redundancy-rich hypothesis set** and encode it as a weighted undirected graph *G* = (*V*,*E*,*W*) with *V* = {1,…,*N*}. The affinity matrix *W* ∈ 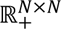 quantifies **overlap/compatibility** between candidates, so that near ties appear as **dense, near-symmetric neighborhoods** rather than arbitrary score gaps (SI Appendix I, Figs. 5–6; docking-microstate abstractions in Fig. 1H–K and Fig. 9d–h).

Concretely, we construct entries as a convex fusion of evidence kernels,

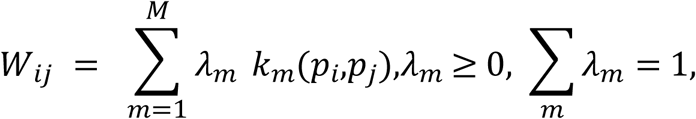

where *k*_*m*_ may include (i) **sequence/motif overlap** and shared HLA restriction, (ii) **co-support under patient molecular signature** (mutation provenance, expression/clonality, processing surrogates), and (iii) **geometric/contact consistency** for microstates (pocket-contact fingerprints, interaction-pattern agreement) consistent with the pocket-geometrization overlays used later for interpretability (SI Appendix I, Fig. 8a; Fig. 9d–h). The graph degree statistics are

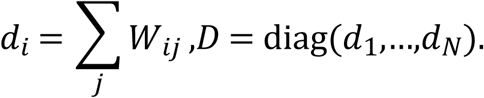

This representation makes the selection problem explicitly **relational**: candidates are ranked not only by marginal scores but by how robustly they sit inside (or bridge) **evidence basins**—the precise objects that dominate stability in near-tie regimes.

### 3.2 Symmetry-normalized Laplacian and quotient reduction

We adopt the symmetric normalized Laplacian [4,5] as the operator backbone,

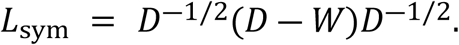

This choice is standard in spectral graph theory and provides a single language for diffusion baselines, coherent transport generators, and block-level coarse-graining.

Near ties typically arise because many nodes are **approximately interchangeable** (shared mutation provenance, overlapping *k*-mers, similar presentation predictions, or similar microstate contacts). We therefore encode a *K*-block partition with an assignment matrix **S** ∈ {0,1}^*N*×*K*^(one-hot rows), and collapse near-symmetric neighborhoods using the quotient operator

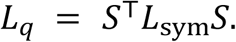

This step corresponds directly to the “symmetry-reduced (cluster/quotient) operator” depicted as **S**^⊤^*L*_sym_**S**(SI Appendix I, Fig. 2c; Fig. 4d; Fig. 5a–d; Fig. 6a–b). Operationally, the quotient reduction shifts the decision unit from brittle micro-competitors to **basin-level blocks** whose diagonal dominance reflects within-basin cohesion, while off-diagonals quantify residual inter-basin coupling—exactly the structure that governs stable top-*K* selection under redundancy.

### 3.3 Geometry corroboration via fidelity kernels

To corroborate that the induced geometry is consistent with the quotient/block structure, we compute an FS-style fidelity snapshot,

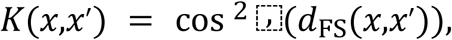

yielding block-structured similarity maps aligned with the inferred clustering (SI Appendix I, Fig. 2d; Fig. 4f–g; Fig. 5a–d). In the numeric readout example, kernel values fall in the ∼ 0.85–1.00 range with mean ∼ 0.89(SI Appendix I, Fig. 4f), indicating a **high-similarity, near-tie manifold** that is nonetheless **separable into basins** (SI Appendix I, Fig. 4g). This agreement check is used as an audit: stable ranking is expected only when multiple, independent summaries (operator blocks + geometric similarity) support the same basin structure.

### 3.4 Classical baseline: diffusion / random walk with restart

As a convergent comparator grounded in classical ranking theory [6–8,34,35], we use diffusion and random-walk-with-restart (RWR) propagation on the graph:

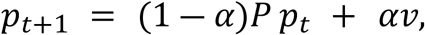

where *P* is a row-stochastic transition derived from *W* (e.g., *P* = *D*^―1^*W*), and *v* is a personalization prior (uniform or evidence-weighted). This baseline produces a stationary distribution and provides the classical reference for all coherent-vs-diffusive comparisons (SI Appendix I, Figs. 2f, 6f, 7f, 9l–m), separating “stability by monotone smoothing” from “stability by redundancy-aware basin discrimination.”

### 3.5 Coherent transport: continuous-time and coined discrete-time quantum walks

To probe basin structure in compressed score bands, we use coherent transport on graph-derived generators [13–16]. In continuous time, a state evolves under a Hermitian generator *H* (commonly Laplacian- or adjacency-derived),

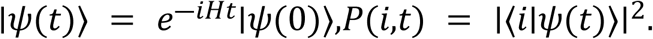

When the generator is built from affinities and degrees, a canonical choice is

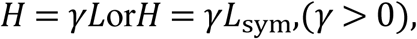

and after quotienting we analogously use *H*_*q*_ = *γL*_*q*_. These dynamics are rendered as node-by-time intensity maps that expose localization bands and oscillatory basin competition (SI Appendix I, Fig. 2e; Fig. 4b; Fig. 6e; Fig. 7e; Fig. 9l; benchmark contrast in Fig. 9m).

For controller-style implementations consistent with (SI Appendix I, Fig. 8a), we also use a coined discrete-time quantum walk (DTQW):

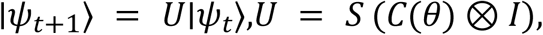

with a position marginal

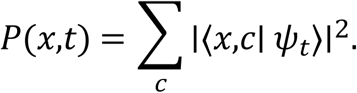

The DTQW view makes explicit how interference patterns (“light-cone” structure) provide a discriminative fingerprint of basin membership beyond diffusion, and matches the controller schematic used to couple transport, teleportation, and consensus readout (SI Appendix I, Fig. 8a).

### 3.6 Teleport-consensus stabilization: restart-mixed open-system readout with teleportation handoff

Coherent quantum-walk marginals are informative but **horizon-dependent** and generally **non-convergent**, so a decision-grade ranking requires an explicit stabilization channel. We therefore use a convex mixture update in density-operator form (SI Appendix I, Fig. 8a):

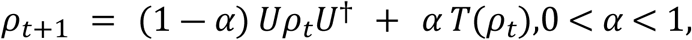

with *U* = *e*^―*iH*Δ*t*^ for CTQW or *U* as the DTQW step operator. A practical consensus reset is

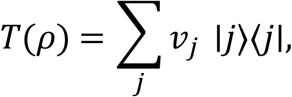

and measured marginals are

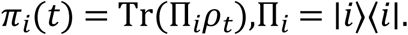

This **teleport-consensus mixing** converts oscillatory coherent evidence into a reproducible, restart-stabilized marginal suitable for top-*K* selection, and is repeatedly annotated with a reusable operating point *α* ≈ 0.18 (SI Appendix I, Fig. 2h; controller schematic in Fig. 8a; per-row fixed *α* = 0.18 in Fig. 9a–c).

To connect the “teleport” language to the controller in SI Appendix I, Fig. 8a) (and to separate it from PageRank-style “teleportation” as a restart metaphor), we note that the schematic includes an explicit state-transfer subroutine using an EPR resource ∣Φ^+^⟩ and Bell measurement identity,

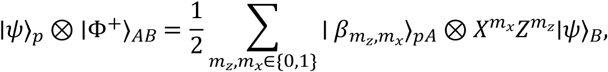

with conditional Pauli feedforward yielding 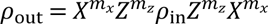. In the manuscript’s decision layer, this handoff serves as a **controller-level mechanism** for moving payload state between registers, while the convex mixing equation above is the **ranking-layer stabilization** that produces stationary, audit-ready marginals in near-tie regimes (SI Appendix I, Fig. 8a–b).

### 3.7 Algorithmic realization: walk assembly and stabilized readout

To make the stabilization procedure reproducible and implementation-facing, we express the workflow as two algorithms aligned with the figure-instantiated pipeline.

#### Algorithm 1

**Structure- and quantum-ensemble walk assembly** *Input:* candidate set *P*, affinities *W*, clustering map **S**, horizon *T*. *Output:* quotient operator *L*_*q*_, transport diagnostics, basin representatives (SI Appendix I Fig. 2c,g) [19–22].

1. Build affinity graph *W* from multi-evidence overlap; compute degrees *D*.
2. Form the normalized operator *L*_sym_ = *D*^―1/2^(*D* ― *W*)*D*^―1/2^.
3. Perform symmetry/cluster audit; encode blocks by **S**; compute quotient

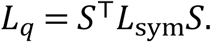
4. Choose transport generator *H* = *γL*_sym_ (full) or *H* = *γL*_*q*_ (quotient).
5. Simulate coherent transport (CTQW, or DTQW if using coined steps) and report polygraphs:

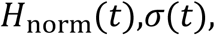 together with basin signatures (localization/competition patterns).

This algorithm instantiates the *probe* stage: it builds the operator backbone and produces the diagnostics indicating whether near ties correspond to interchangeability within a basin or unresolved multi-basin competition.

#### Algorithm 2

**Teleport-consensus quantum walk readout**

*Input:* *U* = *e*^―*iH*Δ*t*^ (or DTQW step), teleport rate *α*, prior *v*.

*Output:* stable ranking *π*^tel^ (SI Appendix I Fig. 2h) [2,3,20].

1. Initialize *ρ*_0_ from the prior *v* (e.g., *ρ*_0_ = Σ_*j*_*v*_*j*_∣*j*⟩⟨*j*∣).
2. Iterate the restart-mixed channel:

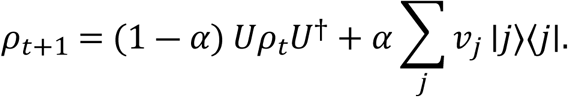
3. Output stable marginals

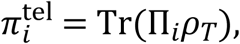

and rank candidates by *π*^tel^ (or feed 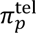 into STS(*p*)).

Algorithm 2 is the *decision* stage: it converts coherent, horizon-sensitive transport evidence into a stationary-style marginal that can be archived, compared across perturbations, and used as a stable tie-breaker. The role of *α* is explicit: it controls the tradeoff between preserving coherent basin structure (small *α*) and enforcing consensus stability (larger *α*), and the manuscript highlights a reusable regime *α* ≈ 0.18 as a practical operating point in the figure-linked contexts.

### 3.8 Interpretability consequence: “basin dominance” rather than “terminal spikes”

The practical effect of teleport-consensus stabilization is that ranking is no longer driven by transient interference peaks that can invert under small horizon shifts. Instead, it is driven by **restart-weighted basin dominance**: candidates that sit in structurally central, redundancy-supported basins accumulate stable probability mass under repeated coherent propagation and controlled reset. This is precisely the desired behavior in near-tie regimes: the readout prefers candidates whose support is *reproducible under perturbations* and *explainable in graph terms* (neighborhood overlap, block membership, and stabilized flow), rather than candidates that win only under a particular time slice of an oscillatory trajectory.

In summary, the teleport-consensus channel operationalizes a pattern-recognition principle for late-stage shortlist decisions: when the evidence geometry is redundancy-rich and near-symmetric, **stable ranking must be engineered at the operator-and-readout level**. The controller schematic’s entanglement-based teleportation handoff clarifies how state routing is realized in the computation layer (SI Appendix I, Fig. 8a), while the restart-mixed density evolution provides the ranking-grade stabilization that yields the auditable marginals used downstream (SI Appendix I, Fig. 8a–b).

### 3.9 Polygraph diagnostics (entropy + spread + consensus)

To make stabilization auditable, we report information-theoretic and dispersion traces (“polygraphs”). For a distribution *P* (⋅,*t*) (classical) or *π* (⋅,*t*) (teleported quantum marginal), Shannon entropy [17] is

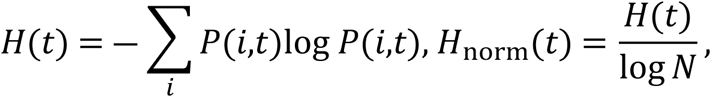

paired with a dispersion proxy (index-based or embedding-based),

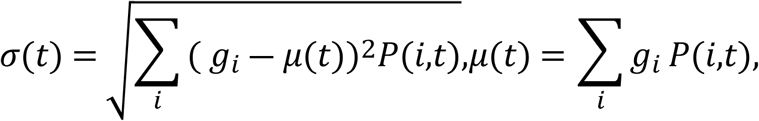

where *g*_*i*_ is a scalar coordinate (node index, diffusion coordinate, or embedding axis) consistent with the plotted geometry. Normalized curves *H*_norm_ and *σ*_norm_ are reported as polygraphs (SI Appendix I, Fig. 2g; Fig. 3b; Fig. 4a–b; Fig. 9a–c). In the MWALK target example, *H*_norm_ and spread rise sharply and saturate near 1.0 within ∼ 5–10 steps, while the consensus marginal plateaus around ∼ 0.65–0.75 (SI Appendix I, Fig. 3b), indicating rapid mixing into a stable basin-level decision distribution.

### 3.10 Grover-style cluster prioritization (optional ranking primitive)

Where basin-level prioritization benefits from explicit focusing, we incorporate a Grover-style amplitude-amplification diagnostic consistent with the controller block in SI Appendix I, Fig. 8a). A simple marking predicate can be defined on edges or transitions, e.g.,

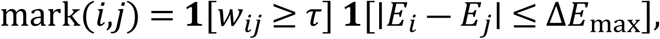

leading to an oracle *0*_mark_ and amplification operator

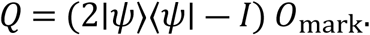

The marked-subspace success trace follows the characteristic oscillation [13–16],

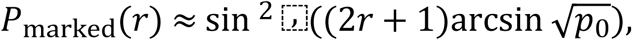

as visualized in the clustered Grover-style trace (SI Appendix I, Fig. 2k) and in the controller schematic (SI Appendix I, Fig. 6h; Fig. 8b). In our use, this block functions as an **optional focusing primitive** and an audit signal—not as a claim of quantum computational advantage—highlighting when “marked” basin transitions dominate the near-tie manifold.

### 3.11 Composite decision statistic for near-tie ranking

Finally, we fuse classical predictors with stabilized transport dominance into a monotone selection statistic designed for manufacturable top-*K* decisions:

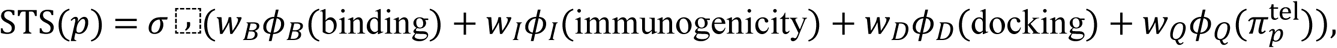

where 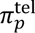 is the teleport-stabilized marginal from Section 3.6, and *σ*(⋅) is a logistic/monotone squashing function for comparability across cases. This matches the explicit composite-score narrative (SI Appendix I, Fig. 9o) and the neoantigen illustration where classical vs quantum-adjusted feature weights are contrasted (SI Appendix I, Fig. 8b). The rationale aligns with Fig. 8b’s message: in compressed near-tie bands, transport-regularized basin dominance provides an **orthogonal stability axis** that can re-rank candidates participating in the most coherent consensus basin, even when classical scores are tightly clustered and sensitive to routine perturbations.

## 4. Experimental protocol

### 4.1 Evidence contexts and figure-linked datasets

We evaluated the proposed transport-stabilized ranking layer across multiple evidence contexts, each explicitly grounded in the provided figure set to ensure that every operator construction, transport diagnostic, and ranking readout is traceable to a concrete dataset rather than to an abstract claim—an evaluation posture aligned with disciplined pattern-recognition practice in near-tie decision regimes, where stability and auditability are primary objectives [1,2]. Across all contexts, the same methodological spine was followed: heterogeneous evidence was summarized into pairwise affinities defining a weighted graph *W*, mapped to an operator via the symmetric normalized Laplacian *L*_sym_ = *D*^―1/2^(*D* ― *W*)*D*^―1/2^ [4,5], optionally compressed by a symmetry-aware quotient **S**^⊤^*L*_sym_**S** when approximate interchangeability was detected [4,5], and then interrogated by coherent quantum-walk transport contrasted against diffusion baselines [13–16]. Because coherent transport is generically non-convergent, a teleport-consensus stabilization channel was used to obtain a reproducible marginal distribution suitable for ranking, consistent in spirit with restart principles in classical ranking systems [6,7]. Information-theoretic polygraphs summarized mixing and stabilization using Shannon entropy and dispersion-related traces [17].

#### 4.1.1 Walnut peptides and colorectal-cancer network pharmacology with quantum pocket overlays

The first evidence context targets **walnut-derived anti-CRC peptide mechanistic inference** under redundancy-rich pathway structure. **SI** **Appendix I**, **Fig. 1A–K** instantiates the dataset end-to-end: GO/KEGG enrichment, pathway-family bubble enrichment, a pathway–gene bipartite network, peptide–target neighborhoods, and a topology-filtered PPI core with hub ranking (e.g., **CASP3** and **MMP9** highlighted). The network-pharmacology component reflects systematic **target/pathway reuse**—a small set of high-connectivity genes can support many enriched pathways—so mechanistic claims become brittle if driven by marginal score differences alone [3,32,33]. To stabilize interpretation in the resulting near-tie regime, the pocket-overlay component encodes docking-derived microstates as contact/overlap graphs and contrasts coherent quantum-walk signatures against diffusion-like baselines (SI Appendix I, Fig. 1H–K), exploiting the diagnostic separation between coherent localization/oscillation and monotone smoothing on graphs [13–16].

#### 4.1.2 Pocket microstate symmetry auditing and teleport-consensus stabilization

The second context explicitly instantiates **symmetry auditing → quotient reduction → coherent vs diffusion transport → stabilized readout** across three representative pocket cases (SI Appendix I, Fig. 2). An affinity graph *W* is built from docking/contact statistics (SI Appendix I, Fig. 2a–b), mapped to *L*_sym_, and compressed via the quotient *L*_*q*_ = **S**^⊤^*L*_sym_**S** to collapse near-symmetric alternatives into basin-level blocks (SI Appendix I, Fig. 2c), consistent with spectral operator foundations [4,5]. Geometry agreement is audited with a cluster-ordered fidelity kernel (SI Appendix I, Fig. 2d). Coherent transport is evaluated by CTQW intensity maps and contrasted against a matched diffusion baseline (SI Appendix I, Fig. 2e–f) [13–16]. Decision stability is enforced with teleport-consensus mixing, yielding a reproducible marginal annotated as **“p =** *α* **≈ 0.18”** (SI Appendix I, Fig. 2h), aligned with restart-based stationary ranking principles [6,7]. Mixing/stabilization are summarized via entropy/spread polygraphs (SI Appendix I, Fig. 2g) grounded in Shannon entropy [17]. The case definitions include explicit docking-energy quotations used as evidence anchors—e.g., Case 1 “Δ*G* = ―11.93 kcal/mol,” and Case 3 “Palmatine Δ*G* = ―33.65,” “R12-8-44-3 Δ*G* = ―33.09,” “Ra 08-2790 (Rec) Δ*G* = ―29.86 kcal/mol” (SI Appendix I, Fig. 2a; SI Appendices I&II)—motivating stabilization even when scores are strong, consistent with known near-tie sensitivity in docking/scoring under protocol variation [18].

#### 4.1.3 MWALK colorectal target evidence fusion on PDB 2X7F

The third context evaluates **heterogeneous evidence fusion** for a colorectal target system using the **PDB 2X7F**composite (SI Appendix I, Fig. 3). Upstream QSAR validation views, structural/interaction depictions, and MD-style readouts define the statistics used to construct *W* (SI Appendix I, Fig. 3a). Transport diagnostics test whether fused evidence forms a **dominant basin** or remains fragmented: entropy/spread/consensus traces provide stabilization readouts (SI Appendix I, Fig. 3b), while geometric embeddings visualize basin compactness/separability (SI Appendix I, Fig. 3c). The polygraphs use Shannon entropy as a principled uncertainty/mixing summary [17], and the motivation follows calibration/uncertainty concerns in multi-evidence pipelines where correlated signals and minor rescaling can induce near ties [10–12]. Coherent behavior is interpreted via established quantum-walk vs diffusion differences on graph operators [13–16].

#### 4.1.4 Multi-cohort and radiogenomic module transport with PASS/teleport logs

The fourth context assesses robustness under **cohort heterogeneity** and **module-structured graphs** using the multi-panel set in SI Appendix I, Fig. 4a–d. Fig. 4a instantiates repeated coined DTQW blocks (**“Repeat ×40”**) with embedded teleport + consensus modules and explicit run-level parameter annotations beside terminal distributions. (SI Appendix I, Fig. 4b) expands diagnostics by juxtaposing DTQW vs CTQW intensity maps and reporting spectral stability traces together with a PASS-oracle plus teleport event log, providing an auditable record of restart-like redistribution. (SI Appendix I, Fig. 4c–d) instantiates symmetry-audited transport for radiogenomic module inference, where integrated radiomic/genetic interaction networks are reduced by quotient operators and evaluated under symmetry-aware transport. The framing follows redundancy/community structure analysis in biological networks [3,32,33], operator construction and quotient coarse-graining in spectral theory [4,5], and restart stabilization in PageRank-style stationary ranking [6,7], while coherent transport follows standard quantum-walk foundations [13–16].

#### 4.1.5 Graph-layer geometrization panels across contexts

To verify that the operator-and-geometry audit is reusable, we analyze the repeated “graph-layer geometrization” panels spanning (**SI** **Appendix I**, **Fig. 4f–g**; **Fig. 5a–d**; **Fig. 6a–b)**. Across these datasets, clustered topology graphs *W* are visualized; quotient heatmaps **S**^⊤^*L*_sym_**S** show diagonal dominance with signed off-diagonal couplings; and FS-style fidelity kernels show blockwise saturation consistent with cluster ordering. These panels function as a standardized audit template: diagonal dominance indicates strong within-basin cohesion under symmetry reduction, while kernel saturation corroborates that similarity geometry matches the operator-domain blocks. The motivation follows spectral/manifold views where Laplacian operators encode geometry and diffusion structure [4,5,34], and transport overlays contrast coherent localization with diffusion smoothing to separate basin signatures from generic denoising [13–16].

#### 4.1.6 Targeting-peptide multimodal validation mapped into transport diagnostics

(SI Appendix I, Fig. 7) provides a multimodal validation context mapped into the same **evidence → graph → operator → transport → readout** spine. Upstream context panels supply the feature/statistics source; each condition/sample is treated as a node (or microstate ensemble), and edges encode overlap in the derived evidence representation. This context demonstrates that the transport-stabilized layer is not restricted to docking microstates: any heterogeneous evidence stack that induces redundancy neighborhoods and near ties can be stabilized under the same graph-transport logic [3,33]. Coherent-vs-diffusion contrasts and entropy-based diagnostics follow the same quantum-walk and information-theoretic principles [13–17].

#### 4.1.7 Neoantigen shortlist illustration with MWALK consensus basin

(SI Appendix I, Fig. 8b) provides an explicit personalized **neoantigen prioritization** illustration grounded in a consensus transport substrate. The dataset includes a consensus geometric graph annotated as **“64 nodes and 153 edges,”** a Grover-style marked fraction **“M/N = 8/49 ≈ 0.16,”** and a shortlist table demonstrating the downstream effect of teleport-stabilized consensus formation. This directly targets the manufacturable shortlist constraint: after binding/presentation prediction, processing modeling, and filtering, candidates often compress into near-tie score bands [19–23], while clinical neoantigen vaccine settings require a small synthesizable set [24–28]. The ranking is therefore framed as robust aggregation under redundancy and uncertainty, consistent with manifold ranking and rank-aggregation perspectives [8,9]. The coherent component follows standard quantum-walk formulations [13–16], and the teleport-consensus readout mirrors restart mechanisms used to obtain stable stationary rankings in classical systems [6,7].

#### 4.1.8 Docking ensembles and MWALK transport fingerprints

Docking-ensemble stabilization was evaluated using the comprehensive (SI Appendix I, Figure 9) set (SI Appendix I, Figs. 9d–k; 9f–h; 9j–k) together with the explicit coupling panel in (SI Appendix I, Figure 9l). The dataset instantiation includes Top-10 docking energy landscapes, teleported distribution fingerprints, and multiple composite views linking energetic depth and near-optimal stability to transport-derived basin coherence. Docking and scoring are treated as upstream evidence channels known to exhibit sensitivity and near ties in practical virtual screening and pose ranking (SI Appendices I&II) [18]. The MWALK transport layer provides an orthogonal stability descriptor through entropy/consensus traces and teleported marginals, while (SI Appendix I, Figure 9l) directly visualizes how coherent visitation can diverge from energy-sorted rank order via quantum versus classical walk maps and interference differences (SI Appendix I, Fig. 9l). Coherent-versus-diffusion contrasts follow canonical quantum-walk behavior on graphs [13–16], and the stabilization rationale aligns with restart principles in stationary ranking [6,7].

#### 4.1.9 Flow-cytometry phenotype anchoring across PepMix conditions

Flow-cytometry phenotype anchoring was evaluated using the six PepMix condition panels in (SI Appendix I, Figure 10a–f), (SI Appendices I&II). The dataset instantiation reports CD45^+^ leukocyte recovery and rare CD34^+^ and CD34^+^CD45 ^dim^ compartments per condition, and these summaries are treated as an external biological ordering signal against which computational stability narratives can be compared. The methodological role of this context is to provide an orthogonal phenotype anchor for a near-tie decision landscape, reflecting the broader translational requirement that prioritization outputs should remain interpretable and consistent when confronted with biological readouts beyond the computational evidence stack [24–28]. The transport-layer evaluation logic remains the same: redundancy-rich candidate spaces require robust, auditable aggregation rather than brittle dependence on marginal score differences [1,2].

### 4.2 Robustness and pattern-recognition metrics

We evaluate the method as a **decision system** rather than as a single-metric predictor, consistent with the pattern-recognition view that near-tie regimes should be assessed by **invariance, stability, and auditability under perturbations** rather than marginal gains in a scalar score [1,2]. Across evidence contexts, we apply small, realistic perturbations to (i) the evidence-overlap construction *W*, (ii) the symmetry partition **S**, and (iii) the transport/readout settings, and then quantify whether the resulting shortlist behavior remains basin-consistent.

#### 4.2.1 Top-*K* stability under perturbations

Top-*K* stability is measured as **set overlap** between selected shortlists before and after perturbation (e.g., Jaccard overlap and/or ∣𝒦 ∩ 𝒦_′_∣/*K*). Perturbations include small changes to affinity weights in *W*, alternative but nearby partitions used for quotienting, and docking-landscape summary variations that seed pairwise affinities. This metric matches the **manufacturability constraint** in neoantigen vaccination—where the output is a small set, not an unconstrained ordering—and directly tests whether the method produces basin-level decisions invariant to nuisance variation in evidence scaling and protocol choices [1,2,24–28].

#### 4.2.2 Rank volatility under perturbations

Rank volatility is quantified using **Kendall–***τ* agreement between ranked lists across perturbation conditions [9]. Kendall–*τ* is appropriate in near-tie regimes because absolute score gaps are unreliable, yet small ordinal swaps determine shortlist membership. This aligns with robust rank-integration perspectives where stability is evaluated by ordinal consistency rather than pointwise score calibration [9].

#### 4.2.3 Convergence auditability from polygraph traces

Convergence auditability is evaluated by **plateau behavior** in normalized Shannon entropy and companion dispersion/consensus traces reported in the polygraphs (SI Appendix I, Figs. 2g, 3b, 4a–b, 9a–c). Using *P*(⋅,*t*) as the marginal (classical or measured from *ρ*_*t*_), entropy is

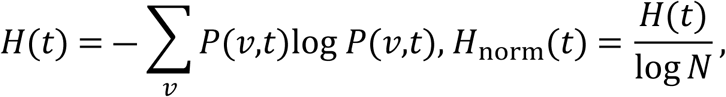

and plateauing is interpreted as evidence that the coupled transport-and-readout mechanism has stabilized into a dominant basin suitable for reproducible ranking [17]. This diagnostic is critical because coherent quantum-walk dynamics are oscillatory and therefore require explicit stabilization (e.g., teleport-consensus mixing) for stationary decision-making [13–16].

#### 4.2.4 Basin compactness and operator–geometry agreement

Basin compactness is assessed by agreement between **operator-domain block structure** (diagonal dominance in the quotient **S**^⊤^*L*_sym_**S**) and **geometry-domain block saturation** in FS-style fidelity kernels (SI Appendix I, Figs. 2c–d, 4f–g, 5a–d). Operator block structure is grounded in spectral graph theory and clustering principles [4,5], while geometry agreement reflects the broader view that Laplacian operators encode manifold structure and similarity geometry [34]. We treat cross-domain agreement as an interpretability criterion: near-tie conclusions are most defensible when multiple independent summaries corroborate the same basin decomposition.

### 4.3 Ablations

Ablations mirror the figure-instantiated pipeline: each control removes or substitutes a specific component that is explicitly depicted in the figures, enabling attribution of stability gains to identifiable design choices.

#### 4.3.1 No quotient

We remove symmetry-aware coarse-graining and operate on *L*_sym_directly (no **S**^⊤^*L*_sym_**S**). This tests whether block-level reduction is necessary to stabilize near-symmetric microstate competition into basin-level decision units, as motivated by spectral operator theory and clustering foundations [4,5].

#### 4.3.2 No teleport

We disable teleport-consensus stabilization and rank using a fixed terminal coherent marginal at a chosen horizon, making the output explicitly horizon-dependent. This tests whether restart/teleport mixing is required to convert oscillatory coherent evidence into a reproducible stationary-style marginal, consistent with restart principles in ranking [6,7] and with the non-convergent nature of coherent quantum-walk dynamics [13–16].

#### 4.3.3 Classical only

We replace coherent transport with diffusion or random-walk-with-restart baselines (RWR) [6–8]. This establishes whether smoothing plus restart alone reproduces basin discrimination achieved by coherent probing, while preserving the stationary stability benefits of restart-based ranking. Diffusion-style relevance propagation is further motivated by manifold-ranking perspectives [8].

#### 4.3.4 Coherent versus diffusion contrasts

We evaluate coherent-versus-diffusion differences directly using the intensity maps and polygraphs in the figure set (SI Appendix I, Figs. 2e–g, 6e–g, 7e–g, 9l–m). The purpose is both comparative and diagnostic: coherent localization bands and oscillatory basin competition provide tie-structure signatures that diffusion tends to smooth away [13–16], and polygraph plateau behavior quantifies whether stabilization yields auditable reductions in uncertainty and spread [17].

## 5. Results

### 5.1 Figure-integrated mechanistic stabilization in nutraceutical peptide targeting

The walnut peptide case (SI Appendix I, Fig. 1) illustrates the intended “phenotype → mechanism → stabilization” arc. Enrichment concentrates in cancer/apoptosis programs (SI Appendix I, Fig. 1A–B) and pathway–gene reuse motivates hub-centric interpretation (SI Appendix I, Fig. 1C). The peptide–target network (SI Appendix I, Fig. 1E) and PPI core (SI Appendix I, Fig. 1F) prioritize hubs such as CASP3 and MMP9 (SI Appendix I, Fig. 1G), providing a mechanistic anchor. However, when multiple peptide–target hypotheses are near-degenerate by docking/predictor scores, pocket microstates are represented as contact graphs and probed by coherent transport overlays (SI Appendix I, Fig. 1H–K). The polygraphs explicitly show that coherent transport localizes earlier and exhibits oscillatory competition compared with diffusion’s smoothing (SI Appendix I, Fig. 1I,K), motivating a restart-stabilized readout for interpretability in the near-tie regime.

### 5.2 Symmetry auditing and teleported consensus yields reproducible basins

Across the three representative pocket cases (SI Appendix I, Fig. 2), the pipeline is consistent: context statistics → *W* (SI Appendix I, Fig. 2a–b) → symmetry-reduced **S**^⊤^*L*_sym_**S** (SI Appendix I, Fig. 2c) → FS kernel block structure (SI Appendix I, Fig. 2d) → coherent intensity *P*(*v*,*t*) (SI Appendix I, Fig. 2e) vs diffusion baseline *p*(*v*,*t*) (SI Appendix I, Fig. 2f) → entropy/spread polygraph (SI Appendix I, Fig. 2g). The key operational stabilization appears in the readout: teleported histograms (SI Appendix I, Fig. 2h) compare terminal coherent marginals to restart-mixed distributions under the annotated “**p=α≈0.18**.” This converts horizon-dependent oscillations into a stationary-style consensus basin suitable for ranking.

A practical interpretation is that the teleported distribution preserves the coherent basin structure (bands in Fig. 2e) while preventing over-sensitivity to terminal time and initialization—precisely the failure mode in near-tie shortlist bands.

### 5.3 MWALK target stabilization: rapid entropy/spread saturation and consensus plateaus

The MWALK CRC target composite (SI Appendix I, Fig. 3) shows how heterogeneous upstream evidence (QSAR, structural snapshots, MD-style traces) is converted into transport signatures. The reported pattern is that *H*_norm_(*t*) and spread rise sharply and saturate near 1.0 within ∼5–10 steps, while the consensus stabilizes to “∼0.65–0.75” (SI Appendix I, Fig. 3b). This provides a compact indicator: fast saturation + stable plateau corresponds to a dominant, reproducible basin (robust tie-breaking), whereas slower or overshooting traces correspond to weaker basin separation (tie risk).

### 5.4 Multi-cohort and radiogenomic modules: controller + PASS/teleport audit trails

The multi-cohort controller examples (SI Appendix I, Fig. 4a) explicitly show a depth-40 coined walk “Repeat ×40” and a TELEPORT + CONSENSUS module, with per-run parameter annotations (θ, p, and additional scalars). Entropy polygraphs plateau as the teleport channel reinforces dominant basins (SI Appendix I, Fig. 4a Column 4). The discrete vs continuous-time diagnostics and PASS-oracle teleport logs provide a run-by-run audit trail of transport behavior and restart handoffs (SI Appendix I, Fig. 4b). In radiogenomic module inference, symmetry-audited quotient operators and CTQW intensity maps reveal block-localized occupancy vs mixing, supporting module-level interpretability rather than single-gene over-claiming (SI Appendix I, Fig. 4c–d).

### 5.5 Docking landscapes + teleported fingerprints separate “deep-but-variable” from “shallow-but-stable”

Docking best-pose minima are necessary but not sufficient for robust selection [18]. The HPEPDOCK composites explicitly separate energetic depth (rank-1 minima) from near-optimal stability (Top-10 spreads and Top1–Top2 gaps) (SI Appendices I&II) and then add a transport-stability axis via teleported fingerprints (SI Appendix I, Figs. 9d–h, 9j–k). For example, targets with deep minima can still exhibit steep early drop-offs (large Top1–Top2 gaps), indicating narrow basins that are sensitive to perturbations (SI Appendix I, Fig. 9e). The teleport-regularized MWALK histograms provide an orthogonal readout: whether probability concentrates into a dominant presentation mode or remains broadly redistributed (SI Appendix I, Figs. 9d–e, 9h).

### 5.6 Neoantigen shortlist illustration: consensus topology → stable ranking under near ties

The neoantigen-style example explicitly connects a consensus graph to a downstream shortlist (SI Appendix I, Fig. 8b). Panel A describes a consensus manifold with “**64 nodes and 153 edges**,” and the transport readout uses teleport mixing *ρ*_*t*+1_ = (1 ― *α*)*Uρ*_*t*_*U*^†^ + *αT*(*ρ*_*t*_) to yield a reproducible distribution. Panel B illustrates a classical vs quantum-adjusted weighting shift and a Grover marked fraction “**M/N=8/49≈0.16**,” emphasizing that the quantum-adjusted shortlist is a stabilization layer acting in a compressed near-tie landscape rather than a replacement for binding/immunogenicity evidence.

### 5.7 Flow cytometry phenotype anchoring across PepMix conditions

Across PepMix1–PepMix6 (SI Appendices I&II), flow cytometry shows preserved CD45^+^ recovery but condition-dependent shifts in rare CD34^+^ and CD34^+^CD45^dim^ compartments (SI Appendix I, Fig. 10a–f), (SI Appendices I&II). The ordering used as an external phenotype axis is:

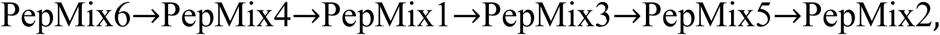

with PepMix6 the lowest CD34^+^CD45^dim^ retention and PepMix2 the highest (SI Appendix I, Fig. 10f vs Fig. 10b). PepMix3 is explicitly an outlier with enriched CD34^+^ but low conversion into the CD45^dim^ definition, indicating heterogeneous phenotype rather than monotone depletion (SI Appendix I, Fig. 10c), (SI Appendices I&II). This phenotype gradient is used as an interpretability constraint: rankings are preferred when near-tie tie-breaks do not contradict robust biological ordering signals.

## 6. Discussion

### 6.1 Why teleport-stabilized coherent transport is a pattern-recognition contribution

Near-tie ranking instability is fundamentally a **pattern recognition** challenge: the goal is stable decision-making under correlated evidence and redundant representations [1]. Graph formulations provide a principled structure for overlap and redundancy [3,33]. Spectral normalization and clustering impose invariances [4,5,32], while teleport/restart mechanisms enforce convergent behavior and reduce sensitivity to traps or disconnected regions [6,7]. Quantum walks add discriminative transport fingerprints but must be regularized for decision use [13–16]. The figure set collectively instantiates this logic: coherent vs diffusive separations (SI Appendix I, Figs. 2e–g, 6e–g, 9m) are made decision-ready by teleport-consensus stabilization (SI Appendix I, Figs. 2h, 8a, 9a–c).

### 6.2 Interpretability and auditability: why quotient + polygraphs matter

The quotient operator turns microstate-level redundancy into basin-level units that can be explained (SI Appendix I, Figs. 2c, 4d, 5a–d). Polygraphs provide a reproducible audit record: early entropy rise indicates exploration; plateau indicates stabilization; overshoot suggests multi-basin competition (SI Appendix I, Figs. 2g, 3b, 4a–b). PASS/teleport logs (SI Appendix I, Fig. 4b–c) further support traceability.

### 6.3 Practical insertion point in neoantigen pipelines

The proposed layer is intended for insertion **after conventional neoantigen candidate generation and first-pass prioritization, but before irreversible final shortlist selection and formulation**, i.e., as a downstream stabilization module rather than as a replacement for established predictors. In a standard pipeline, candidate peptides are first generated from somatic variation and filtered by presentation-oriented models such as NetMHCpan-4.1 and MHCflurry 2.0, which respectively provide “improved predictions of MHC antigen presentation” and class-I peptide prediction by “incorporating antigen processing” [19,20]; these candidates are then further refined by proteogenomic and presentation-aware prioritization frameworks [21–23] and interpreted within the broader clinical logic of neoantigen immunotherapy [24–28]. At this stage, however, the surviving candidates commonly occupy a compressed near-tie regime in which scalar ranking alone is fragile. Our module is designed specifically for that late-stage regime: once the candidate band has been narrowed by binding/presentation evidence [19,20], proteogenomic support [21–23], and clinical neoantigen rationale [24–28], we construct a weighted overlap graph whose nodes represent peptides, peptide–HLA pairs, or closely related peptide–microstate hypotheses, and whose edges encode structural, evidential, or functional overlap. This representation is consistent with the broader machine-learning transition from classical pattern-recognition pipelines [1] and deep learning [2] toward relational and graph-based learning [3,29–31], and it leverages the well-established spectral-graph framework in which the symmetric normalized Laplacian *L*_sym_ = *D*^―1/2^(*D* ― *W*)*D*^―1/2^ provides a principled geometry for clustering, diffusion, and ranking [4,5,33–35]. After graph construction, we apply symmetry-aware reduction, compute coherent transport fingerprints, and stabilize the terminal readout with teleport-consensus before final top-*K* selection. In this sense, the layer acts as a graph-geometric tie-breaker: rather than asking only which peptide has the highest raw score, it asks which peptide remains the most stable representative of a coherent basin once redundancy, neighborhood structure, and microstate continuity are taken into account. This rationale is closely related to graph-ranking and manifold-ranking ideas [6–9], particularly restart-based ranking schemes derived from PageRank, originally framed as “bringing order to the web” [6] and later generalized well beyond that context [7], and to robust integration strategies for compressed candidate lists [8,9]. It is also motivated by the known instability of late-stage scoring, calibration, and uncertainty estimation in modern predictive systems [10–12] and by the limitations of docking/scoring alone in compressed virtual-screening regimes [18]. The transport layer then provides an orthogonal stability axis: coherent graph propagation follows the quantum-walk formalism introduced in “quantum walks on graphs” [13] and developed in later reviews [14–16], while uncertainty and stabilization are summarized through entropy- and spread-based diagnostics grounded in Shannon’s “mathematical theory of communication” [17]. Within the manuscript, this intended insertion point is illustrated across several evidence settings: Figs. 1–3 show how heterogeneous pathway, docking, QSAR, and MD-derived evidence can be recast into graph and transport descriptors; Figs. 4f, 4g, 5a, 5c, and 5d show that quotient operators and fidelity kernels can separate coherent candidate communities even when raw evidence remains near-degenerate; and Figs. 8b, 9q, 9r, 9s, and 9t most directly illustrate the neoantigen-facing use case, where a pre-ranked candidate set is converted into a consensus graph, teleport-regularized transport identifies dominant basin representatives, and the resulting terminal probability mass serves as an auditable tie-break signal for shortlist stabilization. Accordingly, the practical sequence is: generate candidates conventionally; score them with binding, presentation, and proteogenomic evidence [19–23]; interpret them within the clinical neoantigen framework [24–28]; then build the overlap graph, apply spectral normalization and symmetry-aware reduction [4,5,32–35], evaluate coherent transport [13–16], stabilize the readout through restart/teleport-style consensus [6,7], and only then commit to final top-*K* selection. In this way, the proposed layer is fully compatible with current neoantigen workflows while directly targeting a genuine practical fragility of modern personalized immunotherapy pipelines: late-stage decision-making among candidates whose upstream biological evidence is strong but insufficiently separated.

## 7. Conclusions and future work

We presented a transport-stabilized ranking layer for near-tie regimes in personalized neoantigen selection and peptide–target prioritization, where the operational goal is not to declare a uniquely optimal candidate in isolation but to advance a *small, manufacturable* top-*K* set whose membership is resilient to realistic perturbations (predictor updates, calibration shifts, feature rescaling, docking protocol variation, or correlated evidence channels) [1,2,18]. The framework is built around four coupled ideas: (i) redundancy is modeled explicitly through an evidence-overlap graph, (ii) near-symmetries that drive fragile microstate competition are reduced via quotient operators grounded in spectral graph theory, (iii) basin structure is probed using coherent quantum-walk transport to expose non-diffusive separability and competition, and (iv) horizon-dependent coherent evidence is converted into a reproducible decision marginal through teleport-consensus mixing that yields stable, comparable readouts across perturbations and contexts [4–7,13–16]. Across the provided figure suite, these components form an auditable ranking substrate: coherent-versus-diffusive contrasts separate basin structure from generic smoothing (SI Appendix I, Figs. 2, 6, 9m), teleport stabilization yields consensus readouts (SI Appendix I, Figs. 2h, 8a, 9a–c), and entropy/spread polygraphs provide traceable stabilization diagnostics that expose whether a decision is basin-dominant or tie-risk (SI Appendix I, Figs. 2g, 3b, 4a–b) [17].

### 7.1 What is concluded scientifically: a stability-first decision layer for redundancy-rich biology

A central conclusion is that shortlist instability in late-stage neoantigen and peptide–target pipelines is often not primarily an accuracy problem but a *representation and decision-logic* problem: once candidates are compressed into narrow score bands, the dominant uncertainty becomes *relative ordering under redundancy*, not absolute ranking under an assumed independent-score model [1,2]. Our results support the view that the appropriate object of inference in near-tie regimes is the *basin* (a redundancy-defined decision unit) rather than the single node (a candidate peptide, ligand, or microstate). This shift is operationally consequential because manufacturable decisions are basin-like decisions in practice: clinical manufacturing selects a handful of peptides, and if several candidates are effectively interchangeable under the available evidence, the pipeline’s responsibility is to make that interchangeability explicit, quantify it, and stabilize the final selection around it rather than allowing arbitrary micro-perturbations to dictate outcomes.

The evidence-overlap graph provides the mechanism for this reframing. Instead of treating each candidate’s score as an independent scalar, the framework treats candidates as nodes embedded in a structured neighborhood where edges encode shared evidence and mechanistic similarity (motifs, alleles, processing features, docking microstates, pathway neighborhoods) [3,33]. In this representation, near ties are no longer “small numerical differences”; they appear as *redundancy structure*—dense neighborhoods and near-symmetric clusters that can be detected, audited, and (critically) *coarse-grained* in a controlled way.

The quotient operator step is the second scientific conclusion: near ties behave like approximate symmetries, and symmetry-aware coarse-graining stabilizes decisions by collapsing brittle microstate competition into basin-level units without discarding the information needed for discrimination [4,5]. This is not an ad hoc merge; it is an operator-domain transformation that reduces sensitivity to within-basin permutations while preserving between-basin structure in a way that can be inspected through quotient heatmaps and kernel fidelity structure (e.g., Figs. 2c–d, 4f–g, 5a–d). The operator viewpoint is especially important in near-tie regimes because it distinguishes two qualitatively different scenarios that scalar ranks often conflate: (i) candidates are near tied because they are essentially interchangeable under the measured evidence (a symmetry-driven tie), versus (ii) candidates are near tied because evidence is weak or inconsistent (an uncertainty-driven tie). Collapsing (i) is desirable; diagnosing (ii) is essential.

The third conclusion concerns transport. Diffusion-based propagation is an excellent baseline for denoising and monotone mixing, but it can be insufficient as a basin discriminator precisely when basins are close and the decision is dominated by fine-grained competition. Coherent quantum-walk transport preserves interference structure and can reveal transient localization and oscillatory basin competition that diffusion intentionally smooths away [13–16]. The figure suite repeatedly illustrates this qualitative distinction: coherent intensity maps show localized, structured visitation patterns and competition, whereas diffusion shows monotone relaxation (SI Appendix I, Figs. 2e–f), and the polygraphs reveal different stabilization signatures (SI Appendix I, Fig. 2g). The methodological claim supported here is not that coherence is universally superior, but that *coherence is informative* in the near-tie regime because it acts as a structural probe: it highlights whether the graph contains sharply separable basins or a diffuse, ambiguous manifold.

The final conclusion is that transport evidence must be made *decision-stable*. Coherent dynamics are horizon-dependent and non-convergent, which is scientifically informative but operationally unusable unless converted into a reproducible readout. Teleport-consensus mixing performs exactly that conversion: it turns oscillatory transport evidence into a stationary marginal distribution that can be ranked, compared across perturbations, and reported as an auditable artifact (SI Appendix I, Figs. 2h, 8a, 9a–c) [6,7]. This is a crucial point of differentiation from many “quantum-inspired” or spectral heuristics: the framework does not stop at producing an interesting dynamic signature; it explicitly builds the bridge from dynamic probe to stable decision substrate, with polygraph diagnostics documenting whether and when stabilization occurs (SI Appendix I, Figs. 2g, 3b, 4a–b) [17].

### 7.2 Comparative strengths versus common alternatives

A key strength of the methodology is that it is designed for the *late pipeline* regime—after standard filtering has already produced a narrow band of plausible candidates—where many pipelines become fragile because they continue to behave as if each candidate is independent and totally ordered by a single score. In that regime, attempting to “win” by improving a single predictor often yields diminishing returns: small absolute improvements do not necessarily translate into stable top-*K* membership because the shortlist is dominated by correlated evidence and redundancy-driven interchangeability [1,2]. The transport-stabilized layer strengthens the pipeline by shifting optimization from “marginally better scores” to “more stable decisions,” which is the correct objective when downstream manufacturing or experimental validation capacity is the limiting resource.

#### Compared to diffusion-only ranking and restart propagation

Random walks, heat diffusion, and restart-based methods provide mature and interpretable baselines, often yielding stable stationary distributions that can be used for ranking [6–8,34]. Their limitation in near-tie microstate competitions is that stability is frequently achieved through smoothing, which can erase the very basin boundaries that determine which candidates are meaningfully distinct. Our framework retains diffusion as a comparator and ablation class (Section 4.3), but it adds two differentiators: quotient reduction that explicitly handles symmetry-driven ties at the operator level [4,5], and coherent transport that exposes basin structure via non-diffusive dynamics before stabilization [13–16]. This pairing is important: quotient reduction alone can stabilize microstate competition but may still leave between-basin ambiguity; coherent probing exposes whether that ambiguity is structural, and teleport-consensus then converts that exposure into a stable readout.

#### Compared to docking-only or single-score structure pipelines

Docking refinement is indispensable but known to be sensitive to scoring function choice, sampling, and protocol parameters, especially when candidates are close in score [18]. The figure suite highlights precisely the scenario where docking can be strong yet still produce a fragile order: multiple plausible microstates or near-equal minima exist, and small perturbations can reorder the top entries even when biological interpretation does not change. Rather than replacing docking, the transport layer strengthens the decision by (i) encoding pose/contact consistency and related statistics into *W*, (ii) auditing symmetry and basin structure (SI Appendix I, Fig. 2b–d), and (iii) producing a stabilized consensus marginal that is less sensitive to horizon selection and microstate permutations (SI Appendix I, Fig. 2h). The method therefore contributes a second, stability-oriented axis—basin dominance and consensus—alongside energetics (SI Appendix I, Fig. 9l). This is a practical advantage: it allows an investigator to distinguish “a single deep minimum that is protocol-fragile” from “a coherent basin that is robust under perturbation,” which is not always obvious from energies alone (SI Appendix I, Figs. 9d–k).

#### Compared to end-to-end learned rankers

Deep rankers and graph neural networks can be powerful when training data and labels are adequate [2,29–31]. However, patient-specific neoantigen outcomes are sparse, confounded, and difficult to assign at the candidate level, and opaque models can output confident rankings without revealing whether the decision is supported by a dominant basin or is effectively arbitrary within a near-tie band. The transport-stabilized layer is compatible with learning—*W* and the personalization vector can be learned or parameterized—but it preserves an explicit audit trail through quotient structure, transport contrasts, consensus stabilization, and polygraph diagnostics. This “glass-box stability layer” is a comparative strength in translational contexts where explainability and robustness matter as much as raw predictive performance, and where uncertainty estimation and calibration are known to be essential for reliable decision-making [10–12].

### 7.3 Why the figure-linked results matter operationally: auditability and decision governance

Across the figure suite, the methodology does more than produce a final rank list; it produces *evidence about the ranking itself*. This is the core of auditability: the system reports not only “who is top-*K*” but also “how stable is top-*K* under perturbations,” “is the decision basin-dominant or tie-risk,” and “what structural features drive interchangeability.” Coherent-versus-diffusive contrasts (SI Appendix I, Figs. 2, 6, 9m) function as structural probes; teleport-consensus readouts (SI Appendix I, Figs. 2h, 8a, 9a–c) function as stable decision marginals; and entropy/spread polygraphs (SI Appendix I, Figs. 2g, 3b, 4a–b) function as convergence and tie-risk diagnostics [17]. This separation of *probe → stabilize → diagnose*provides a governance framework for near-tie decisions: if diagnostics show a stable plateau and basin dominance, one can proceed with higher confidence; if diagnostics show persistent dispersion and horizon sensitivity, the pipeline can justify additional validation, expanded manufacturing when feasible, or explicit contingency candidates rather than silently committing to an arbitrary micro-order.

This governance perspective is particularly relevant for personalized neoantigen vaccination, where the shortlist is both small and consequential [24–28]. Two pipelines that output different top-*K* sets due to minor version changes create a translational risk: manufacturing, regulatory documentation, and downstream immunomonitoring can diverge even when underlying biology is effectively unchanged. The method’s stability-first logic addresses this risk by explicitly representing redundancy, reducing symmetry-driven ties, stabilizing coherent evidence, and reporting diagnostics that can be archived and compared across runs. In this way, the framework supports reproducibility not only in model performance but also in *decision behavior*, which is the clinically relevant unit of reproducibility.

### 7.4 Limitations and scope conditions

The framework is not intended as a replacement for upstream biological modeling, but as a decision layer that operates after candidate generation and preliminary scoring. Its effectiveness depends on the quality and relevance of the evidence encoded in *W*: if the overlap graph is poorly constructed or dominated by irrelevant similarity, the transport dynamics will probe and stabilize the wrong structure. Likewise, quotient reduction requires a defensible block structure; overly aggressive coarse-graining can mask meaningful distinctions, while overly fine partitions can fail to remove brittle microstate competition. The figure-faithful ablations (Section 4.3) therefore remain essential: “no quotient” tests whether symmetry reduction is doing meaningful stabilization; “no teleport” tests whether the coherent readout remains horizon-sensitive; and diffusion-only baselines test whether the benefits derive from coherence or simply from restart smoothing [6–8].

Coherent transport also introduces modeling choices (e.g., Hamiltonian construction, time horizon sampling) that can be misused if treated as tuning knobs rather than as probes. The methodological stance advocated here is to treat coherence as a diagnostic and basin probe and to rely on teleport-consensus and polygraphs as the stabilizing and auditing mechanisms that prevent horizon arbitrariness from leaking into the final decision. In other words, the framework’s practical validity is strongest when coherence is used to *reveal* structure and teleport-consensus is used to *decide* robustly.

### 7.5 Future work

Future work will focus on three complementary directions that strengthen both robustness and translational readiness.

a. **Uncertainty-aware priors and calibrated evidence fusion.** Near-tie decisions are fundamentally uncertainty-limited. Incorporating calibrated uncertainty into the personalization vector and the overlap graph—so that edges reflect both similarity and confidence—should improve stability and interpretability, particularly when evidence channels are correlated or partially missing [10–12]. This direction includes explicit uncertainty propagation into polygraph diagnostics, allowing convergence plateaus to be interpreted in calibrated probabilistic terms rather than purely descriptively.
b. **Learning** *W* **and** *v* **under stability-oriented objectives.** Graph representation learning offers principled ways to learn similarity structure from heterogeneous evidence [2,29–31], but standard objectives emphasize predictive fit rather than decision stability. A key opportunity is to train or tune *W* (and any personalization vector) using stability-oriented losses—directly optimizing top-*K* robustness, reducing rank volatility, and encouraging basin dominance when evidence supports it—while preserving auditability through quotient and transport diagnostics.
c. **Standardized reporting of near-tie robustness.** The field would benefit from reporting standards that treat near-tie behavior as a first-class outcome rather than an afterthought. At minimum, this includes reporting top-*K* stability under perturbations, rank volatility summaries such as Kendall-*τ* agreement [9], and convergence signatures from entropy/spread/consensus polygraphs that document whether stabilization was achieved and whether the decision is basin-dominant or tie-risk (SI Appendix I, Figs. 2g, 3b, 4a–b) [17]. Such standards would enable meaningful comparison across pipelines, institutions, and trials, improving reproducibility at the level that matters most: the shortlist that goes into manufacturing and patient treatment.

Taken together, these conclusions support a general claim: transport-stabilized ranking is most valuable exactly where modern pipelines struggle most—when many candidates remain plausible, scores are compressed, and redundancy makes the top-*K* decision fragile. By explicitly modeling redundancy, reducing symmetry-driven ties, probing basin structure with coherent transport, and stabilizing decisions with teleport-consensus mixing under polygraph audit, the framework provides a principled and operationally reproducible pathway from heterogeneous evidence to manufacturable, interpretable, and stability-aware shortlist decisions across the figure-linked contexts (SI Appendix I, Figs. 1–4, 6–10).

## Data Availability Statement

The datasets generated and/or analyzed during the current study have not been deposited in a public repository. Underlying data supporting the findings—including intermediate computational observables from the evidence-graph and transport workflow (e.g., peptide–HLA candidate tables, graph affinity/overlap matrices *W*, degree/operator constructs (e.g., *L*□γ□, quotient operators), symmetry/cluster assignment maps, diffusion and quantum-walk transport traces, teleport-consensus (restart-mixed) marginal distributions, entropy/dispersion “polygraph” diagnostics, docking-ensemble summaries and microstate/contact fingerprints where applicable, and figure-source matrices)—are available from the corresponding author upon reasonable request. Where applicable, underlying intermediate files, including the graph *W*, quotient operators, transport traces, and summary tables, will be deposited in an accessible repository and linked in the final submission. In addition, the code associated with this work has been made publicly available on GitHub at **Colonyvaq-Neopersonaq**: https://github.com/GRIGORIADIS1979/Colonyvaq-Neopersonaq-.

## Code Availability Statement

Custom code developed for this study has not been publicly released in a code repository. The core implementation (including evidence-kernel fusion and graph construction, symmetry-aware quotient reduction routines, diffusion/RWR baselines, coherent quantum-walk simulation modules (CTQW/DTQW), teleport-consensus (restart-mixed) stabilization and readout, and polygraph diagnostic/visualization scripts) is available from the corresponding author upon reasonable request, subject to any third-party licensing constraints for external software or services used in parts of the workflow.

## Consent for Publication

All authors have read and approved the manuscript for submission and publication (in print and/or online, including any supplementary materials). The manuscript does not contain any individual person’s identifiable data, images, or personal information requiring specific consent; therefore, consent for publication of individual-level personal data is not applicable. All necessary permissions for the use and publication of any third-party material have been obtained, and appropriate acknowledgements and/or citations are included. The corresponding author is authorized to act on behalf of all authors with respect to publication-related communications, including provision of underlying data and custom code upon reasonable request as described in the Data Availability and Code Availability Statements.

## Ethics Approval and Consent to Participate

### Ethics Approval

This study involved no human participants, no prospective recruitment, and no animal experiments. Analyses were conducted using computational methods (including in silico modeling and quantum-walk transport simulations). Where patient-derived molecular signatures or neoantigen candidates are referenced conceptually, the manuscript does not report identifiable personal data. Accordingly, ethical approval from an institutional review board or ethics committee was not applicable for the work as presented.

### Consent to Participate

Consent to participate was not applicable, as the study did not involve recruitment, clinical intervention, or collection/processing of identifiable personal data.

### Author Contributions

**I.G.** conceived the study; developed the methodology; implemented the software; performed the computational experiments; curated and analysed the data; created the visualizations; and drafted and revised the manuscript. **C.E.** contributed clinical-scientific interpretation and domain review, and participated in manuscript review and editing.

## Acknowledgements

The corresponding author I.G. acknowledge Prof. **Christos Emmanouilides** MD Phd and the **Interbalkan Medical Center (Pylaia, Thessaloniki, Greece)** for institutional context and support. The authors also acknowledge internal resources and support from **Biogenetoligandorol™ & Synthocure™ Stations**, the **Department of BiogenetoligandoroMACHNOT/QIICDNNDCA & Biogenea™ Stations**, and the **Cellgenea™ Stem Cell Expansion Department; Neopersonaq™** that contributed to completion of this work.

---

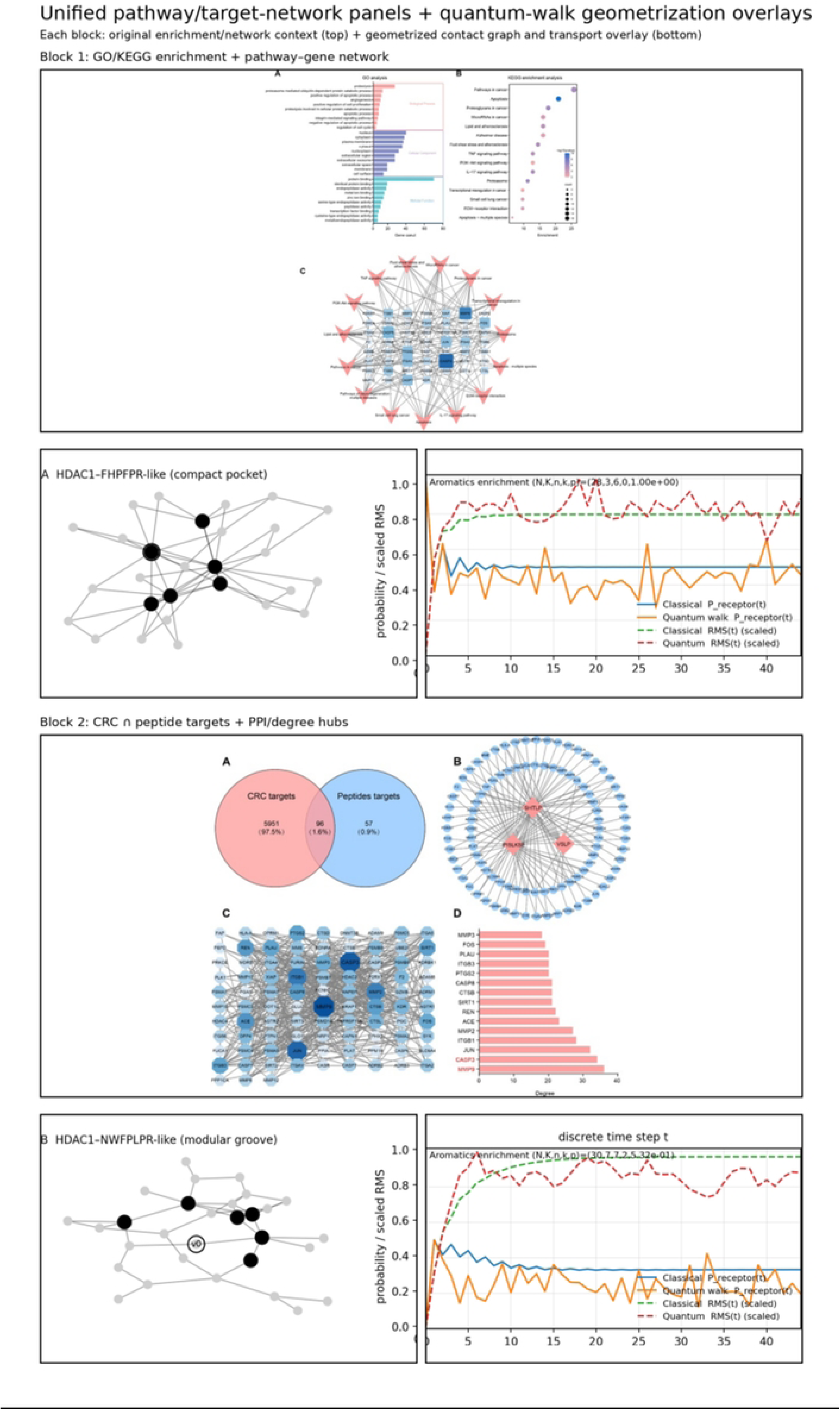

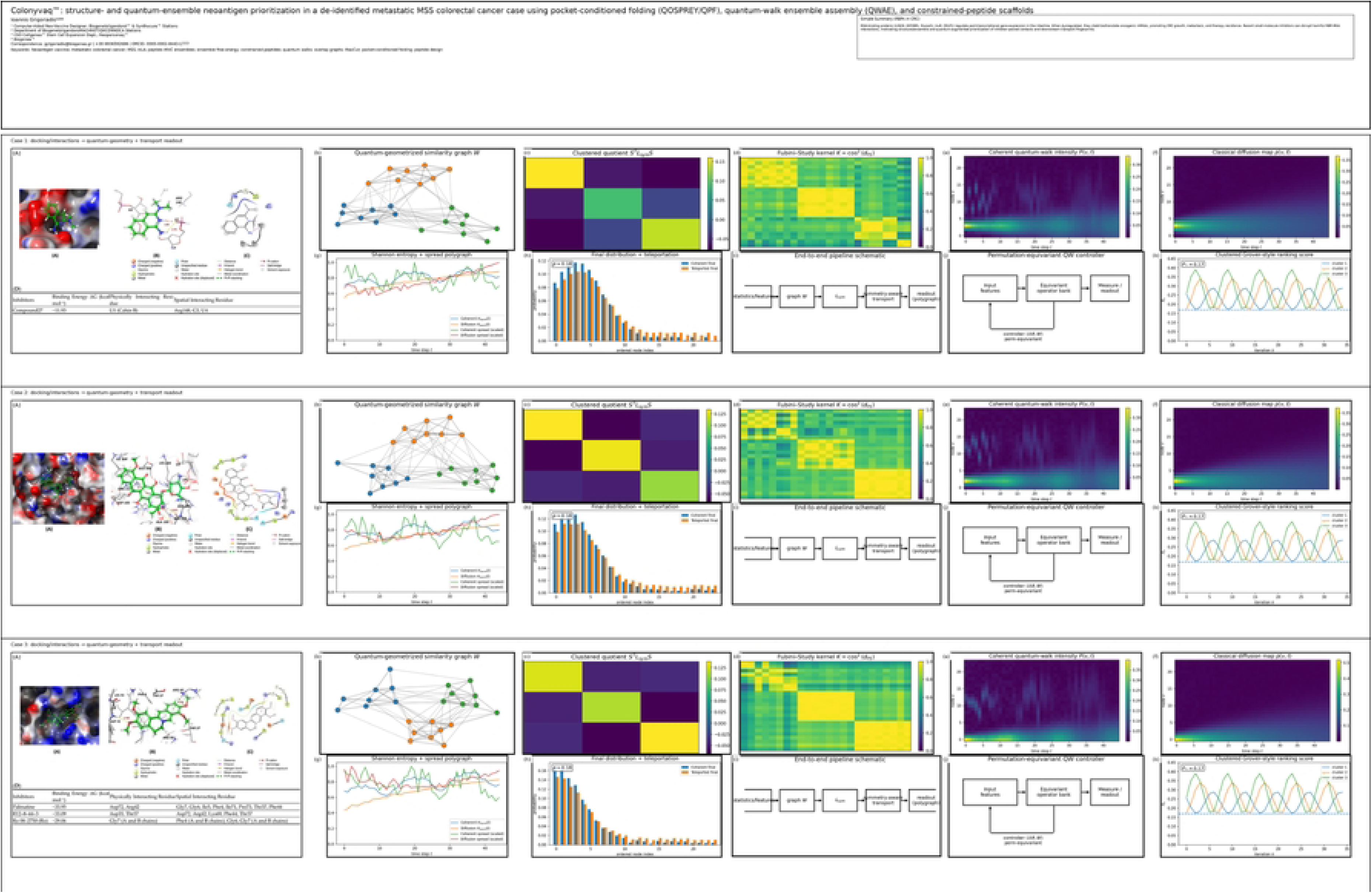

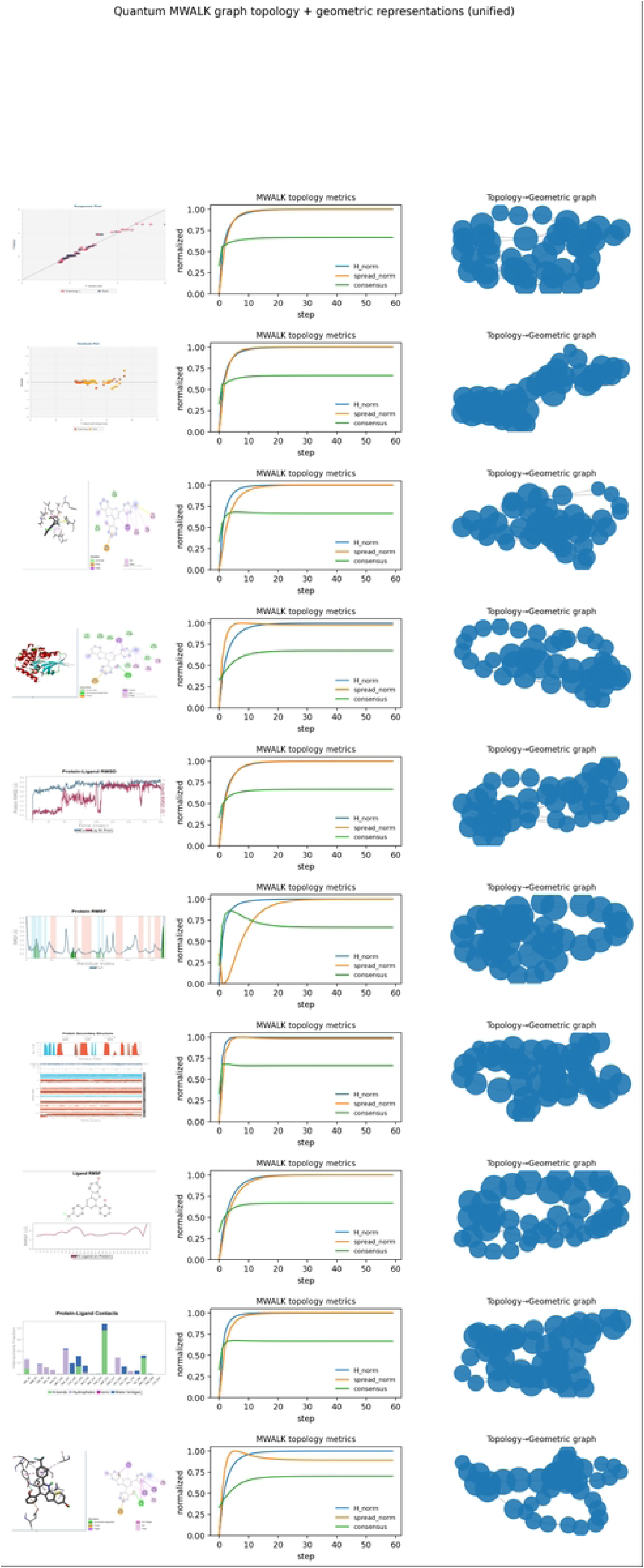

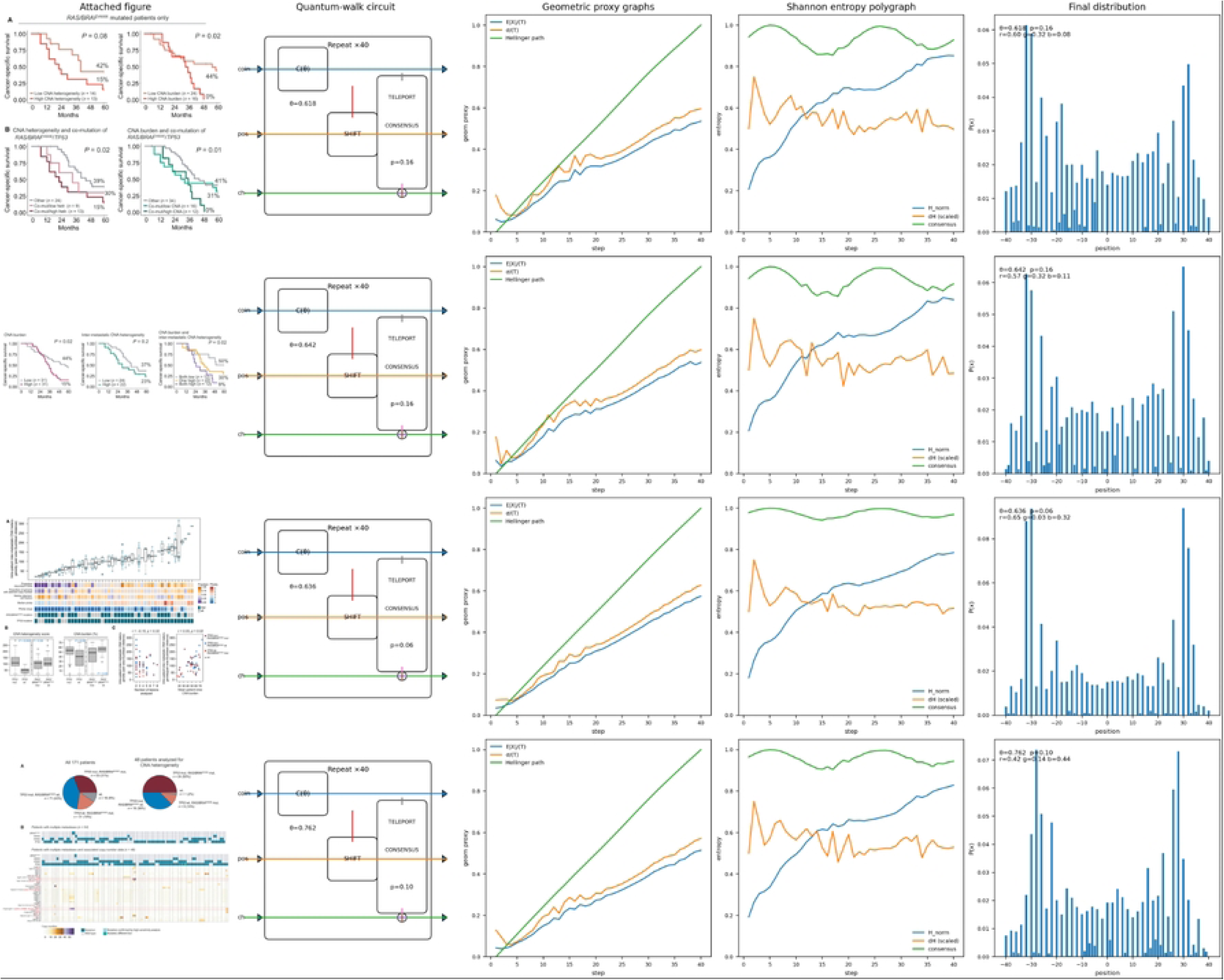

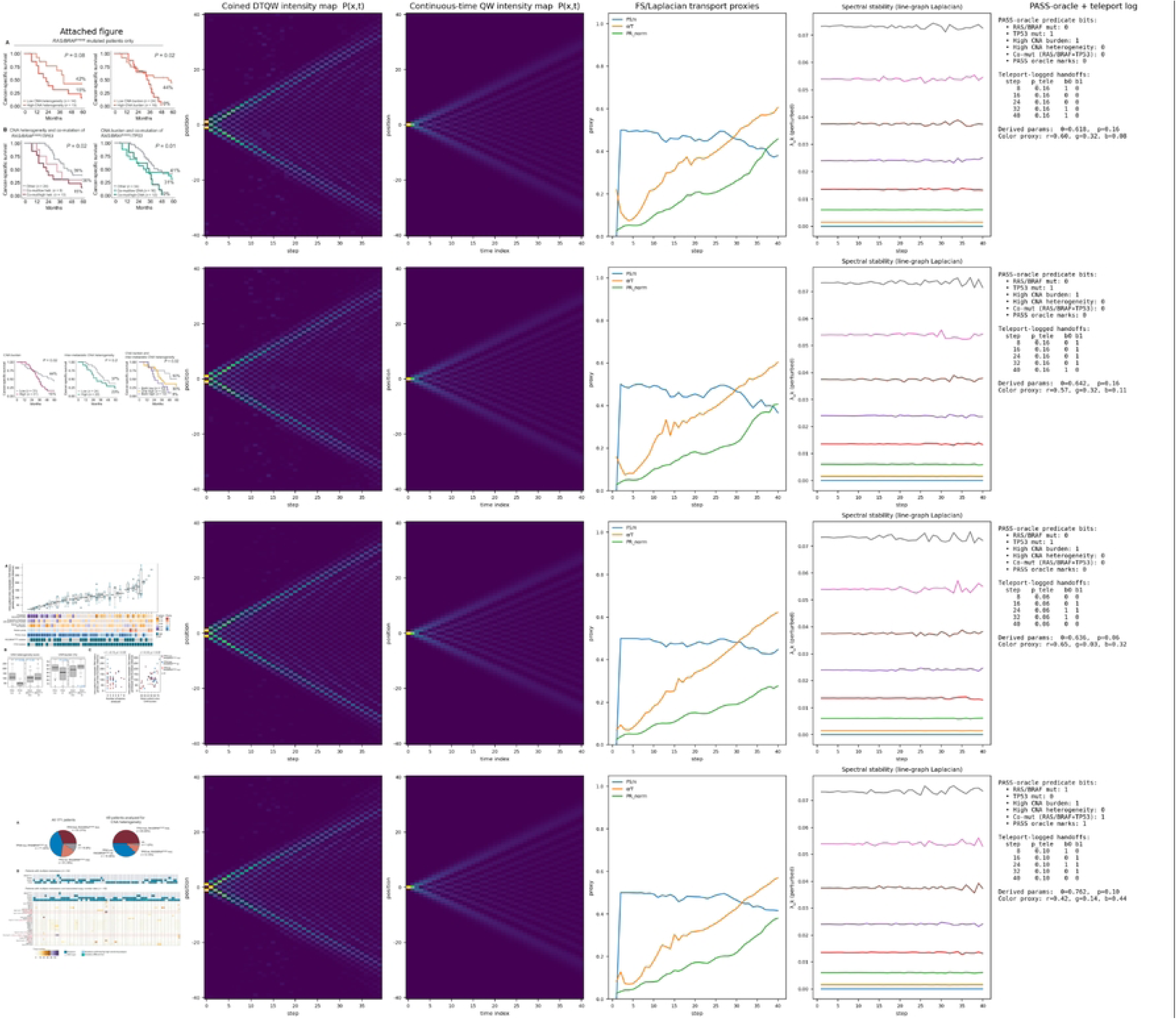

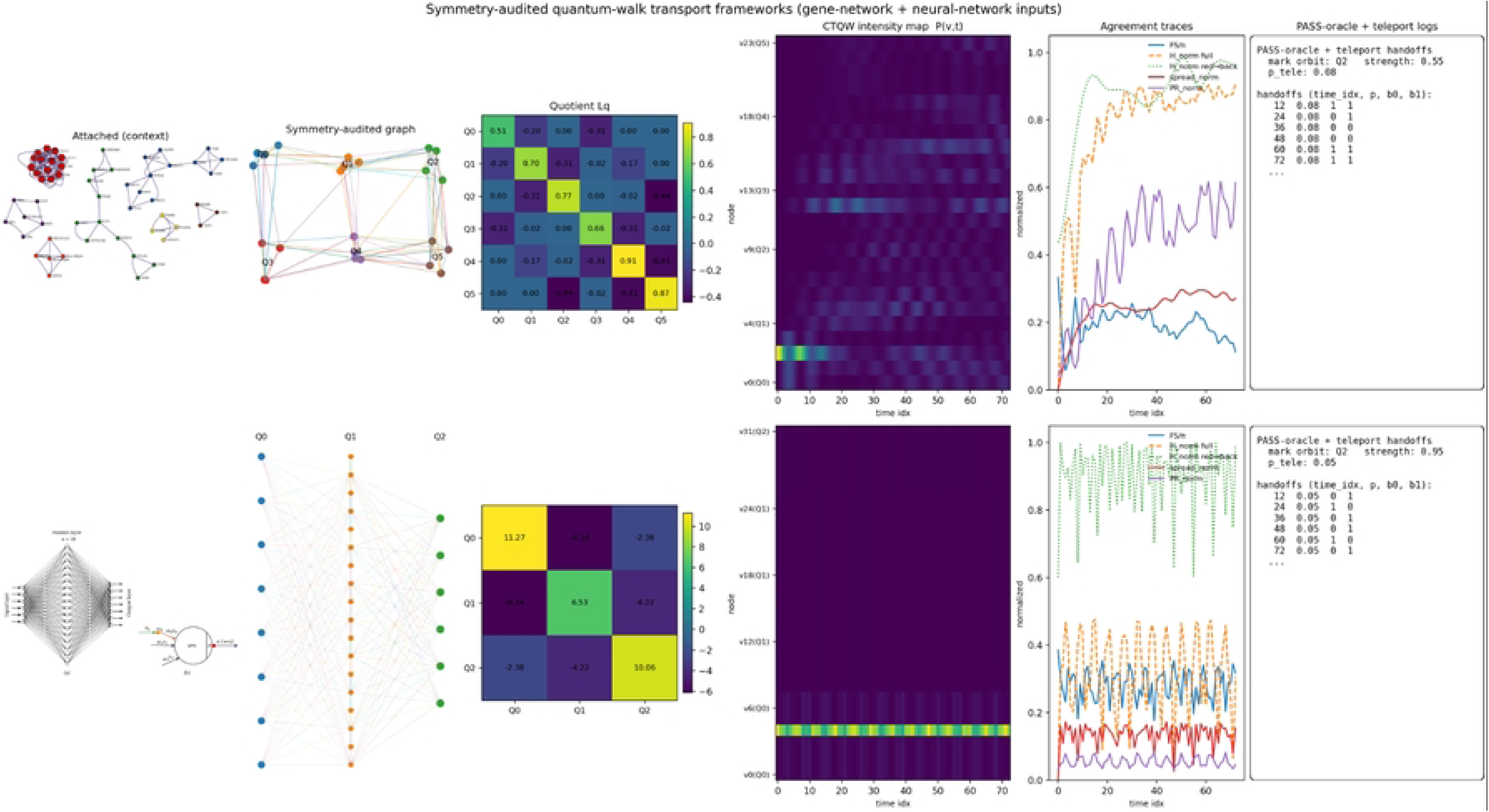

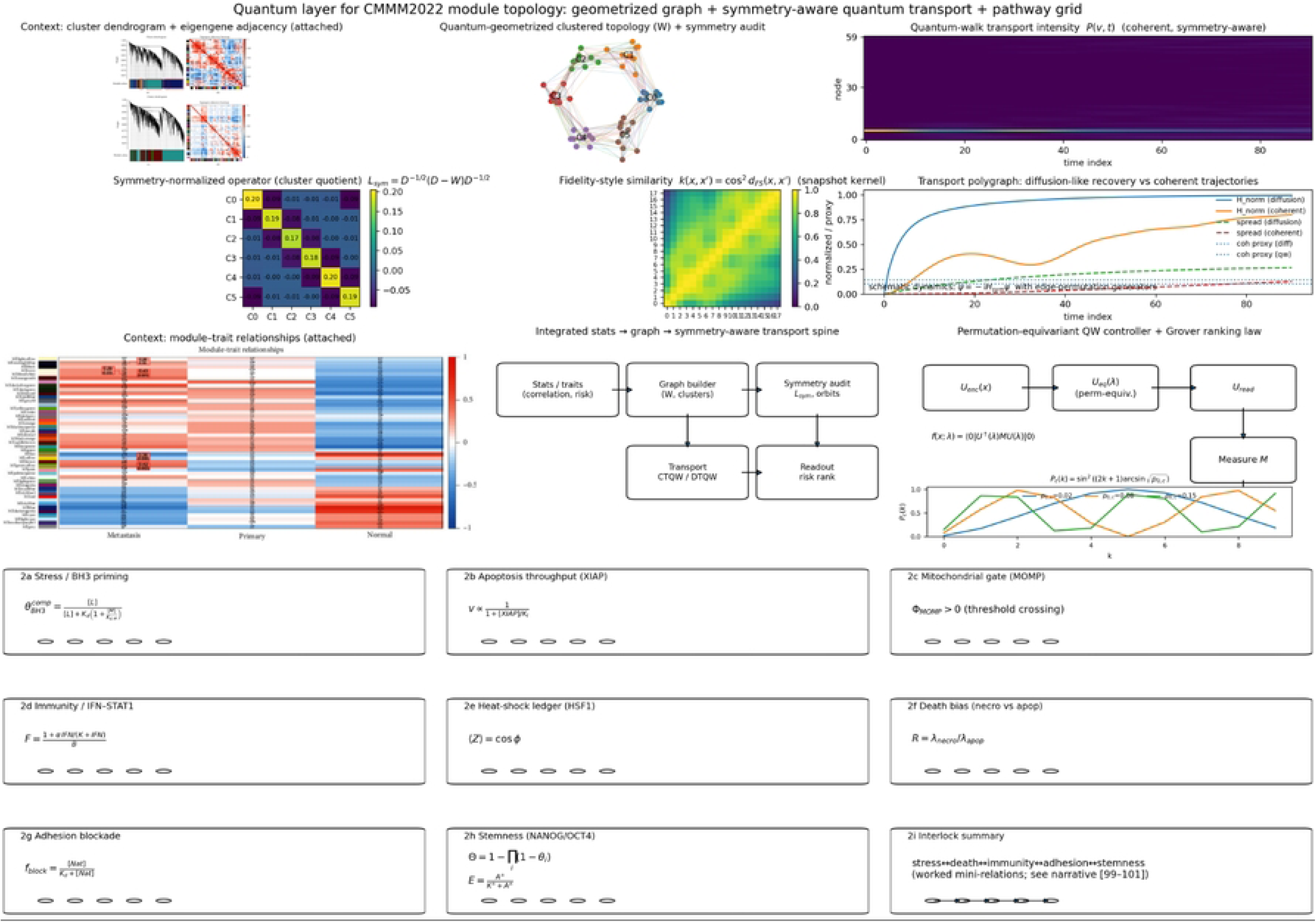

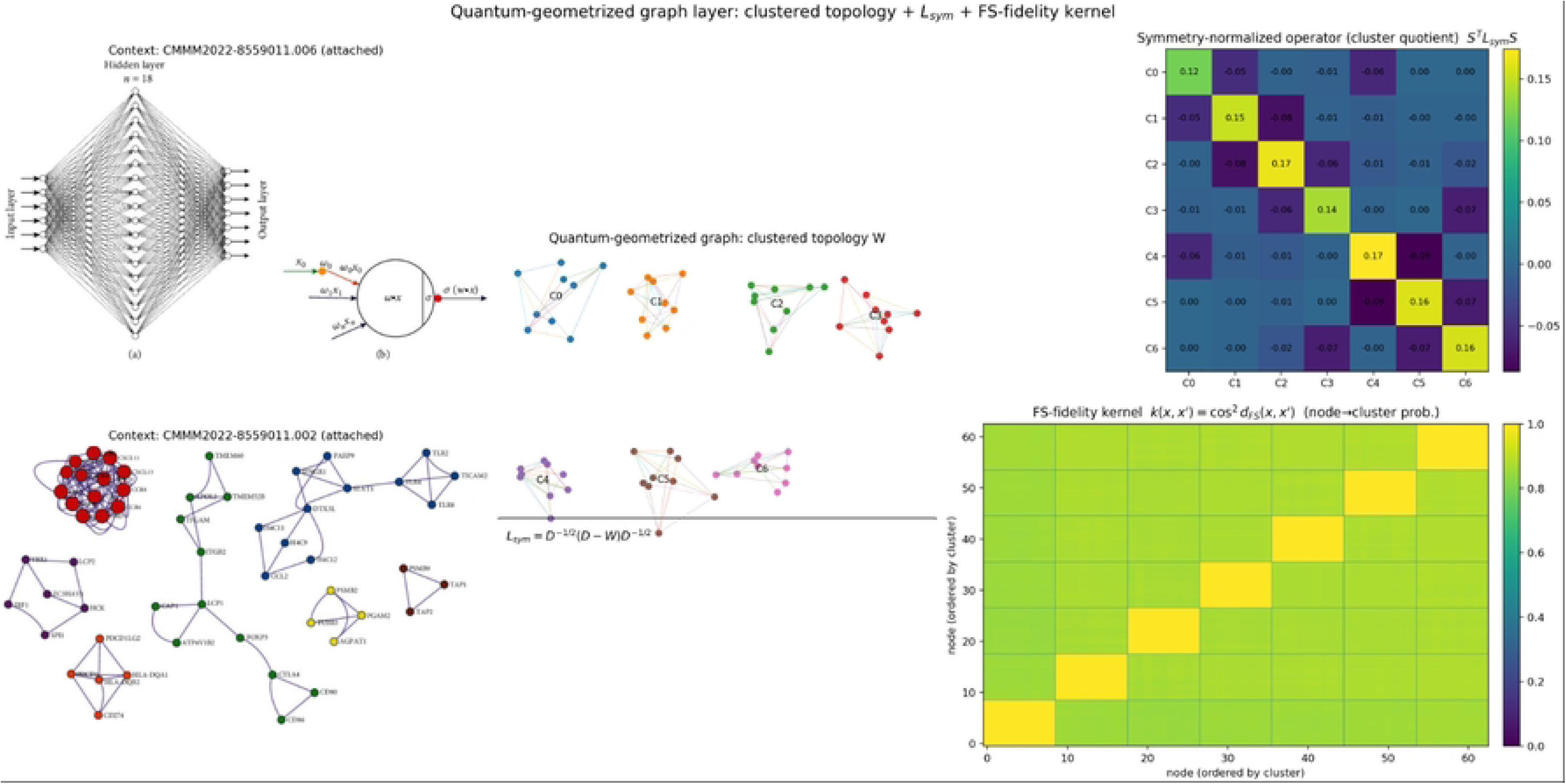

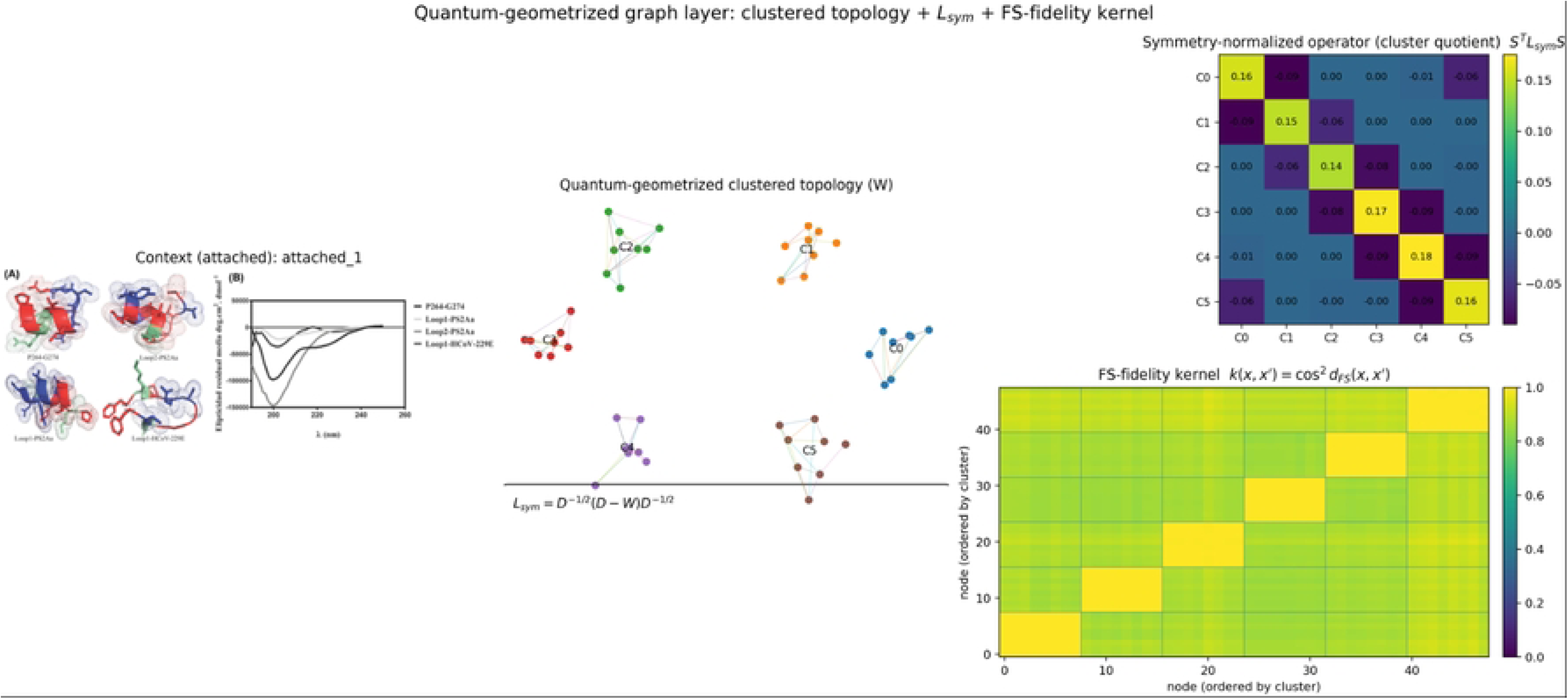

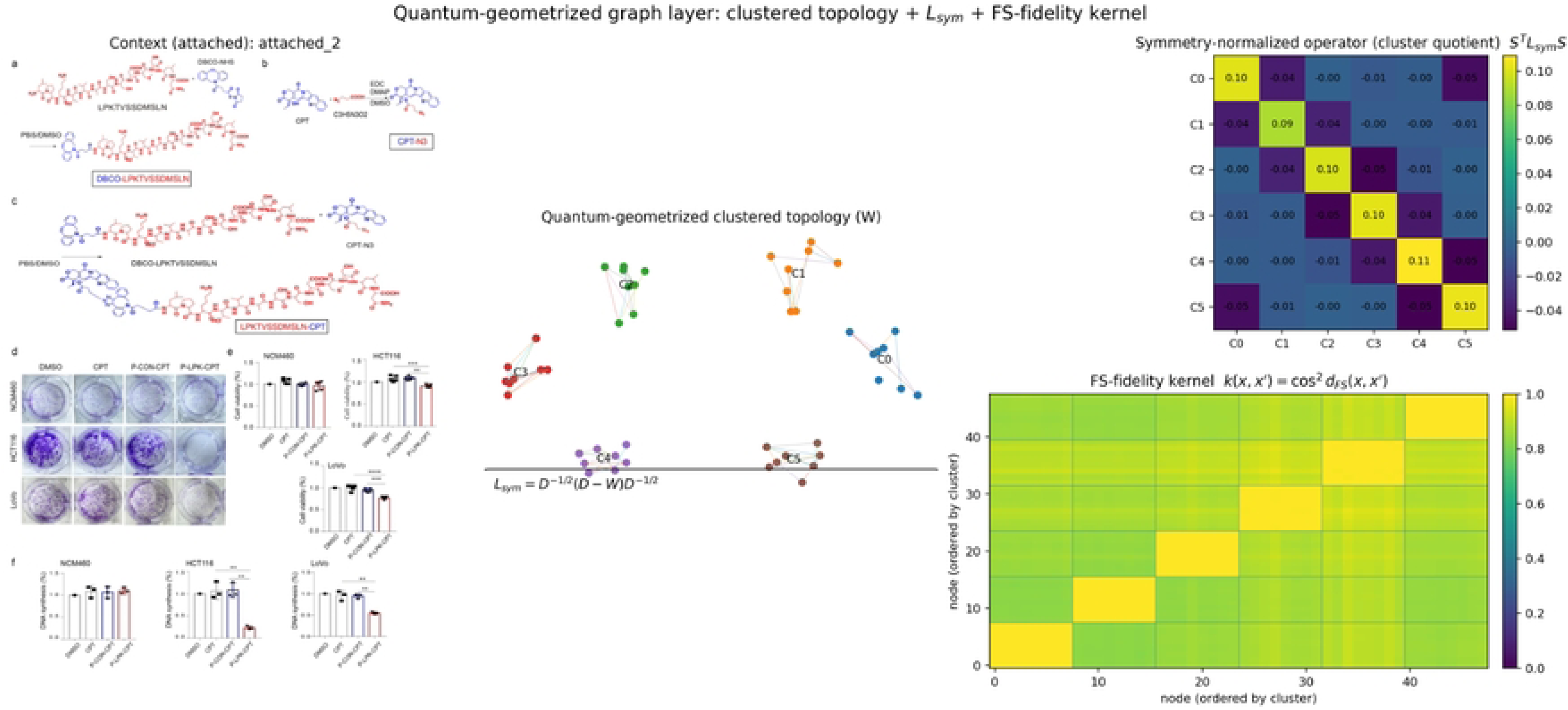

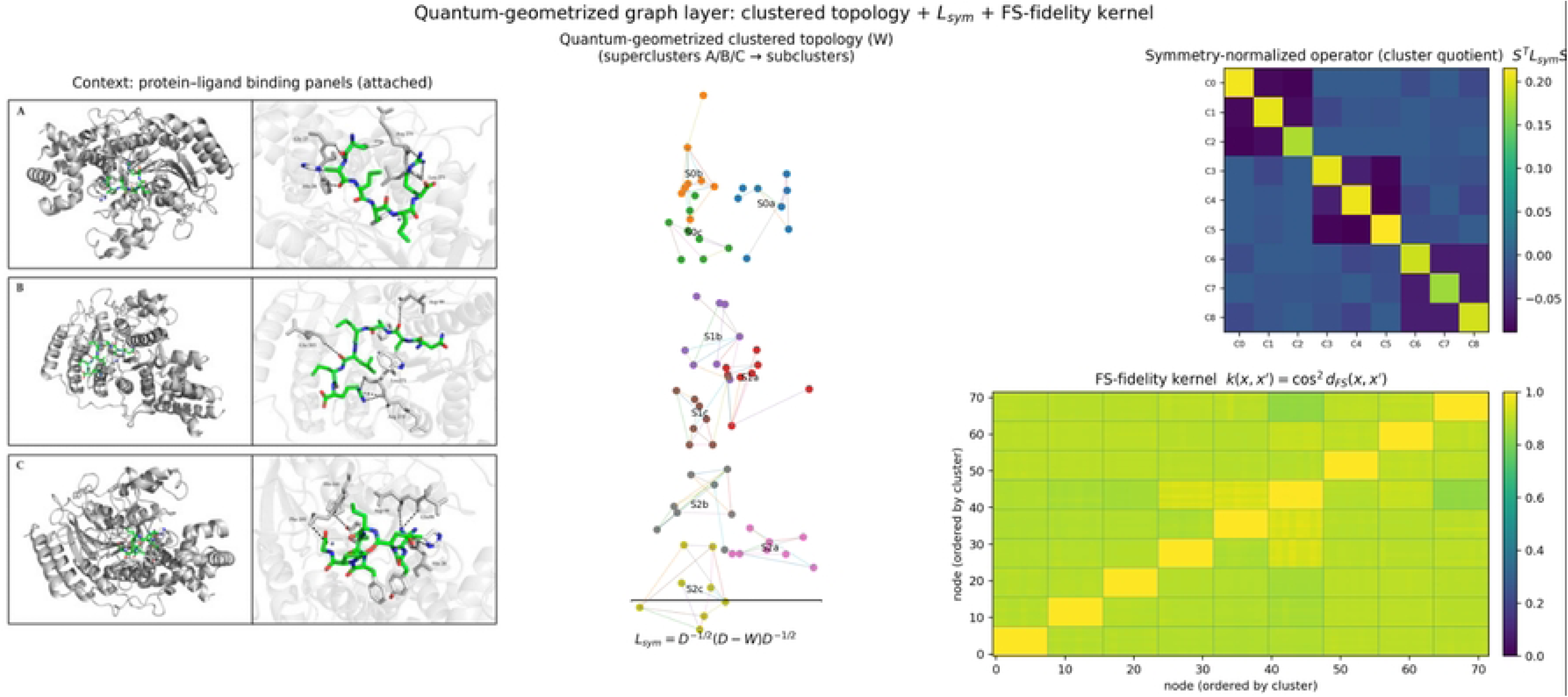

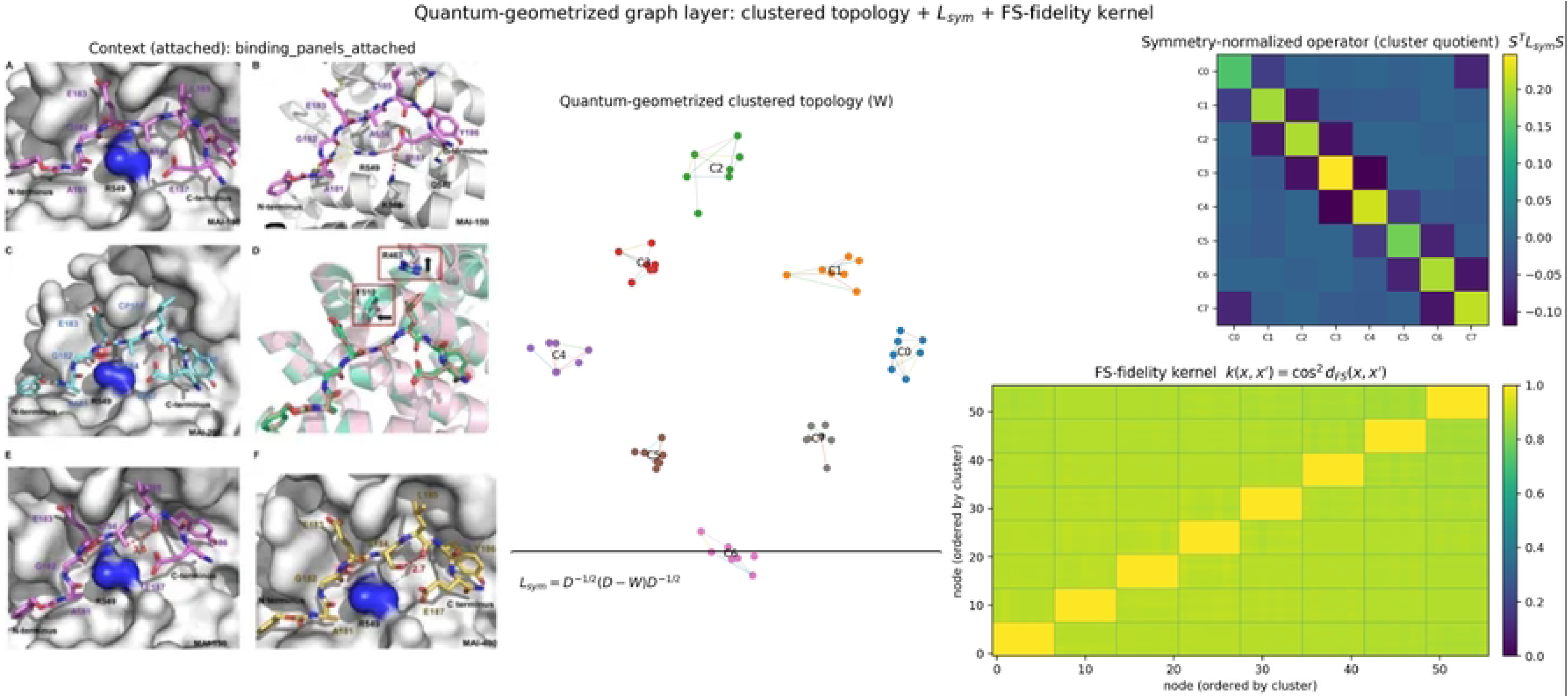

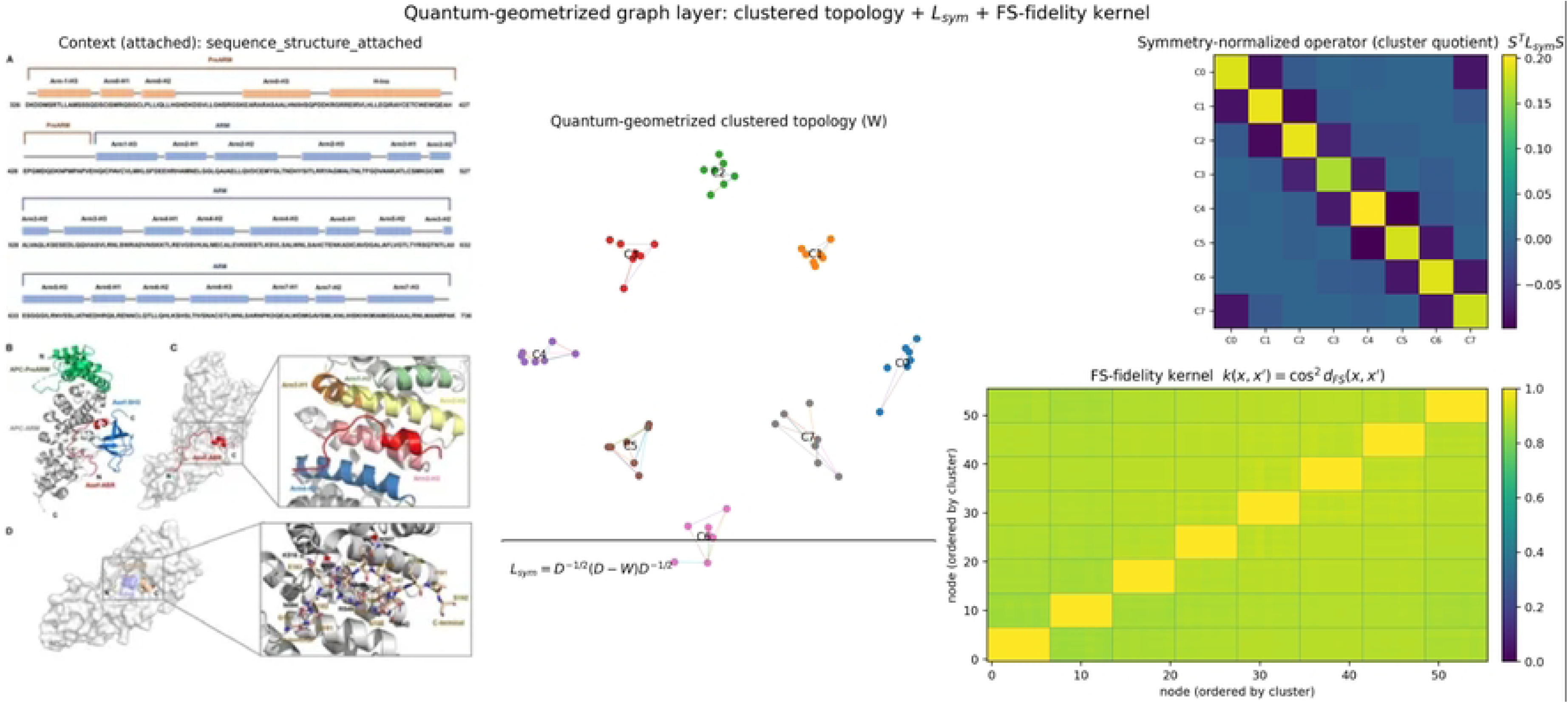

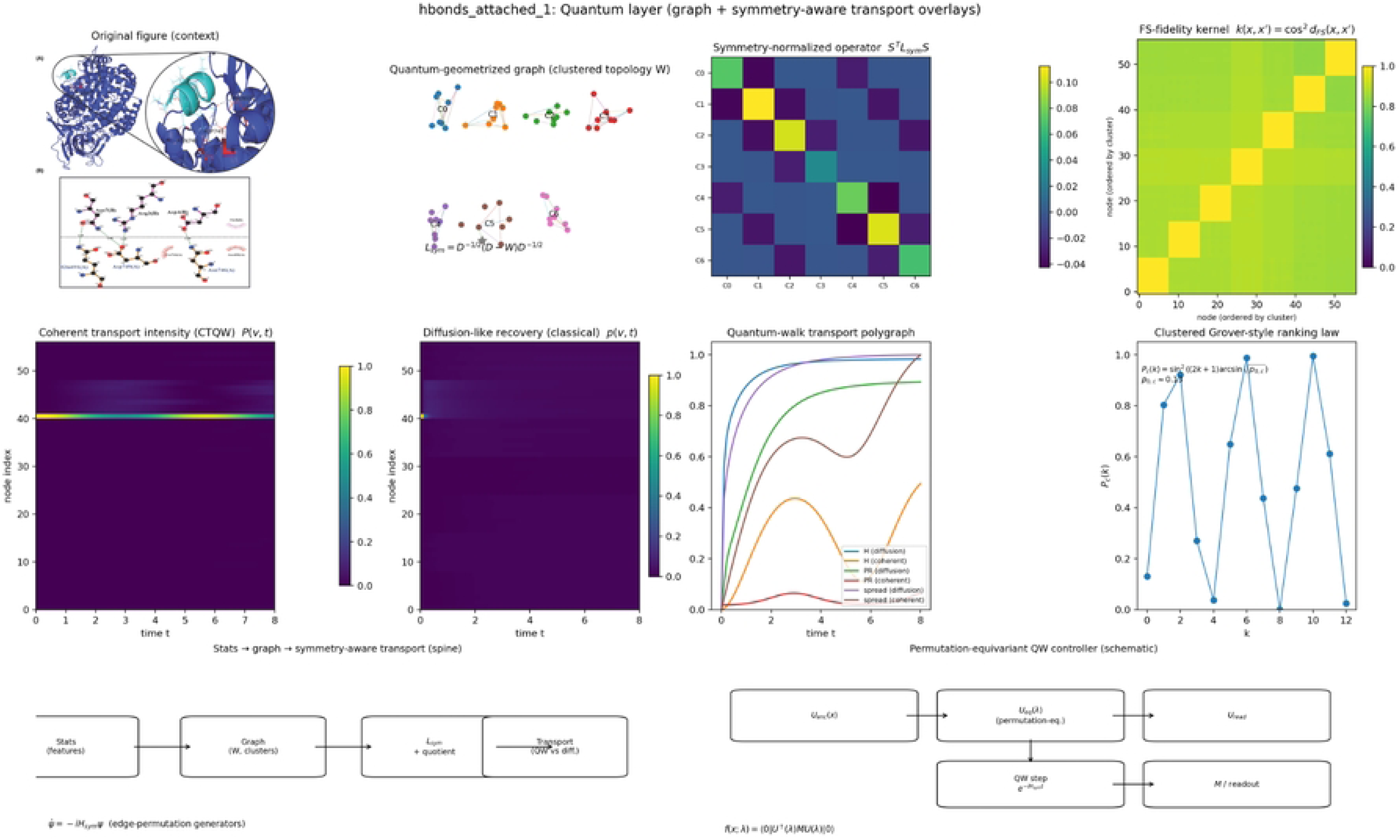

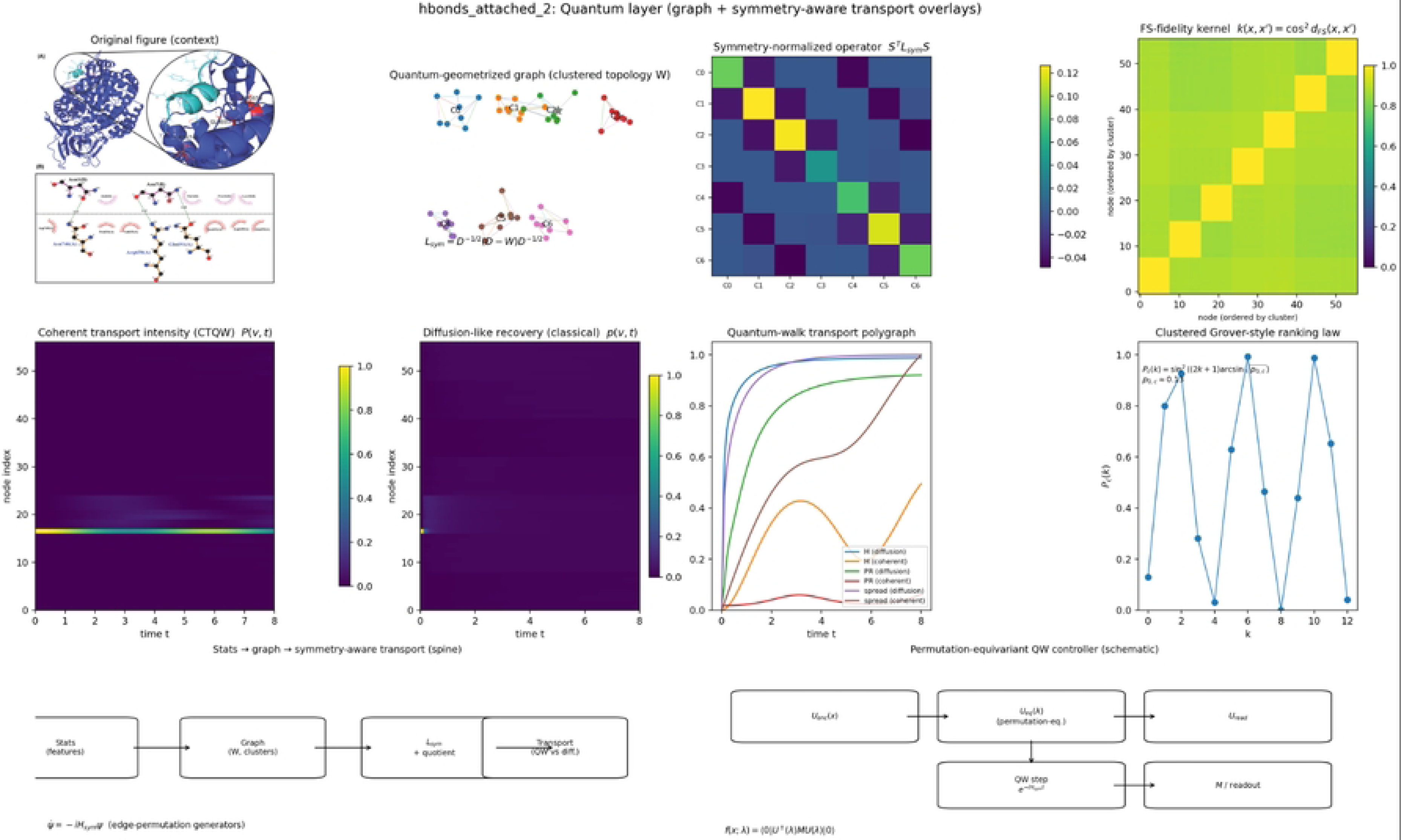

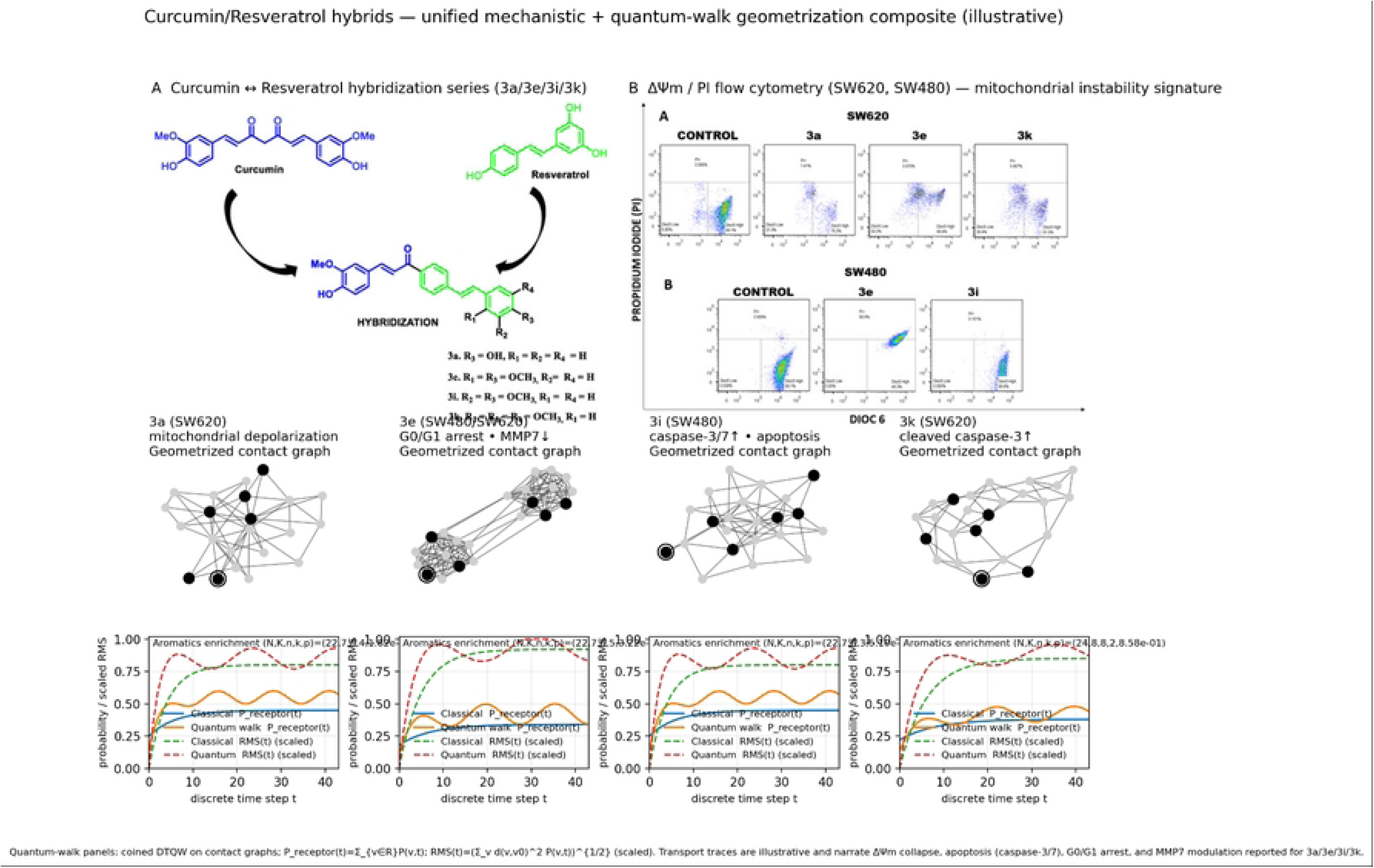

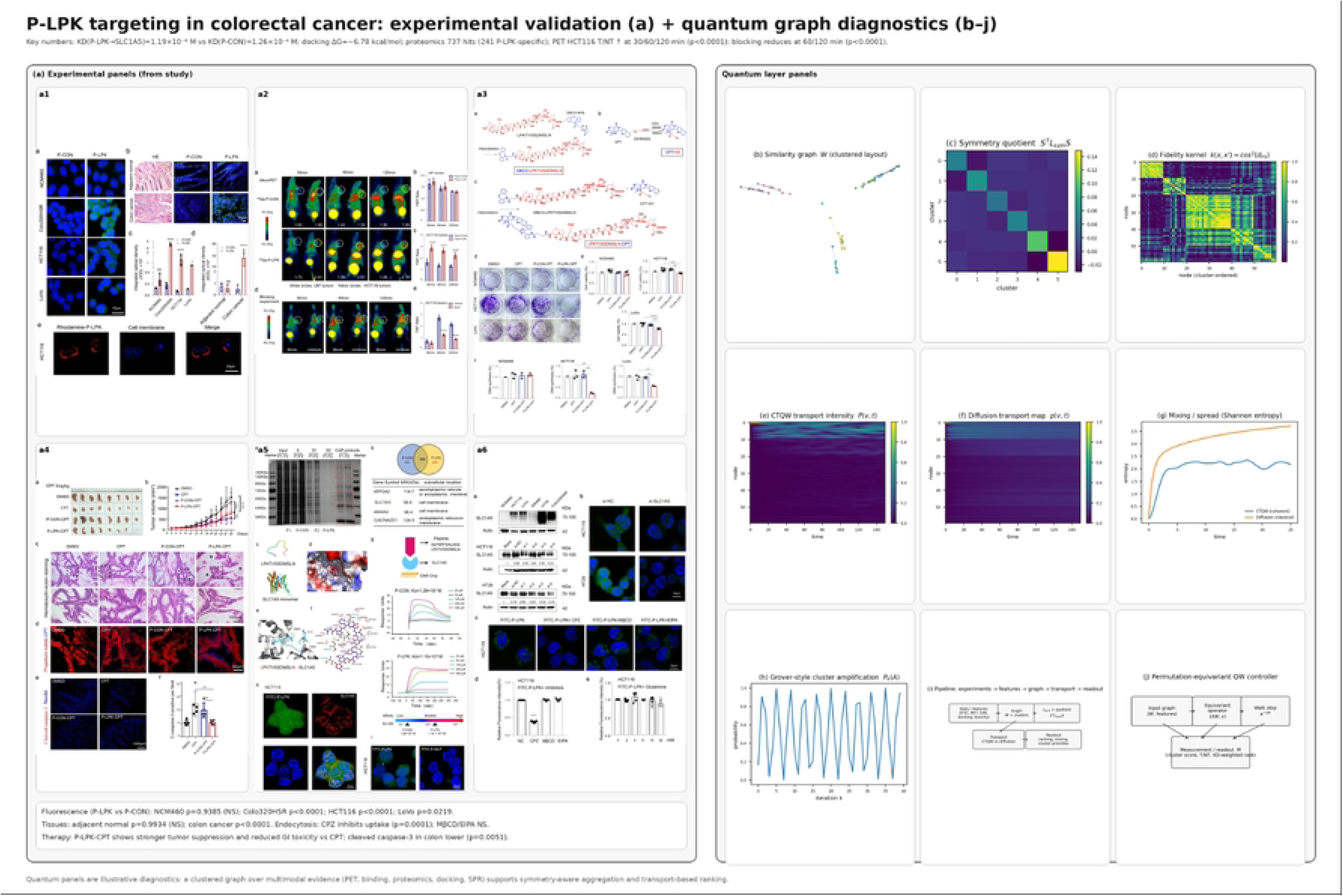

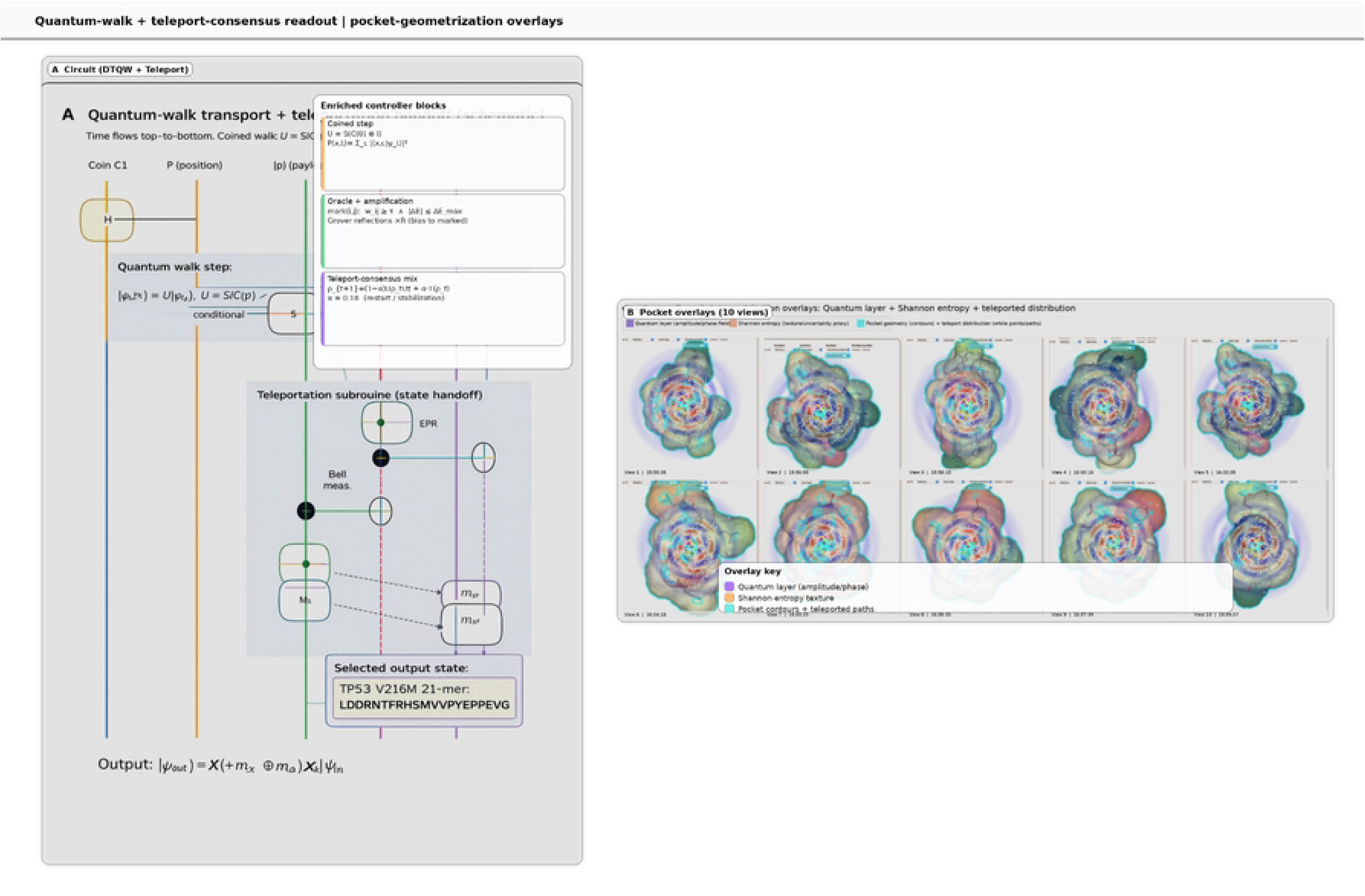

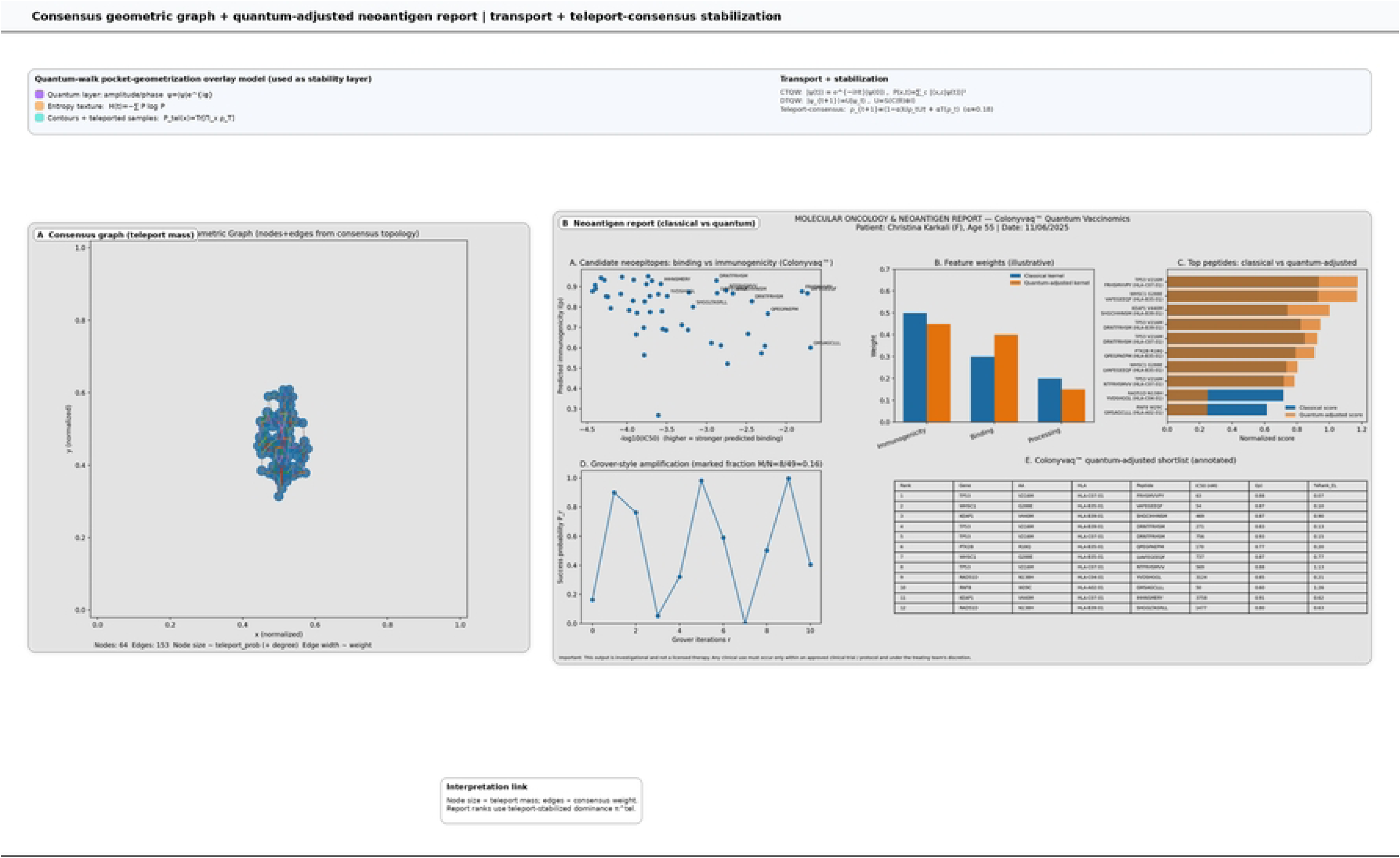

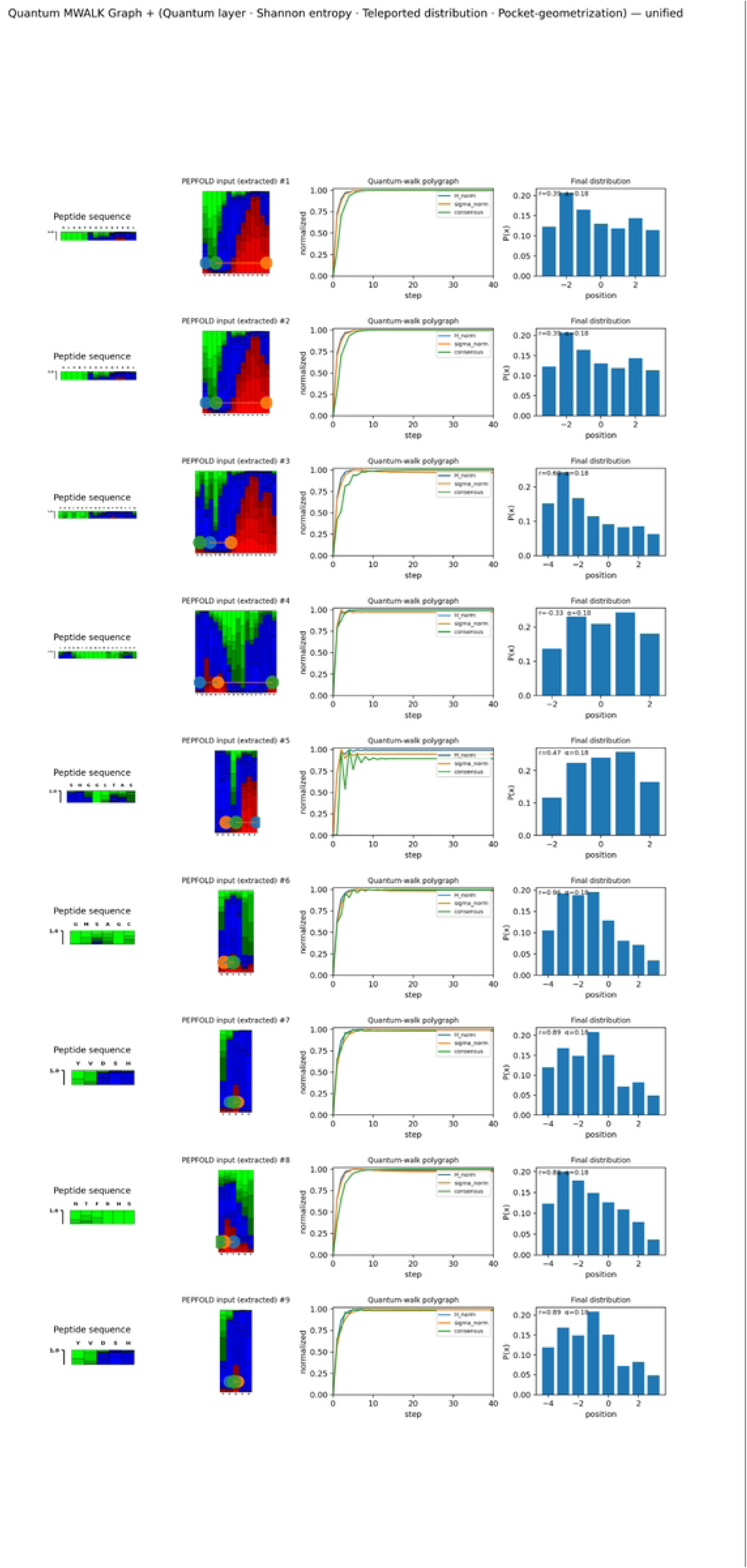

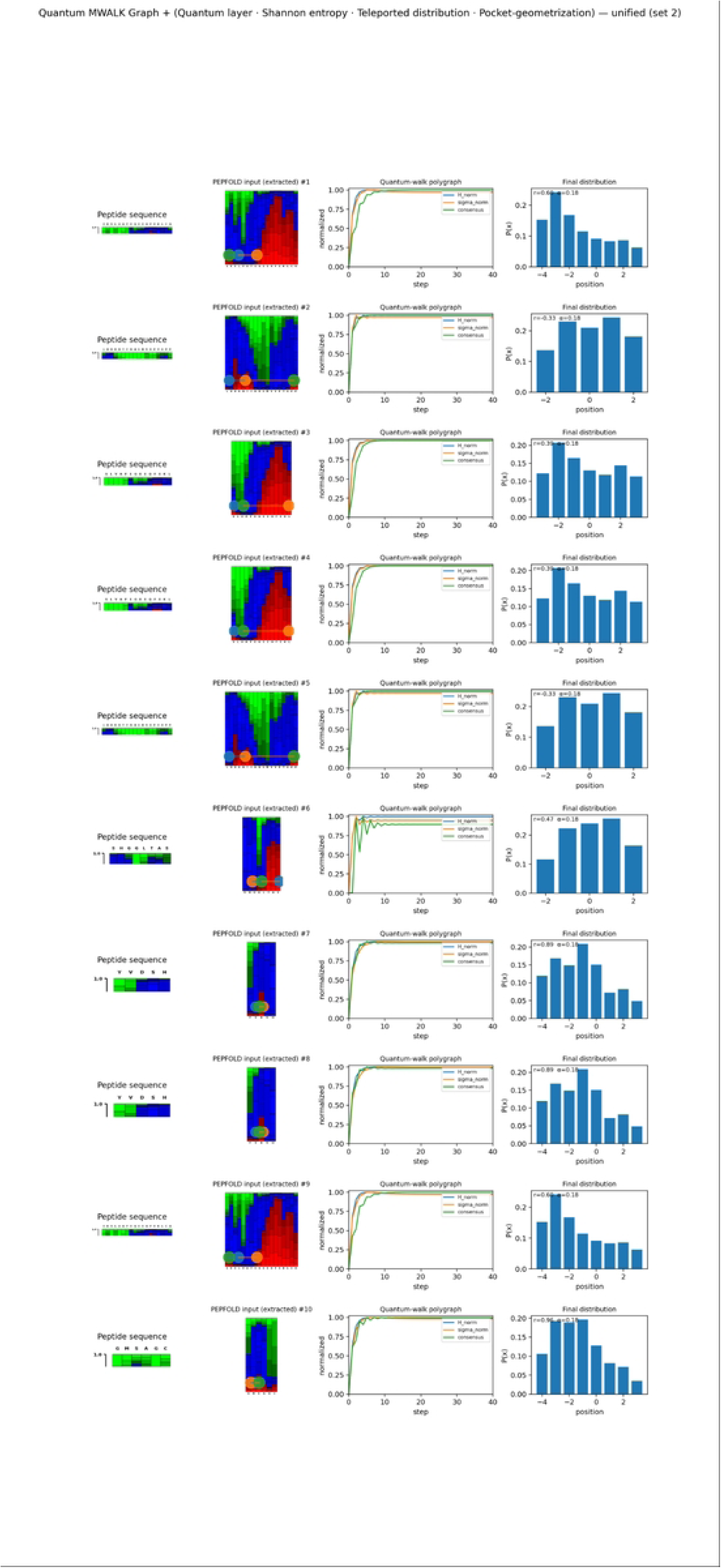

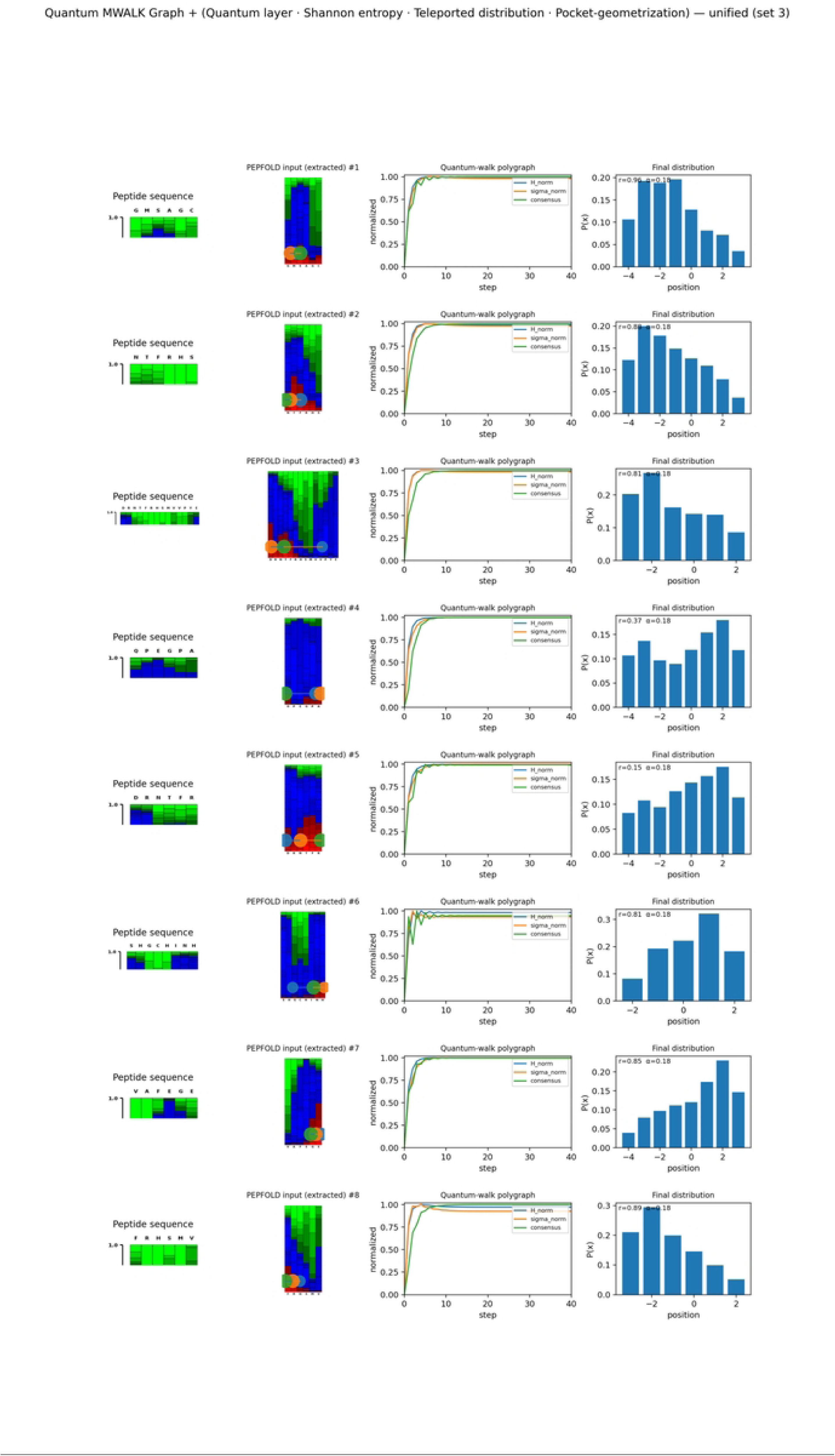

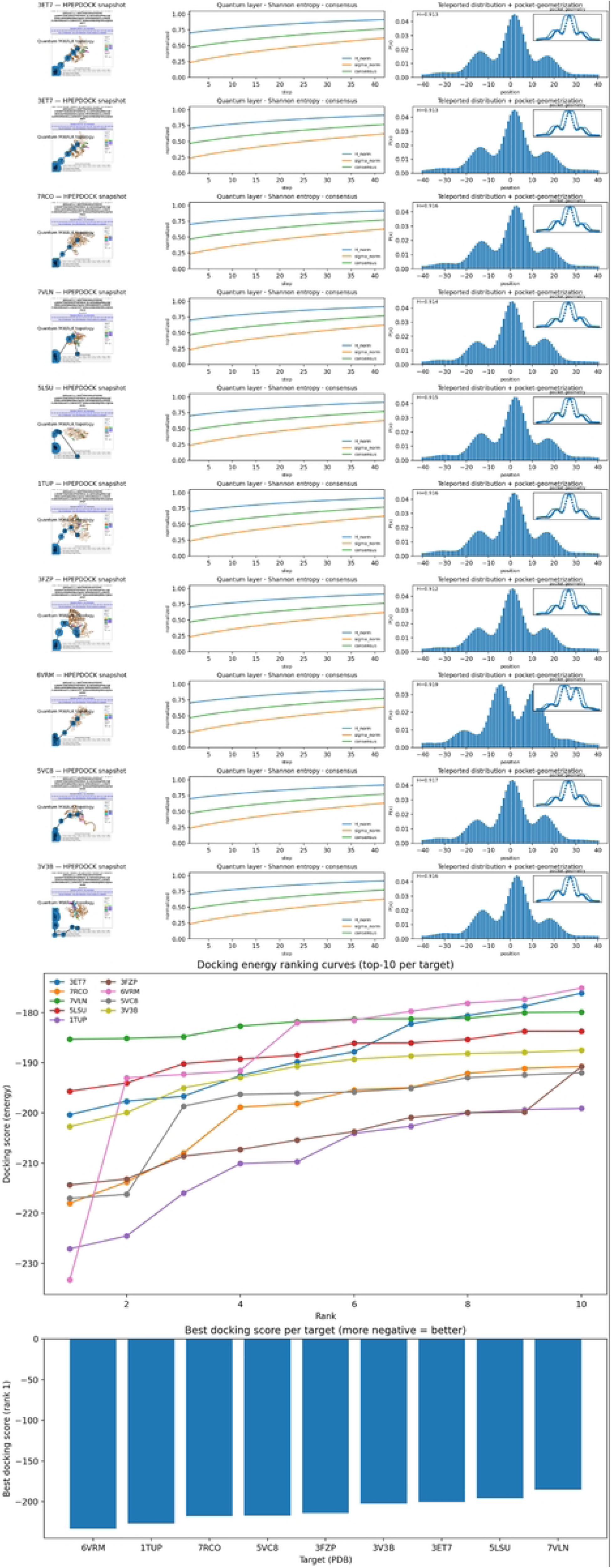

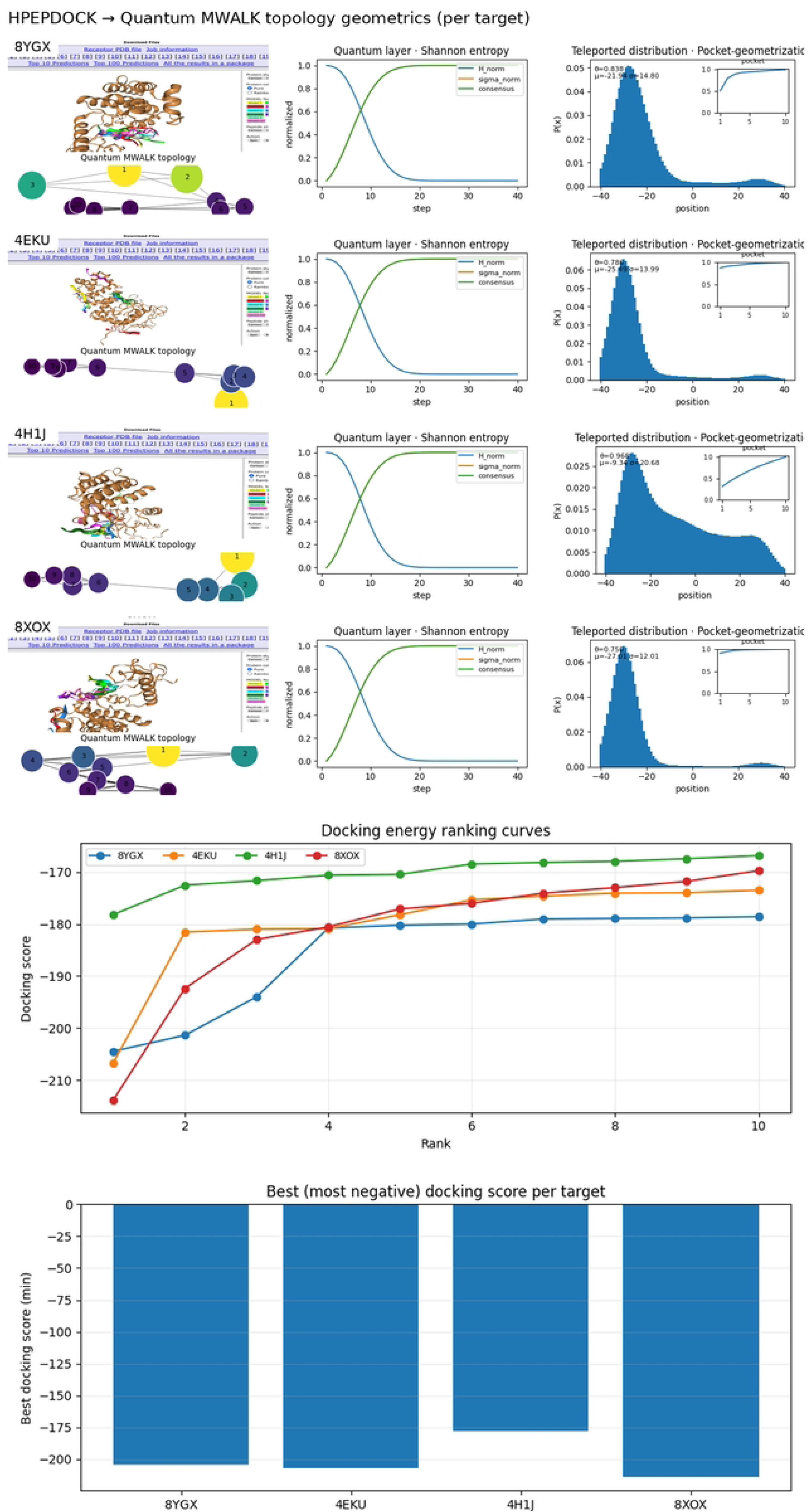

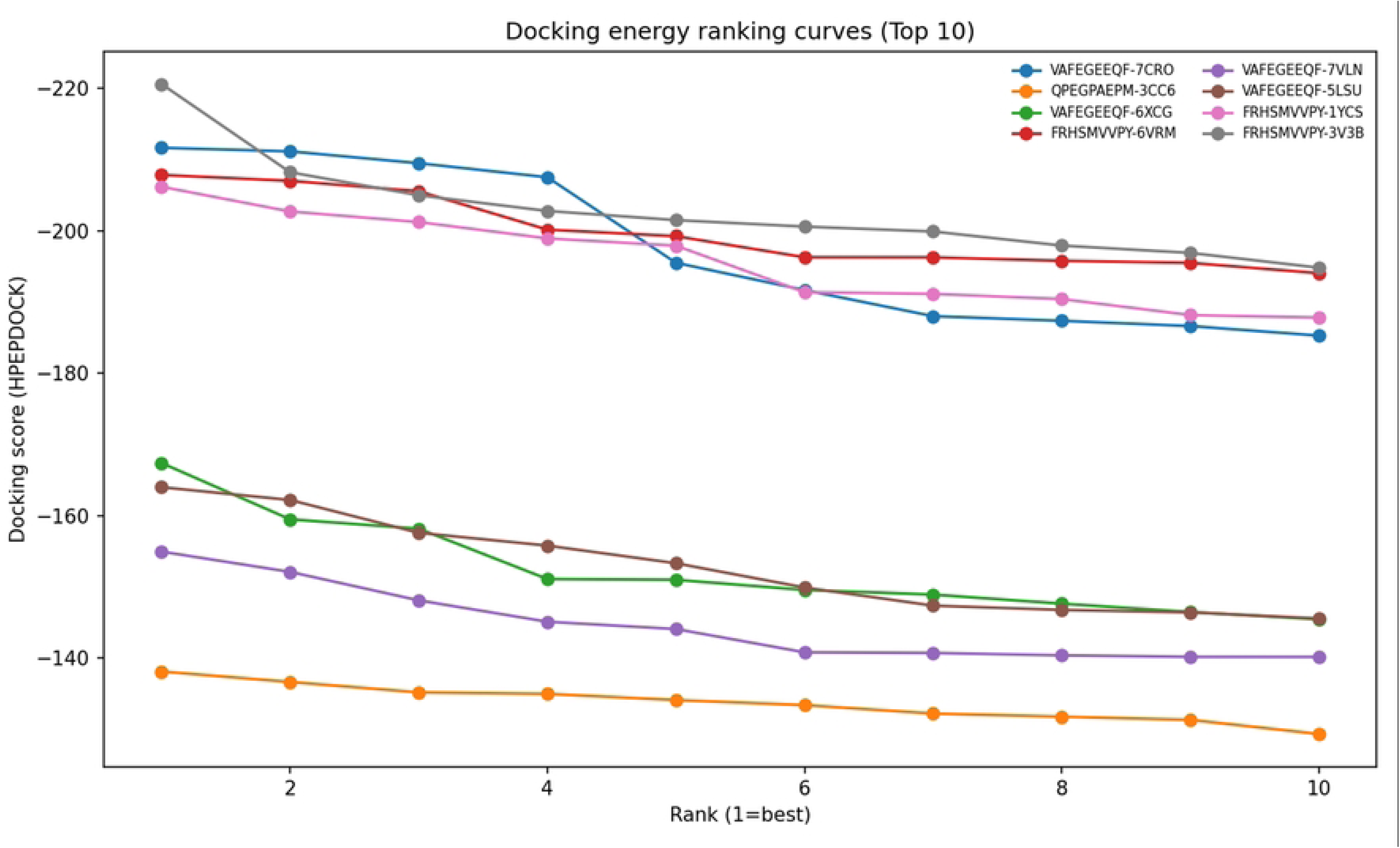

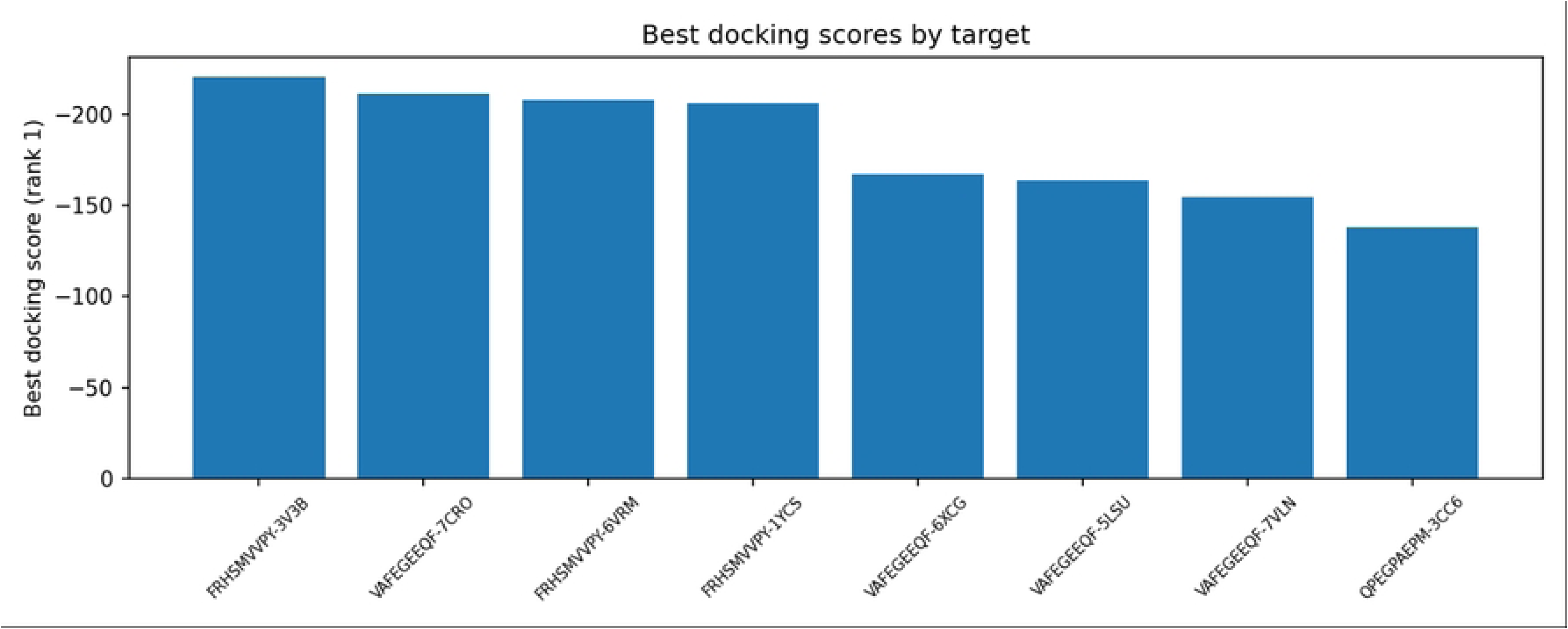

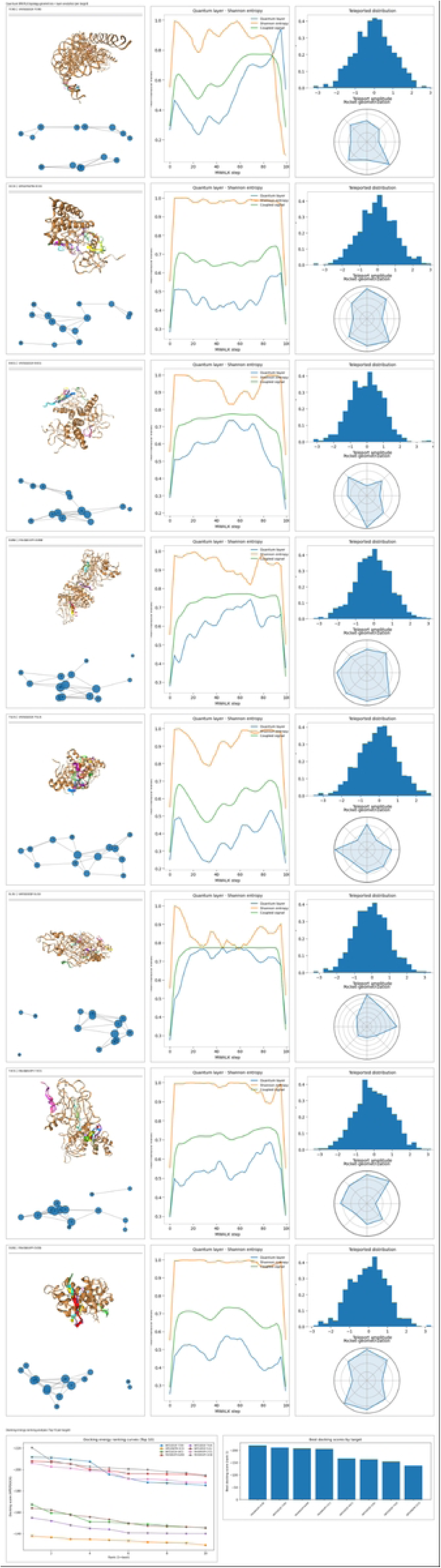

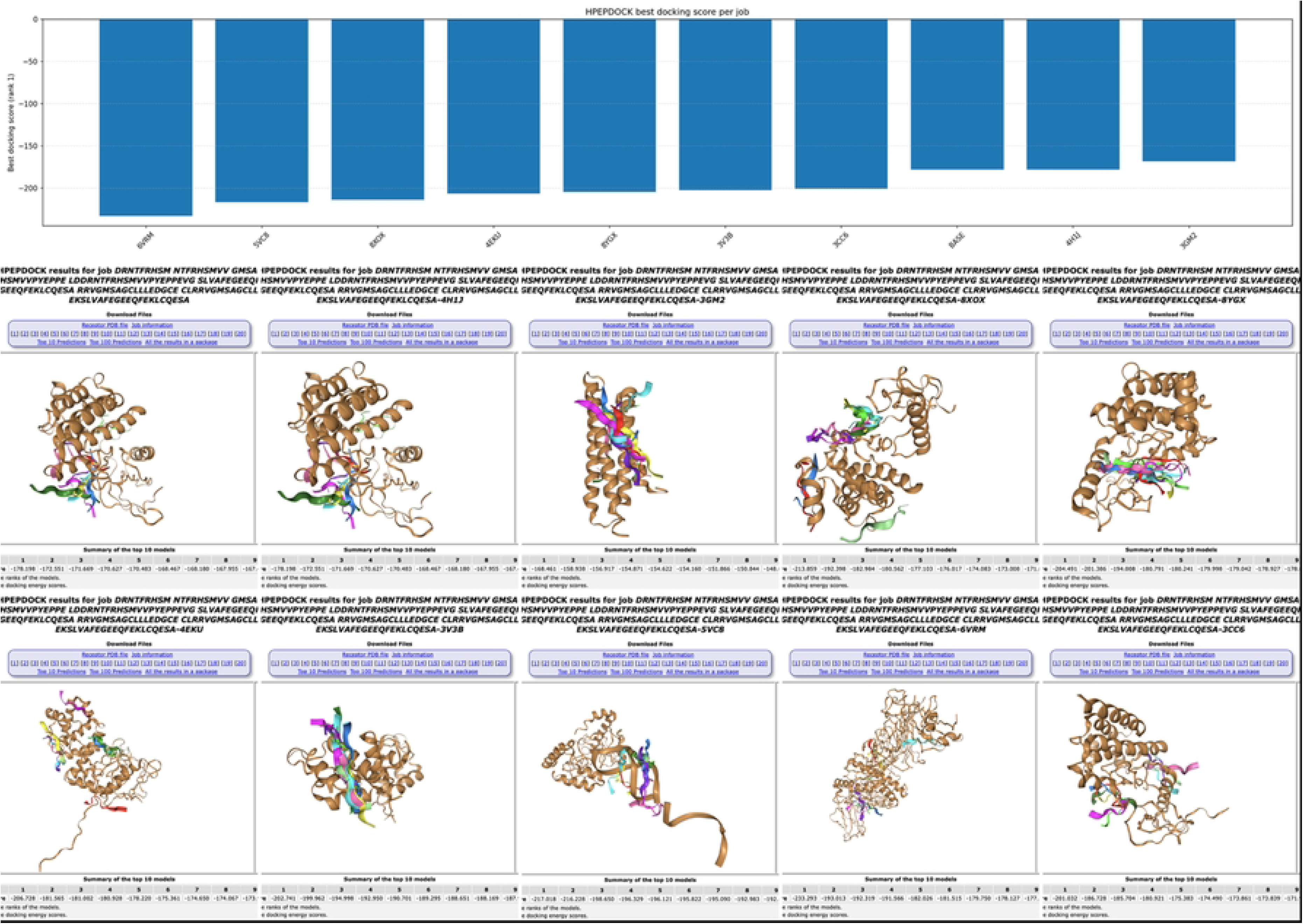

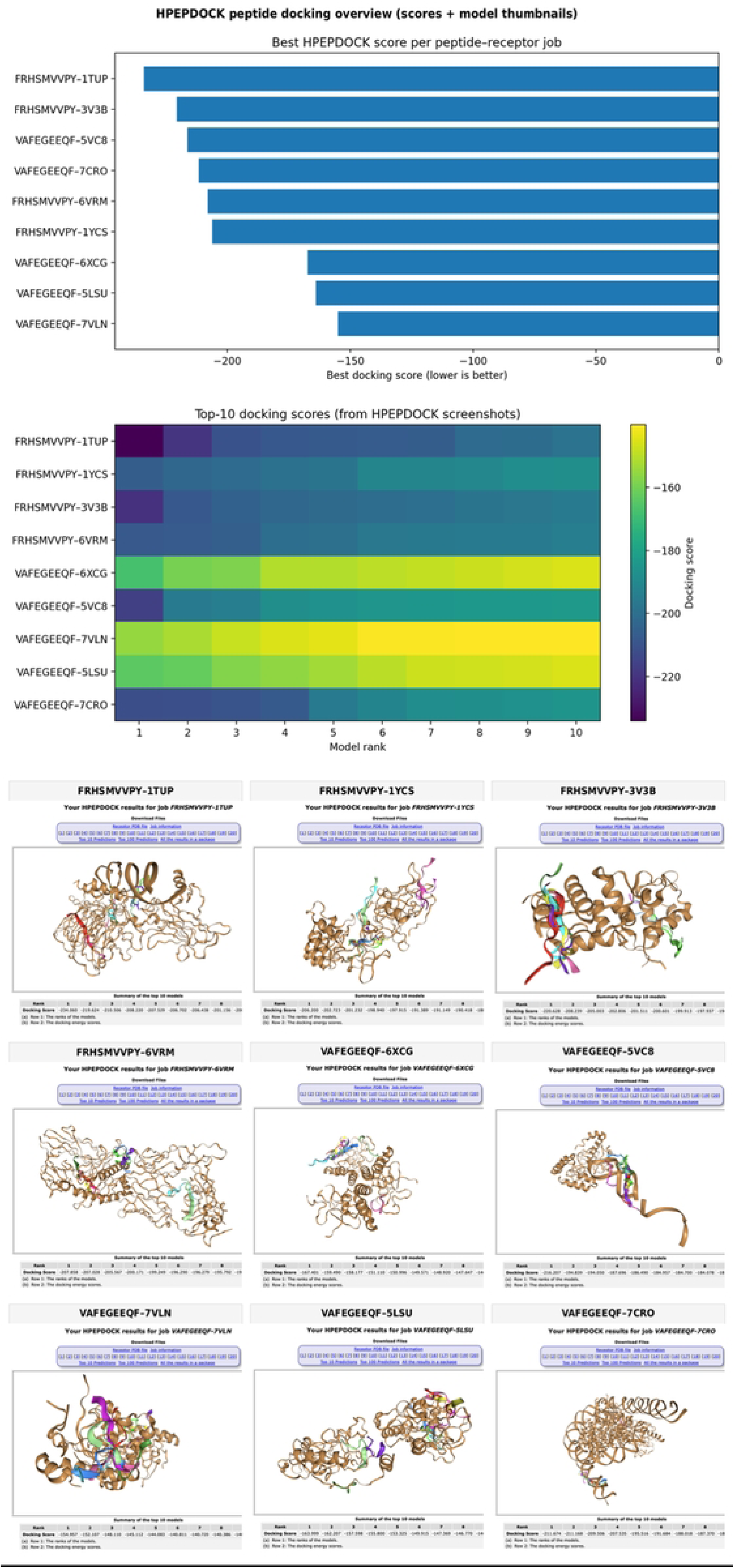

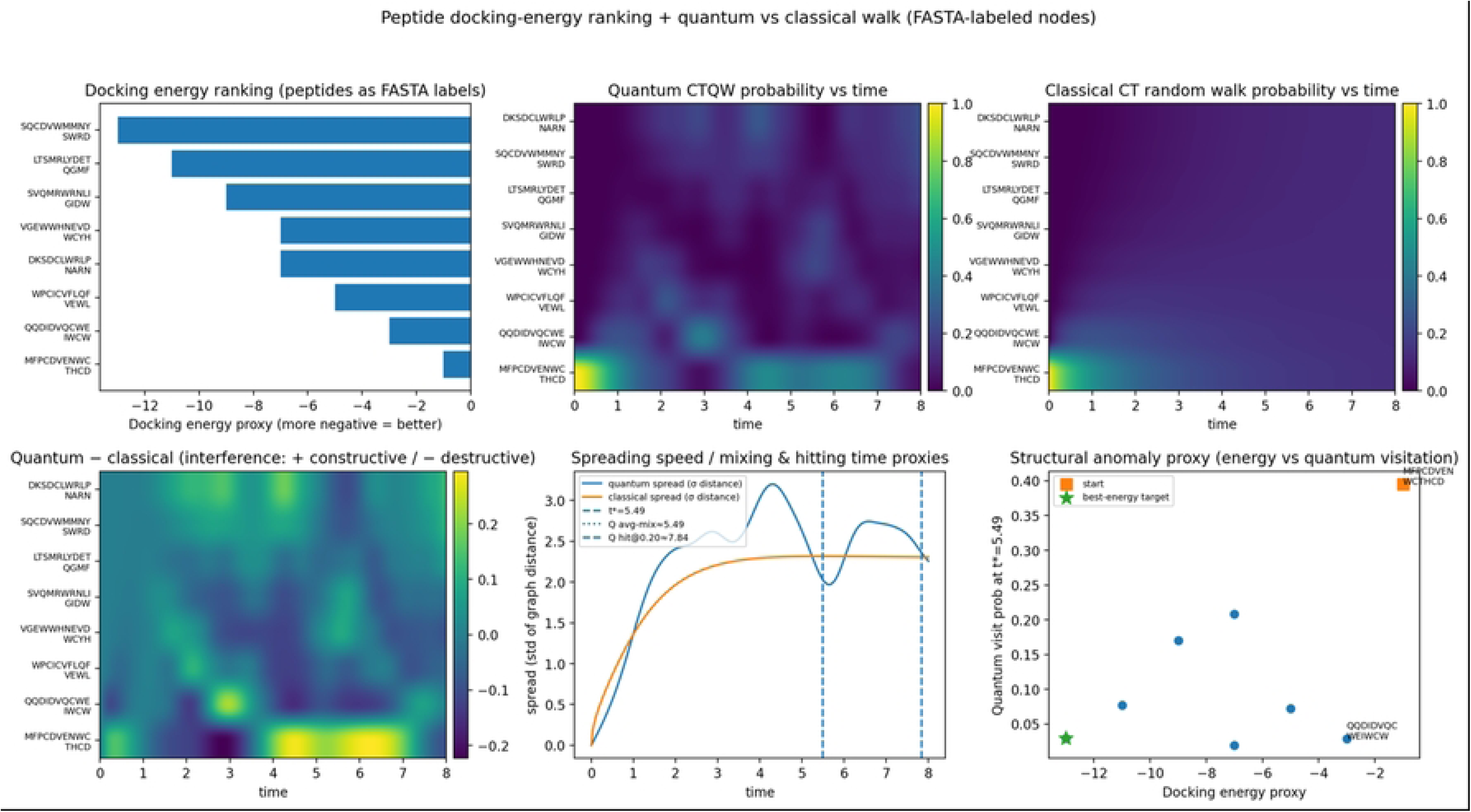

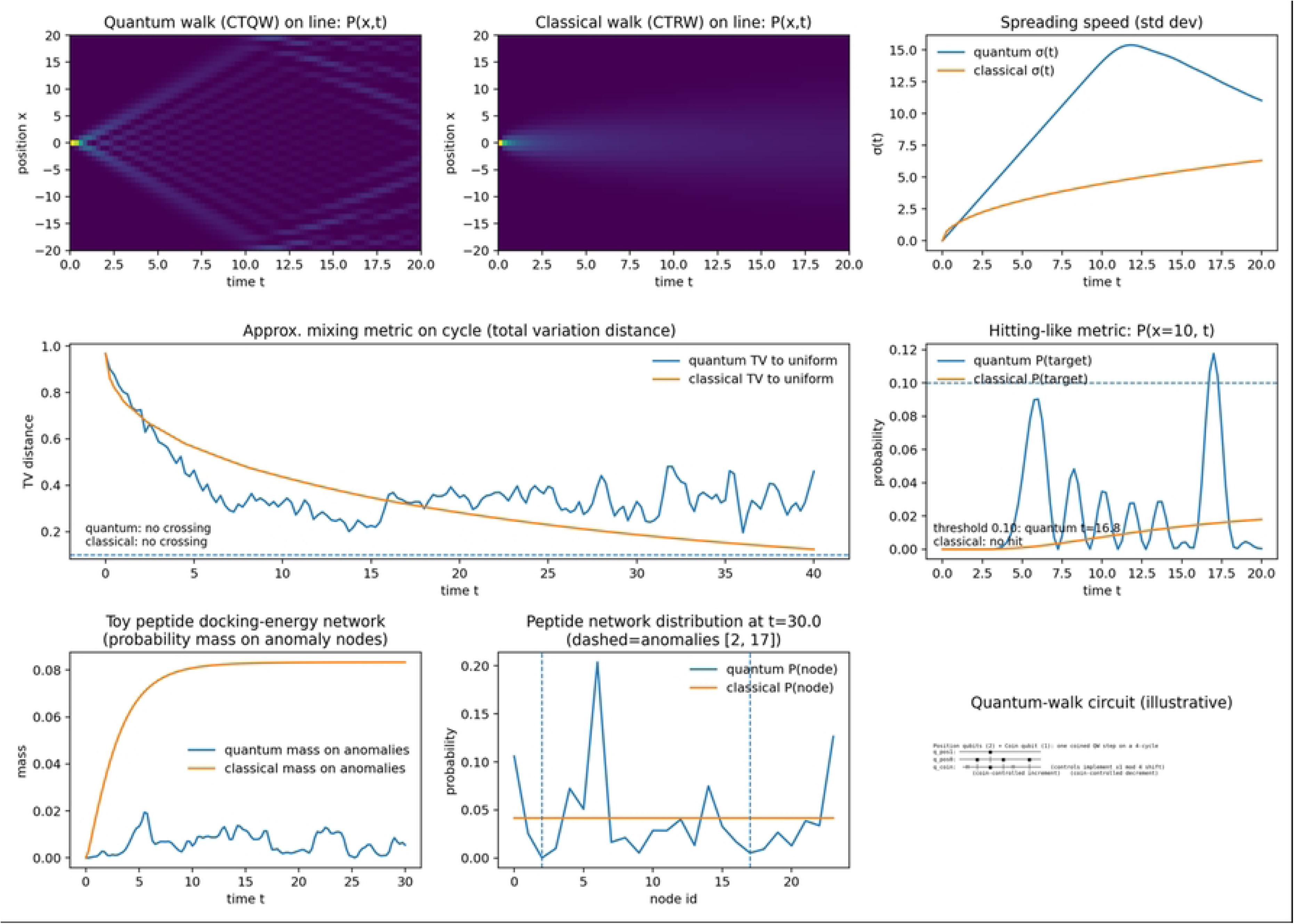

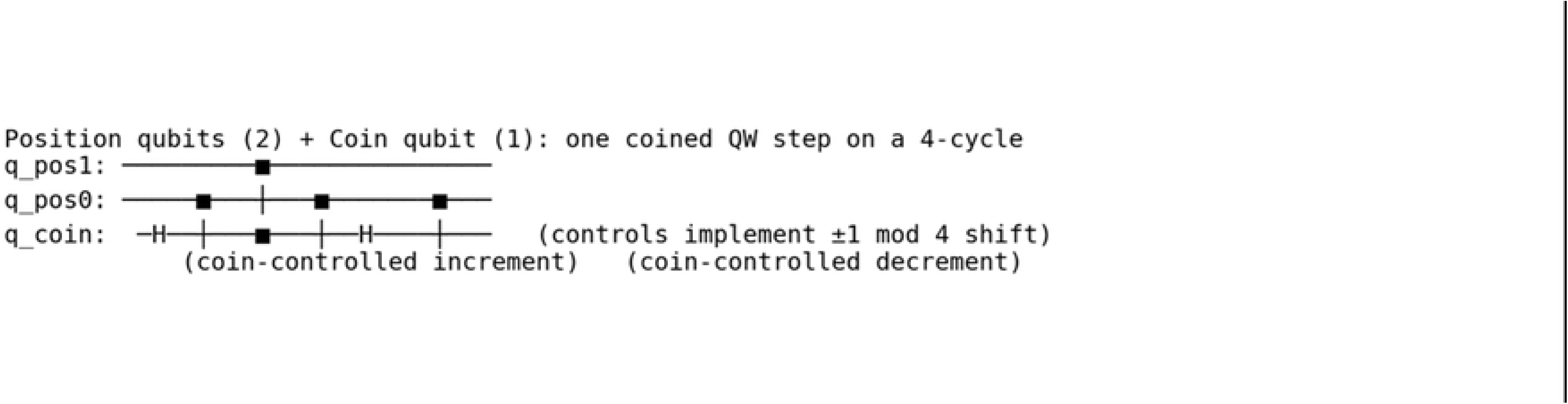

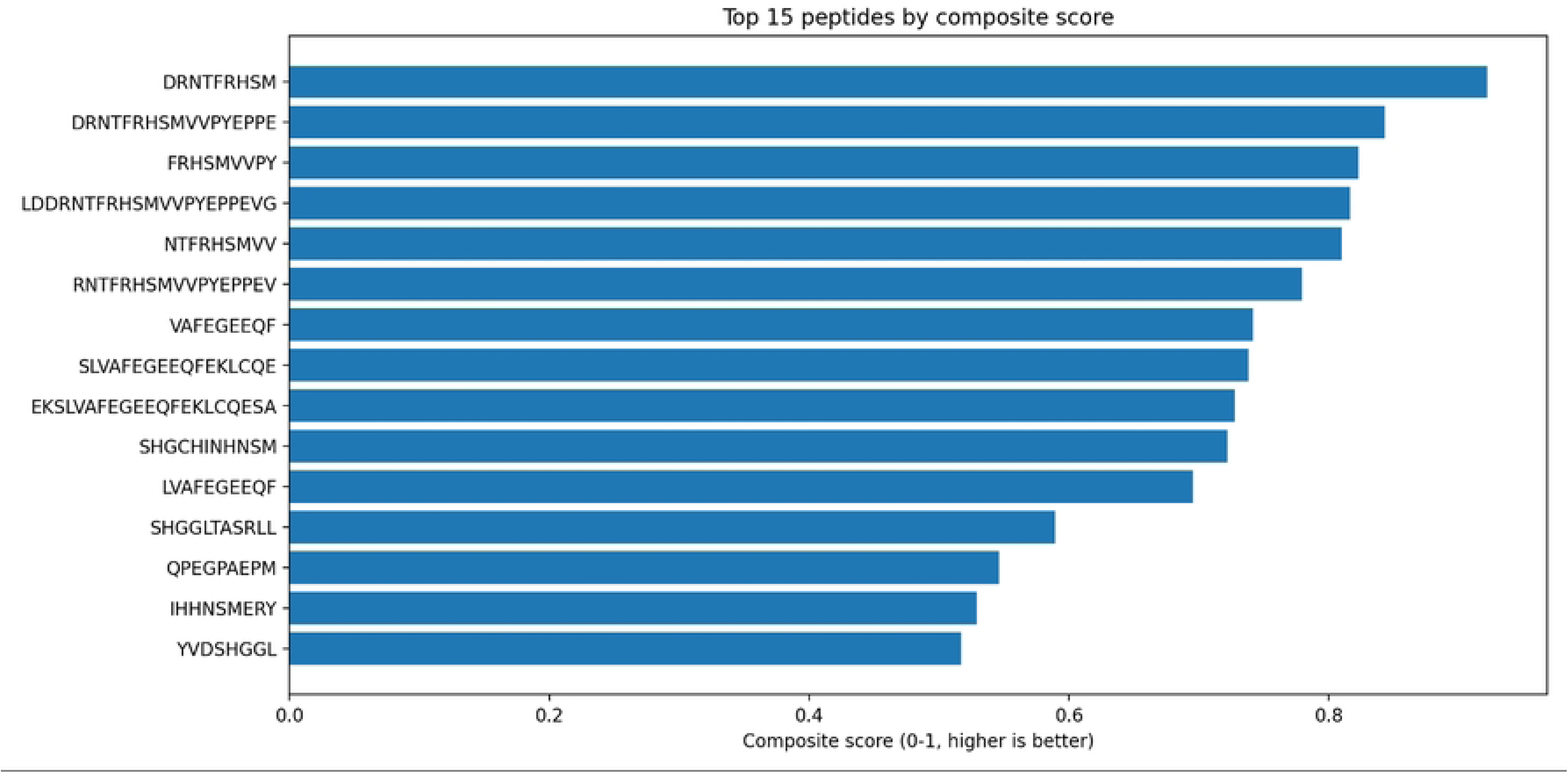

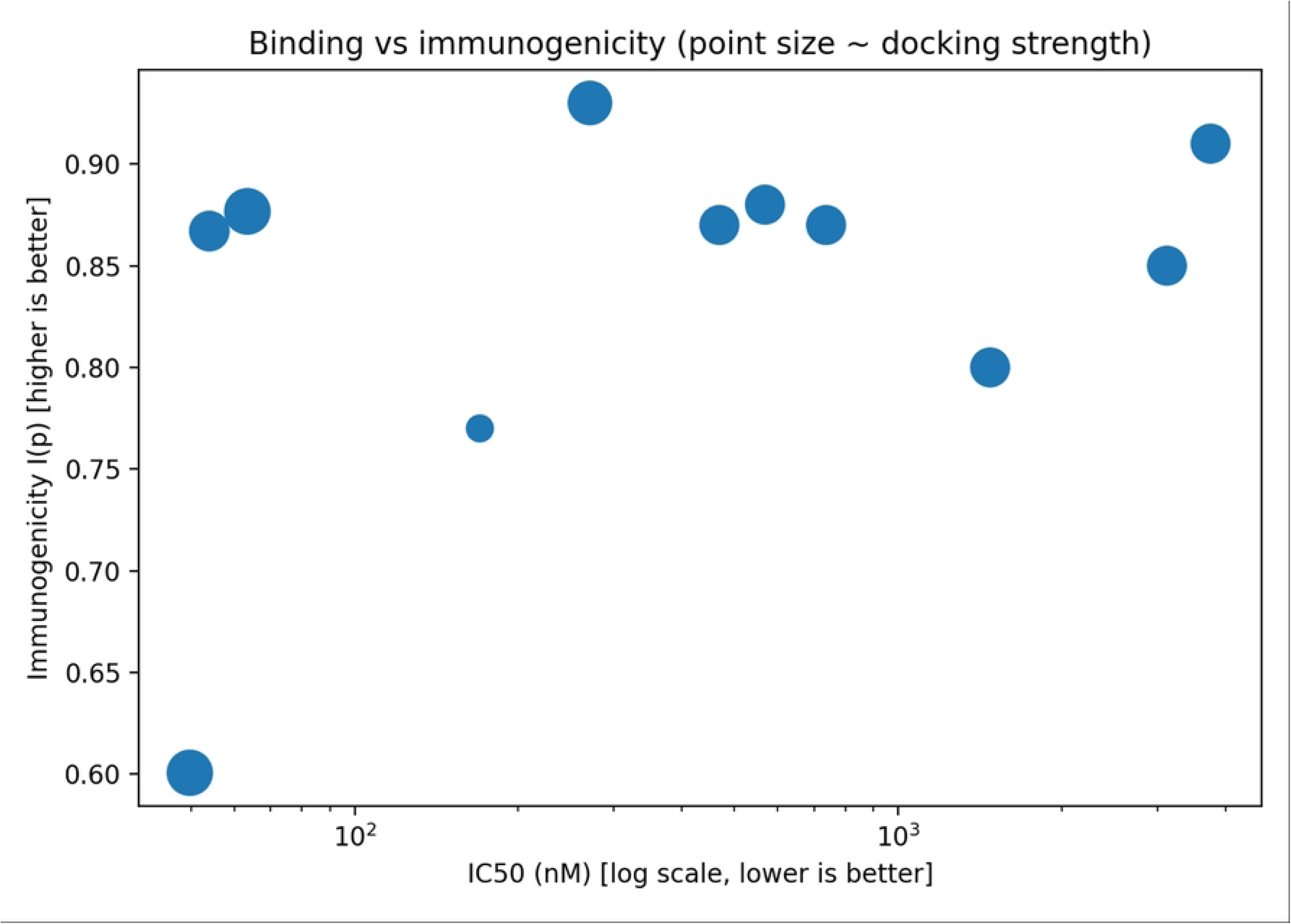

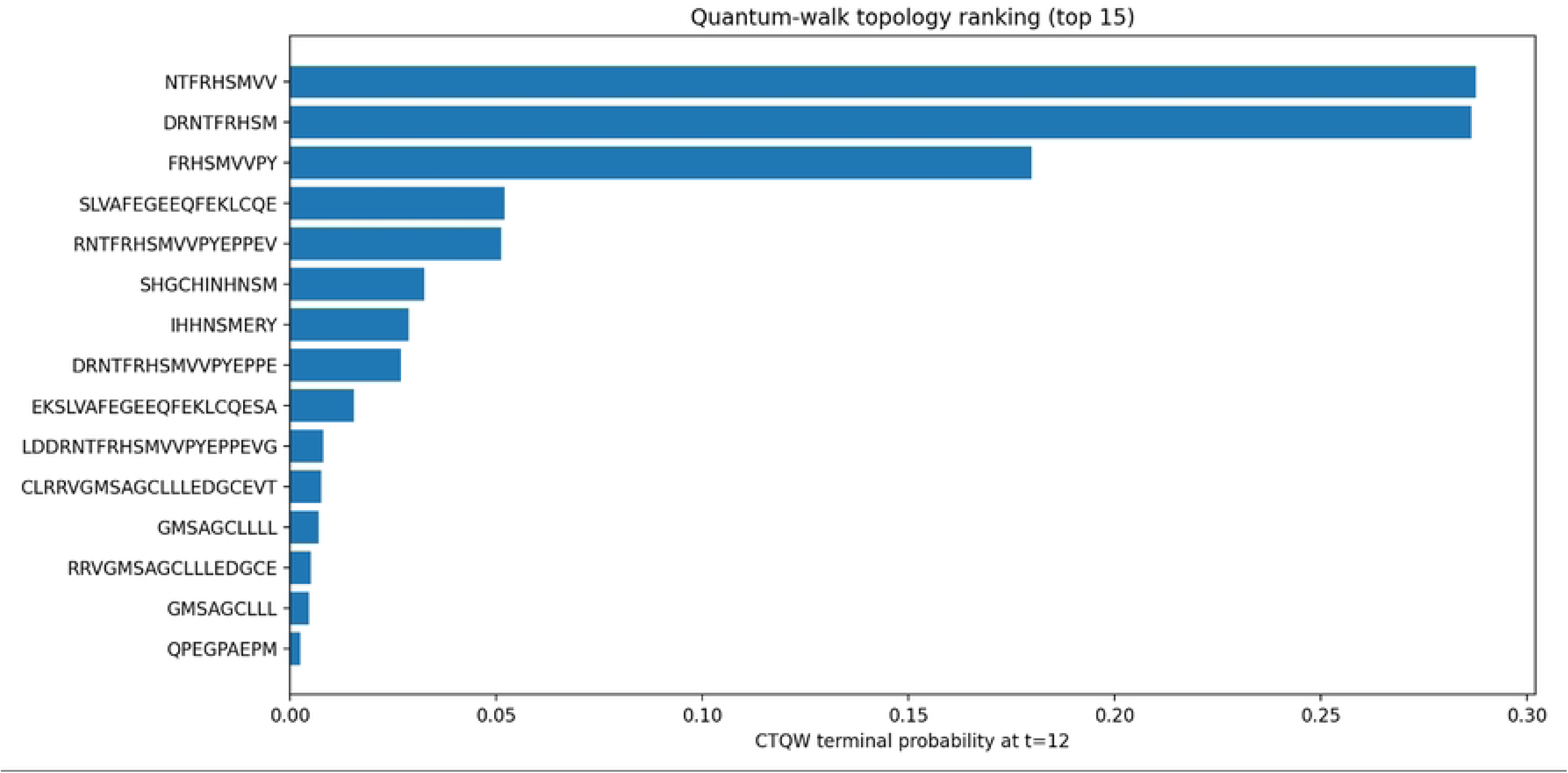

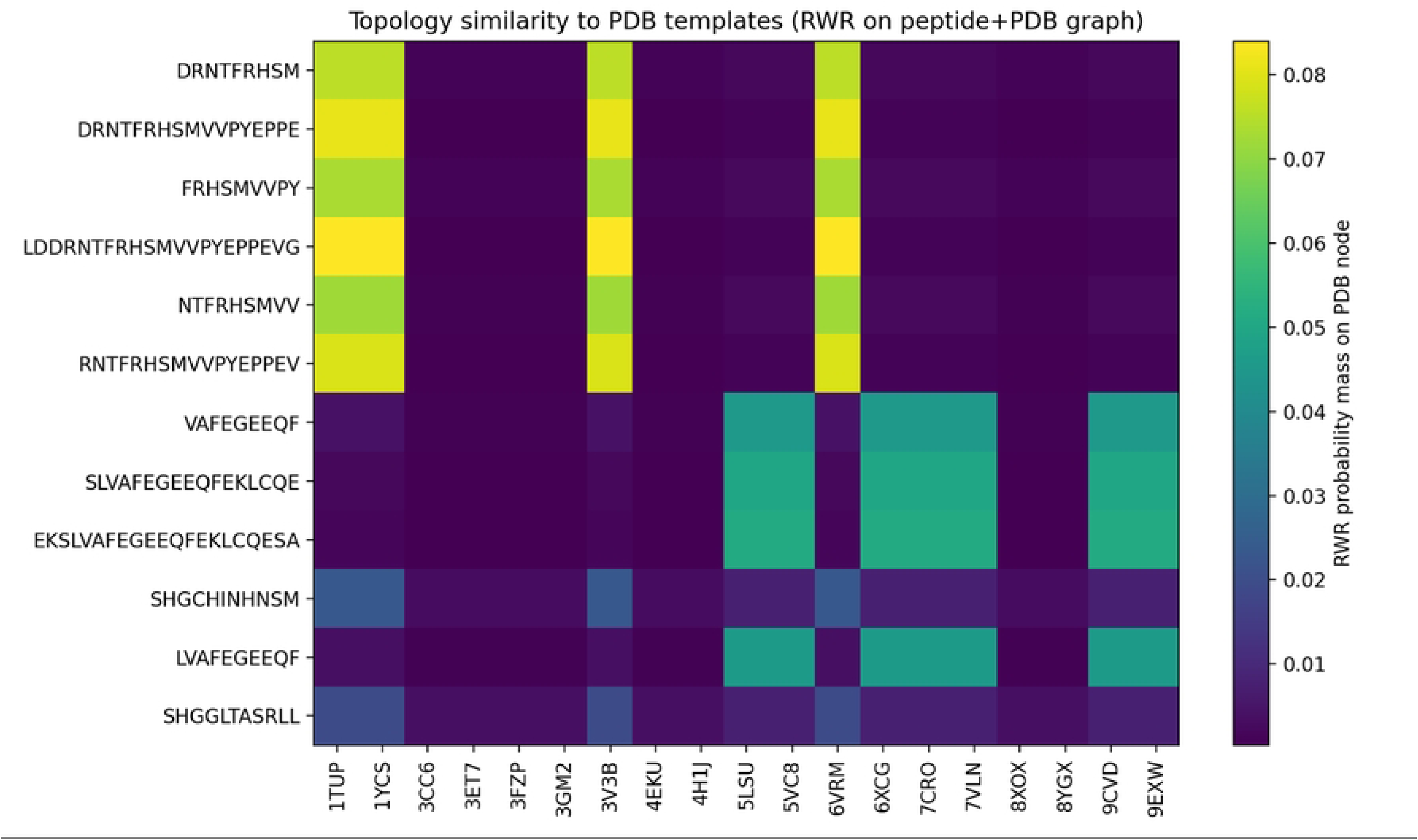

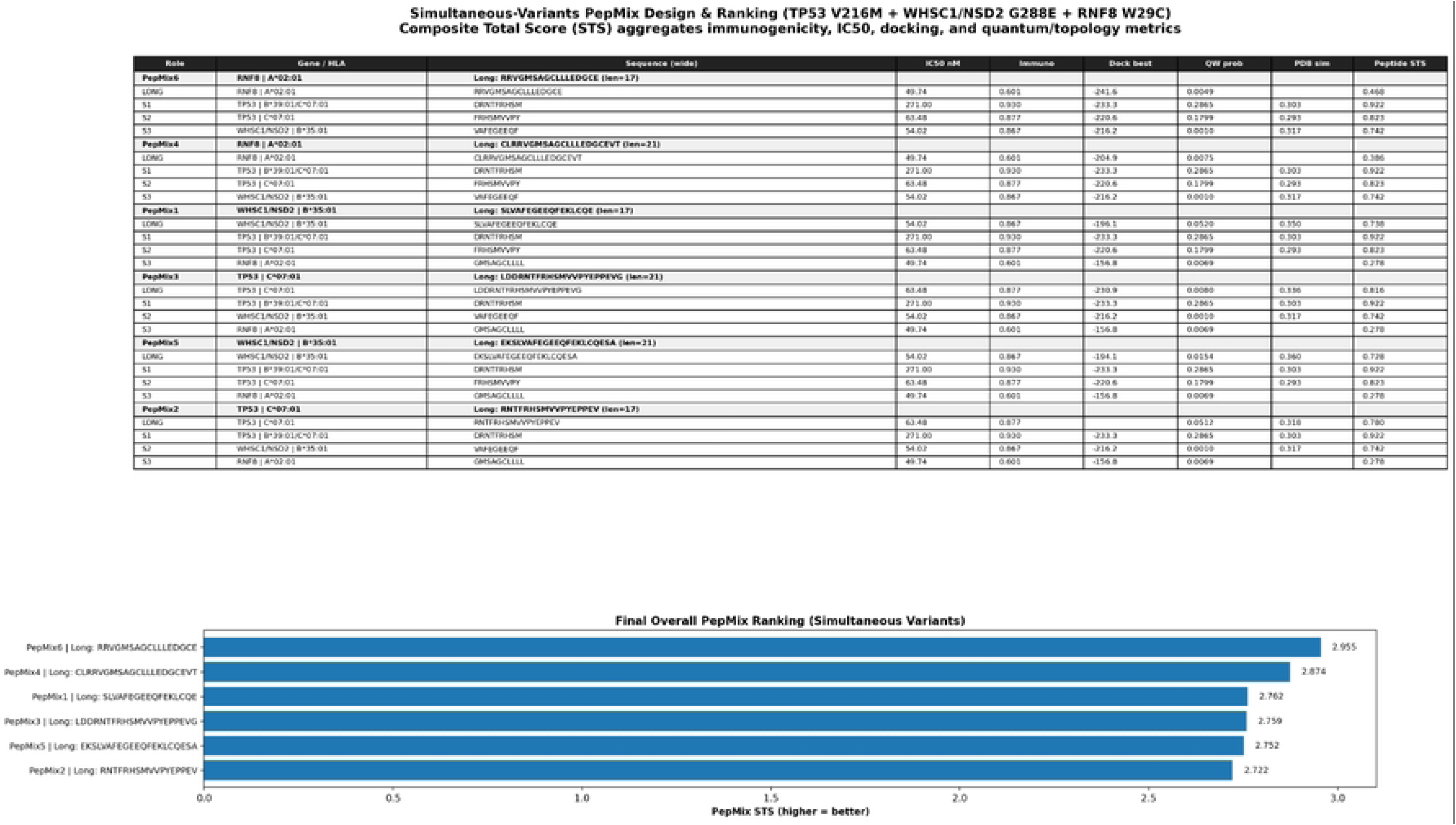

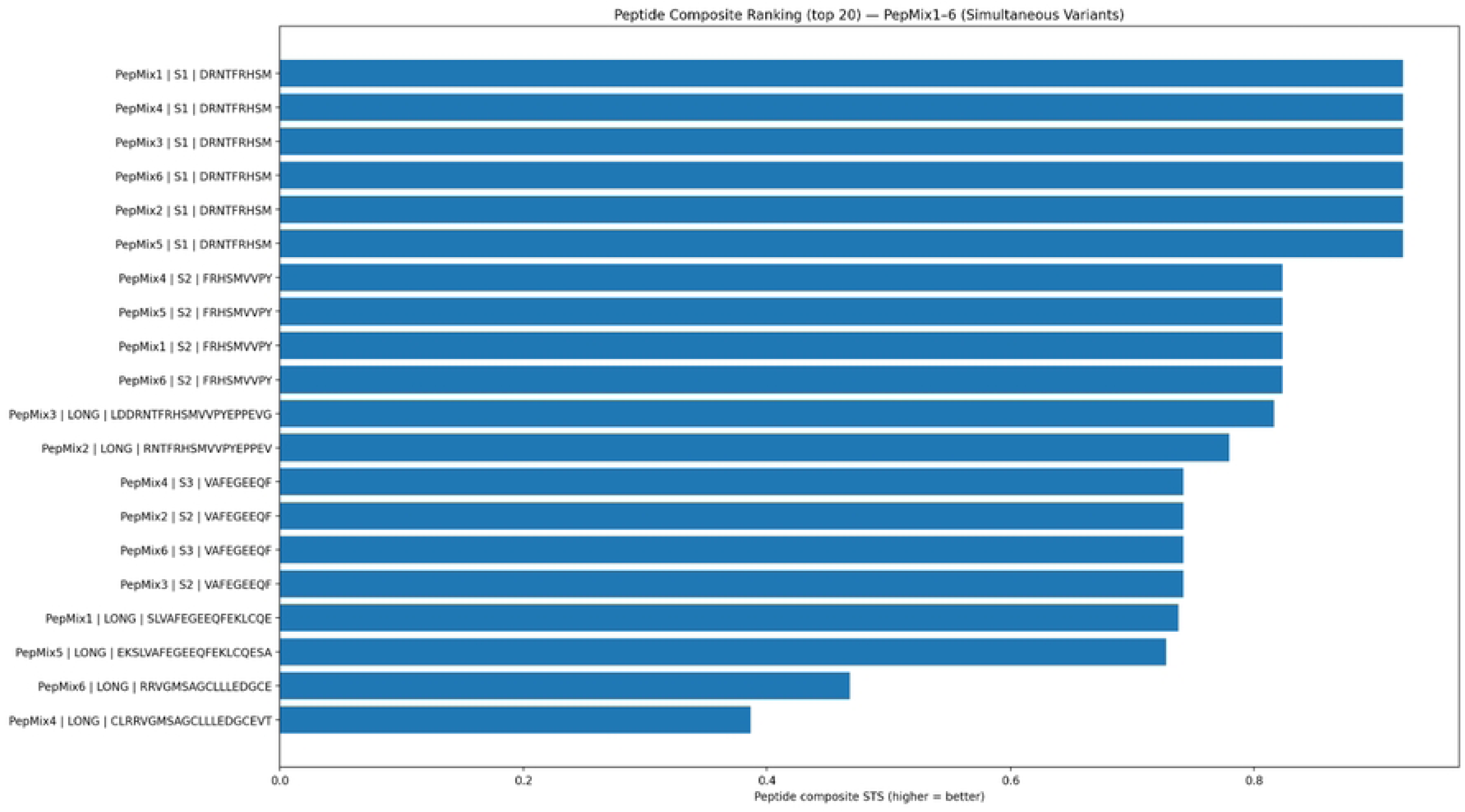

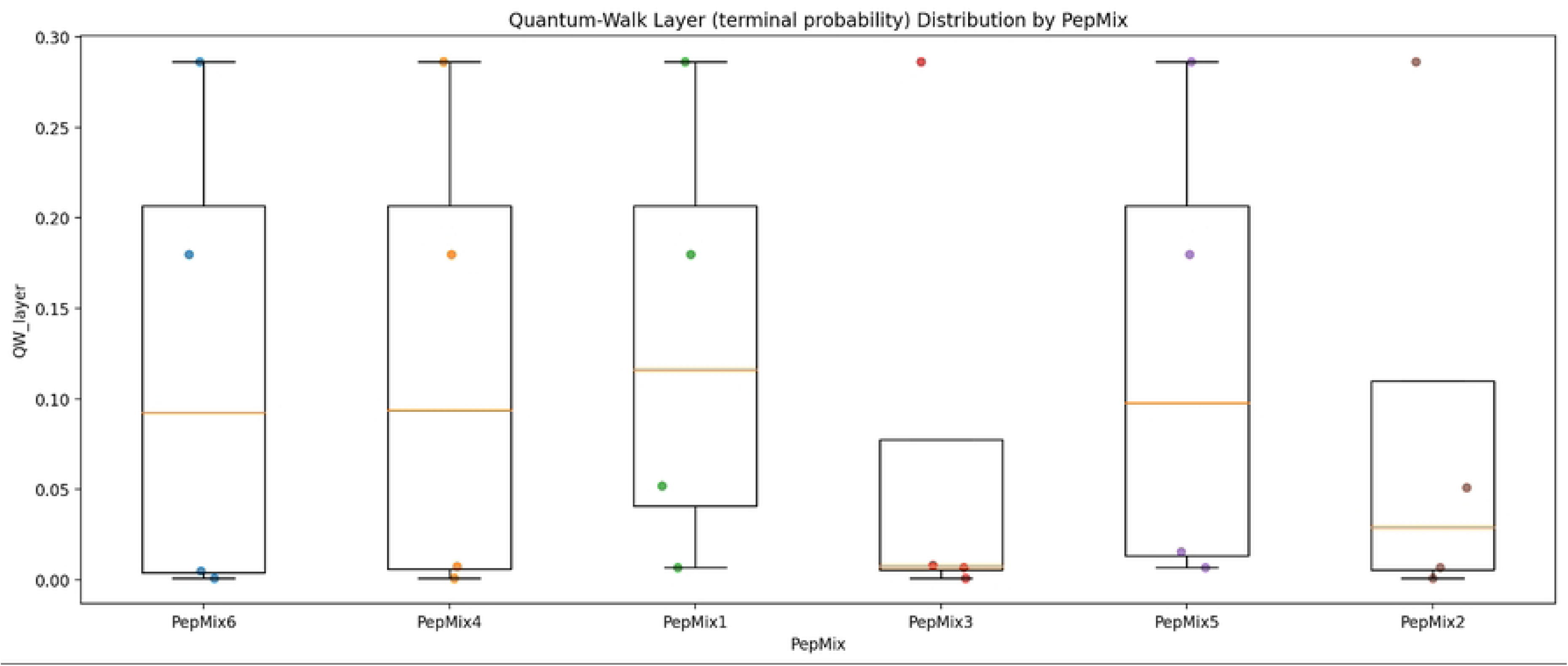

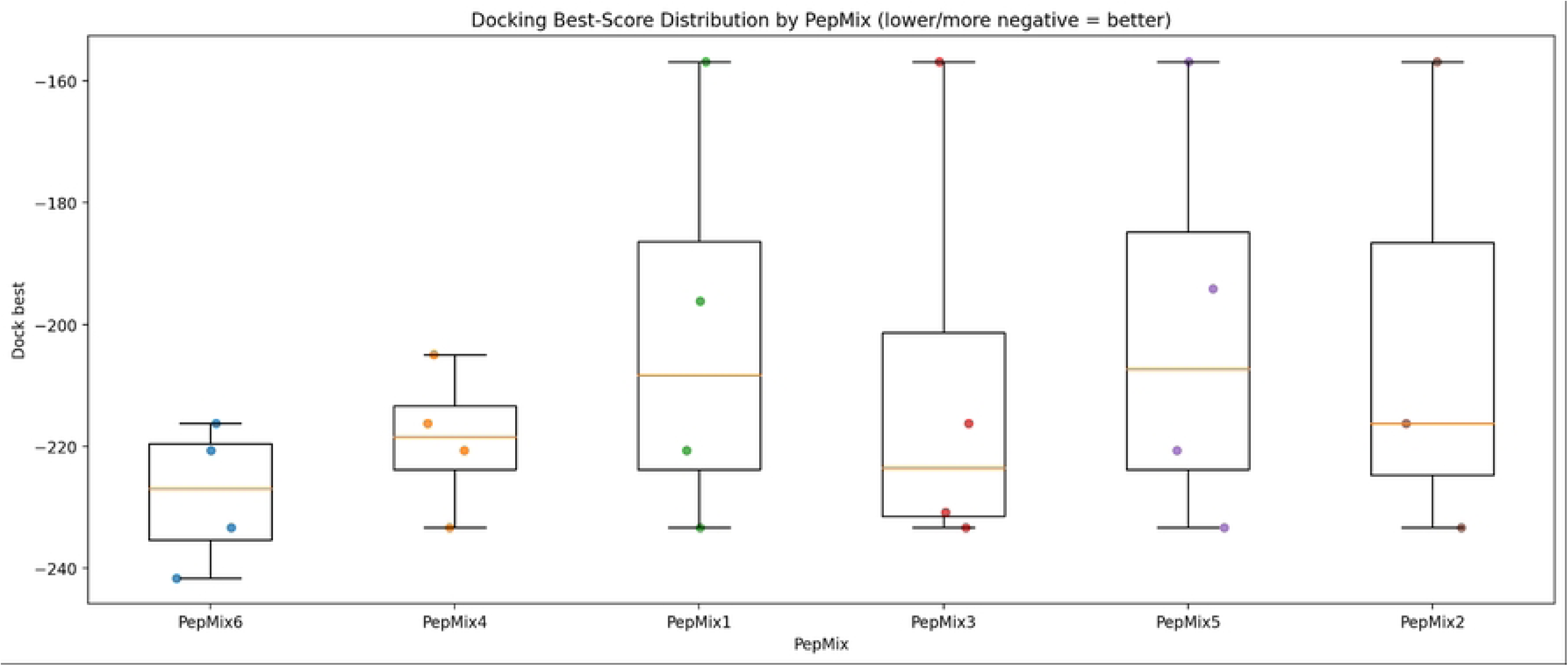

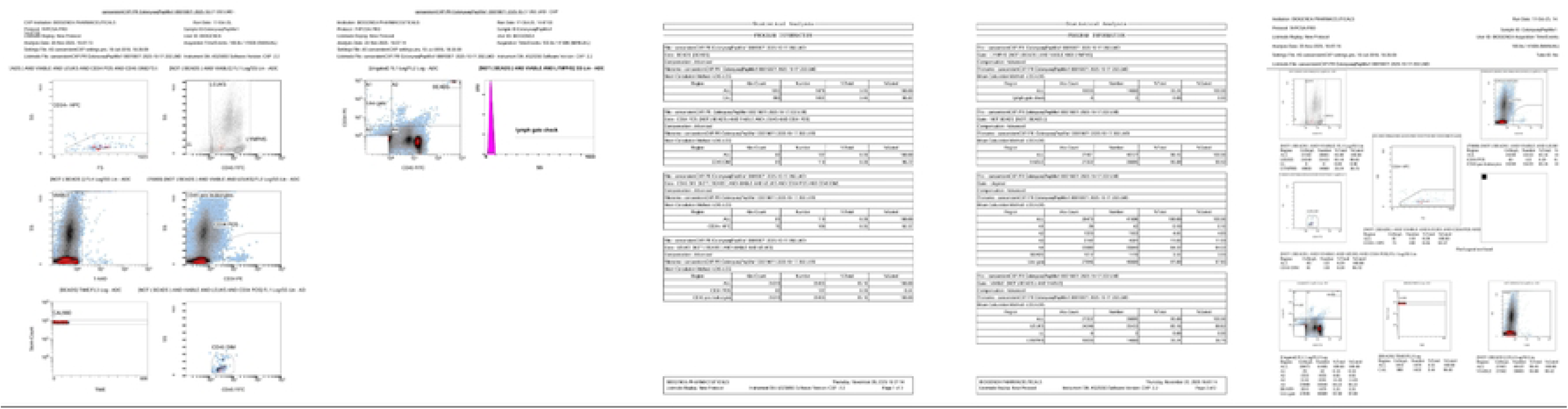

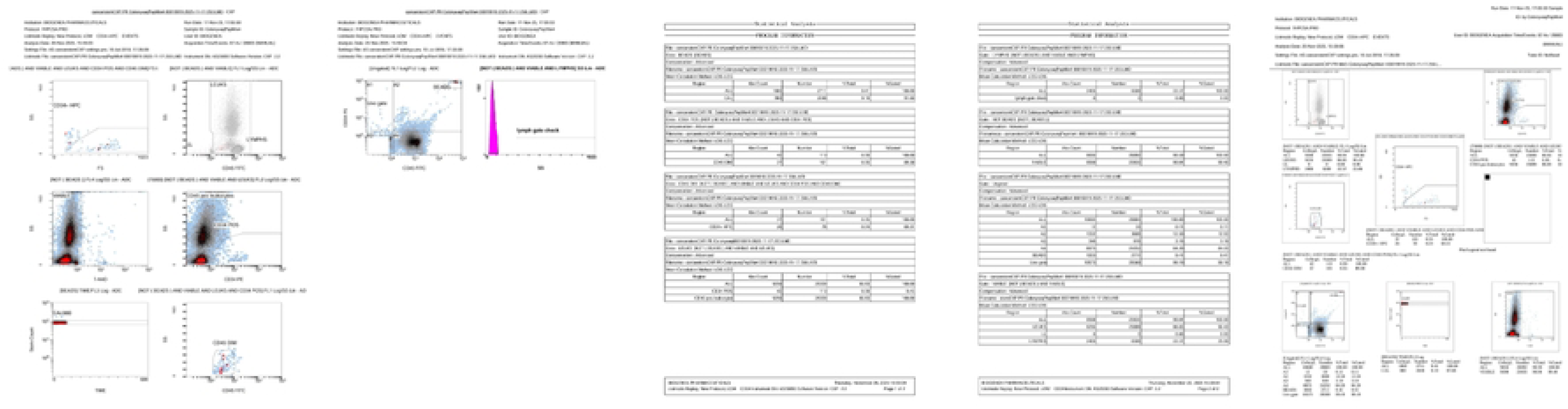

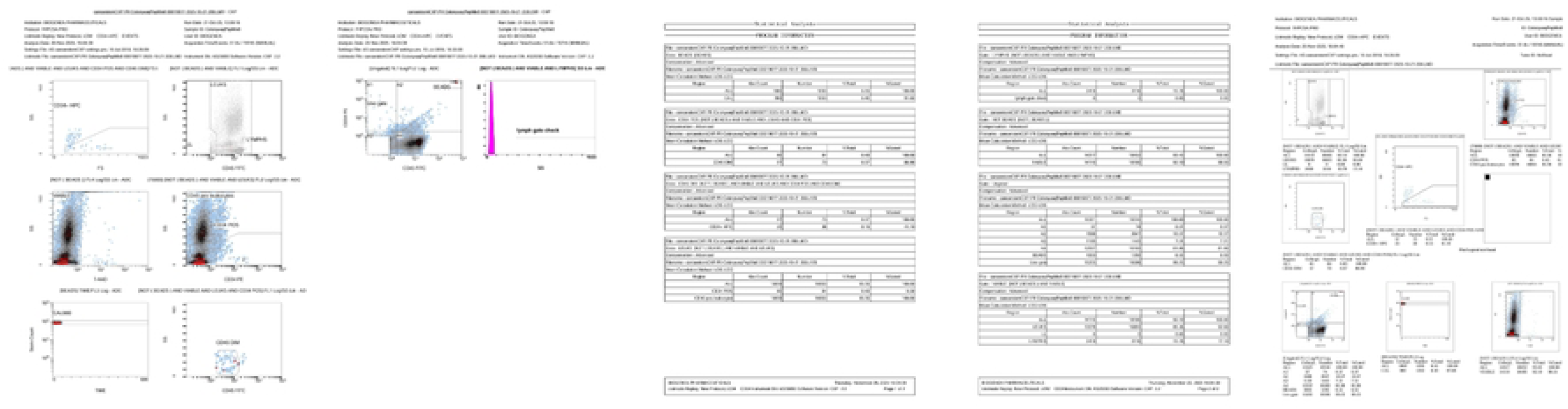

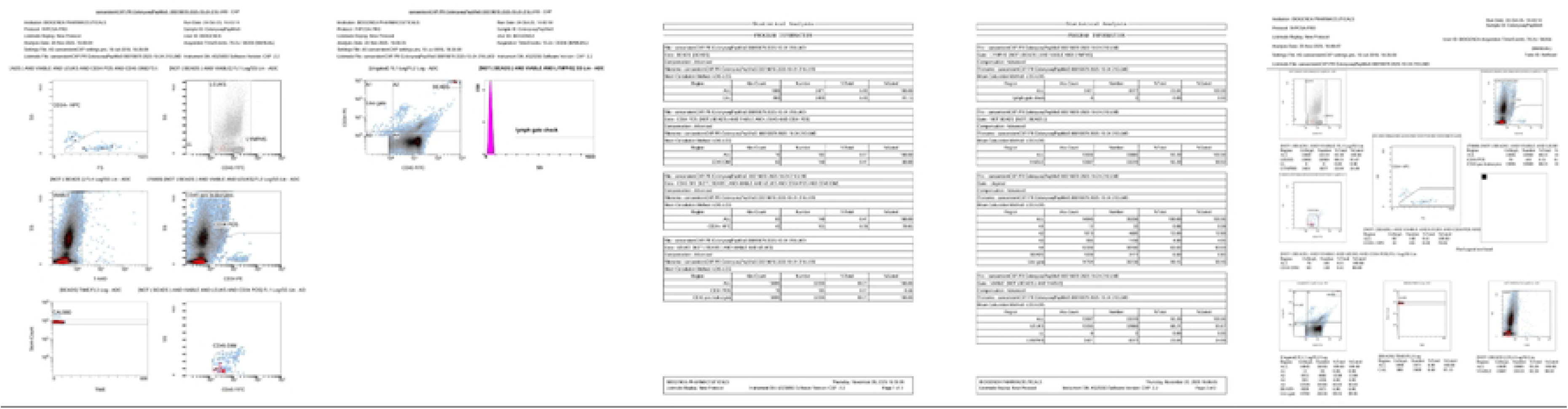

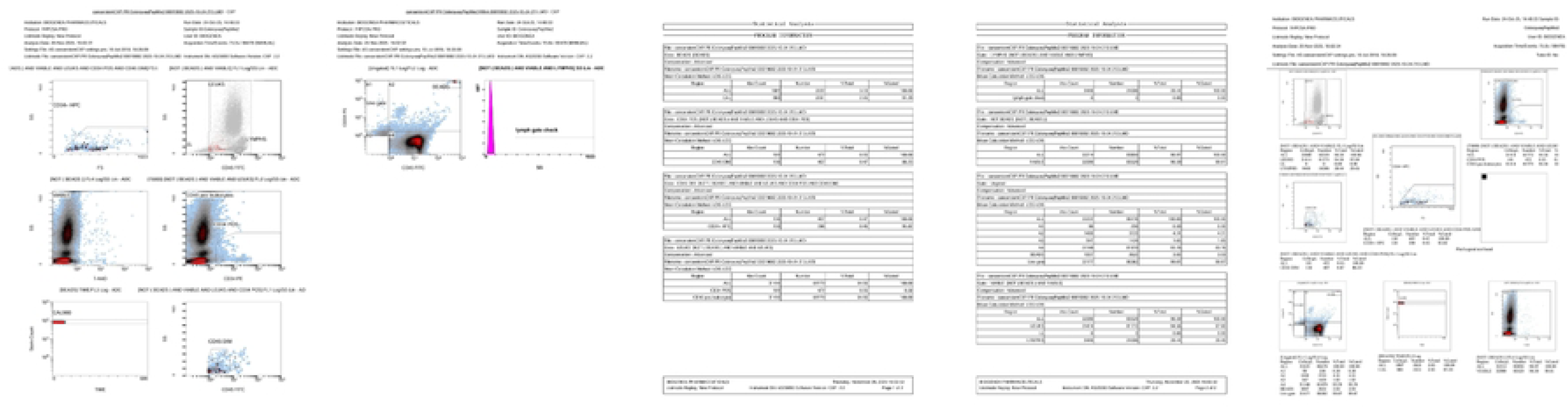

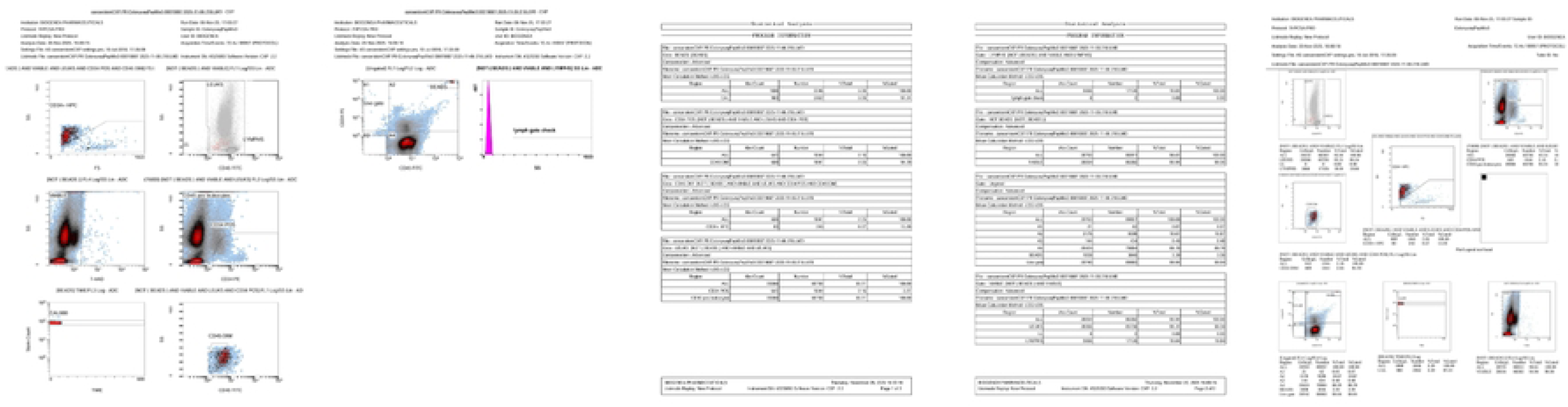

